# Complex waste stream valorisation through combined enzymatic hydrolysis and catabolic assimilation by *Pseudomonas putida*

**DOI:** 10.1101/2023.02.13.528311

**Authors:** Guadalupe Alvarez-Gonzalez, Micaela Chacόn, Adokiye Berepiki, Karl Fisher, Piya Gosalvitr, Rosa Cuéllar-Franca, Neil Dixon

## Abstract

The use of biomass and organic waste as a feedstock for the production of fuels, chemicals and materials offers great potential to support the transition to net-zero and circular economic models. However, such renewable feedstocks are often complex, highly heterogeneous, and subject to geographical and seasonal variability, creating supply-chain inconsistency that impedes adoption. Towards addressing these challenges, the development of engineered microorganisms equipped with the ability to flexibly utilise complex, heterogenous substrate compositions for growth and bio-production would be greatly enabling. Here we show through careful strain selection and metabolic engineering, that Pseudomonas putida can be employed to permit efficient co-utilisation of highly heterogeneous substrate compositions derived from hydrolysed mixed municipal-like waste fractions, with remarkable resilience to compositional variability. To further illustrate this, one pot enzymatic pre-treatments of the five most abundant, hydrolytically labile, mixed waste feedstocks was performed – including food, plastic, organic, paper and cardboard, and textiles – for growth and synthesis of exemplar bio-products by engineered P. putida. Finally, prospective life cycle assessment and life cycle costing illustrated the climate change and economic advantage, respectively, of using the waste-derived feedstock for biomanufacturing compared to conventional waste treatment options. This work demonstrates the potential for expanding the treatment strategies for mixed municipal waste to include engineered microbial bio-production platforms that can accommodate variability in feedstock inputs to synthesise a range of chemical and material outputs.

## 1. Introduction

Global supply chains are dominated by linear, carbon positive, economic models; extract, use, dispose, which create unsustainable systems reliant on finite geological reserves. Growing awareness of the damaging impact of this model upon the climate has spurred urgent action by the international community to find solutions to mitigate greenhouse gas emissions^1–6^. Biotechnology provides opportunities to shift from a linear to a more circular bio-based economic model. Towards this, the concept of biorefinery emerged as an analogue to the petroleum-based refinery industry, where fuel, power, chemicals and materials can be derived from renewable biomass, to afford both economically and environmentally sustainable outcomes^7^. While first generation biomass feedstocks provide carbon circularity, there are concerns around the diversion and expansion of arable land use towards agro-industrial production^8^. In light of projected population growth and rising sea levels, land use will become more crucial, and its dedicated allocation to feedstocks for biorefineries less sustainable^9,10^. Indeed, focus has now shifted towards the use of organic process residuals and wastes from the agricultural, forestry, paper and pulp and food industries as biorefinery inputs for sustainable production of fuels, chemicals and materials ^11–14^. However, overcoming the inherent compositional complexity and inconsistency of these feedstocks to achieve efficient conversion into bio-products is imperative for process economics. Towards this, the development of improved, more flexible, feedstock pre-treatment and microbial host strains is ongoing.

The compositional heterogeneity and variability of waste feedstocks across time (season) and space (geographic location) poses a significant biorefinery challenge, hindering their integration into current biomanufacturing processes. As such, all aspects of a platform - both in the upstream pre-treatment and downstream fermentation stages - need to be designed to tolerate a wide envelope of feedstock characteristics. With regards to the latter, when faced with a multitude of available carbon sources derived from a heterogenous feedstock, most microbes will consume them selectively due to incompatible metabolism, or sequentially with long diauxic lag times between preferred substrates. These phenotypes can prolong cultivation time and reduce the efficiency of carbon conversion – essential aspects of a cost-effective and resilient production platform^15^. Whilst huge advances have been made to engineer bacterial stains to utilise non-native substrates for growth and bio-production, to date, attempts to consolidate these metabolic capabilities into a single *super host* are still nascent. *Pseudomonas putida* is a soil dwelling bacterium that displays high solvent tolerance and is considered to be a robust host for bioproduction and bioremediation applications^16,17^. It possesses remarkably flexible non-archetypal metabolism that can accept divergent carbon substrates with no overflow metabolism and rapid shift/adaptation to nutrient availability without a long lag-phase^18^. This catabolic activity paired with facile genetic tractability make *P. putida* an ideal host for assimilating heterogeneous feedstocks. Previous reports have shown that engineered strains of *P. putida* are capable of co-utilisation of up to five substrates - sugars, aromatics, and acids - derived from pre-treated lignocellulose^19^, and more recently, substrates derived from metal-catalysed autoxidation of mixed plastics^20^. These pivotal works have enabled bioproduction of numerous valuable products, however the carbon substrates used have predominantly comprised of single carbohydrates, however these studies have predominantly utilised single feedstocks e.g., lignocellulose or plastics. This provokes the question as to whether *P. putida* could be a model host to overcome the inherent complexity and variability of mixed and complex waste feedstocks for bioproduction.

As an extreme example of a complex and variable waste feedstock, municipal solid waste (MSW) represents a highly heterogeneous mix of biogenic and non-biogenic household, commercial and industrial waste collected by a local municipality. The composition of MSW can vary considerably depending on climate, season, culture, and income level, reflecting different global patterns of consumption. Currently, global MSW generation is estimated to exceed 2 billion tonnes per year, and increase to 3.4 billion tonnes per year by 2050^21,22^. Moreover, with dumps and landfills across the globe contributing 1.6 billion tonnes of carbon dioxide emission equivalents per year to the atmosphere, improved waste mitigation and treatment approaches are needed ^22,23^. Depending on the region, between 60-85% of mixed MSW is carbonaceous, made up of food waste, garden waste, paper and board, plastic, textile, leather and rubber^24^. Current strategies for MSW utilisation/valorisation depend on the nature of the waste. Organic fraction municipal solid waste (OFMSW), such as food waste, can be composted or fed into anaerobic digestion (AD) where the outputs are fertilizer and biogas; while non-biogenic fractions can undergo incineration or pyrolysis to generate energy. These valorisation strategies, however, require strict constraints on waste-feedstock composition^23,25^, while the end-products are low value and generally fixed. Beyond these established treatment strategies, the high carbon content in mixed MSW offers enormous potential for carbon recovery and valorisation through aerobic microbial processing.

In this work, by exploiting native and engineered metabolism of *P. putida*, we demonstrate co-utilisation of common monomeric substrates derived from mixed hydrolysis of food waste, paper and cardboard pulp, mixed ester-based plastic, garden waste, and blended textile waste, namely: glucose, xylose, ferulic acid, coumaric acid, C_12_-C_18_ fatty acids, glycerol, terephthalic acid (TPA), ethylene glycol (EG), and lactic acid (**Fig. 1**). Further, we illustrate the flexibility and resilience of this platform to accept divergent, enzymatically pre-treated waste compositions as an input, and to produce distinct value-added products as an output. This proof-of-concept work illustrates the versatility of microbial based valorisation of complex mixed MSW-like feedstocks, and explores the feasibility of using this compositionally variable, land-use negative, waste feedstock in a biorefinery context.

**Figure 1.**
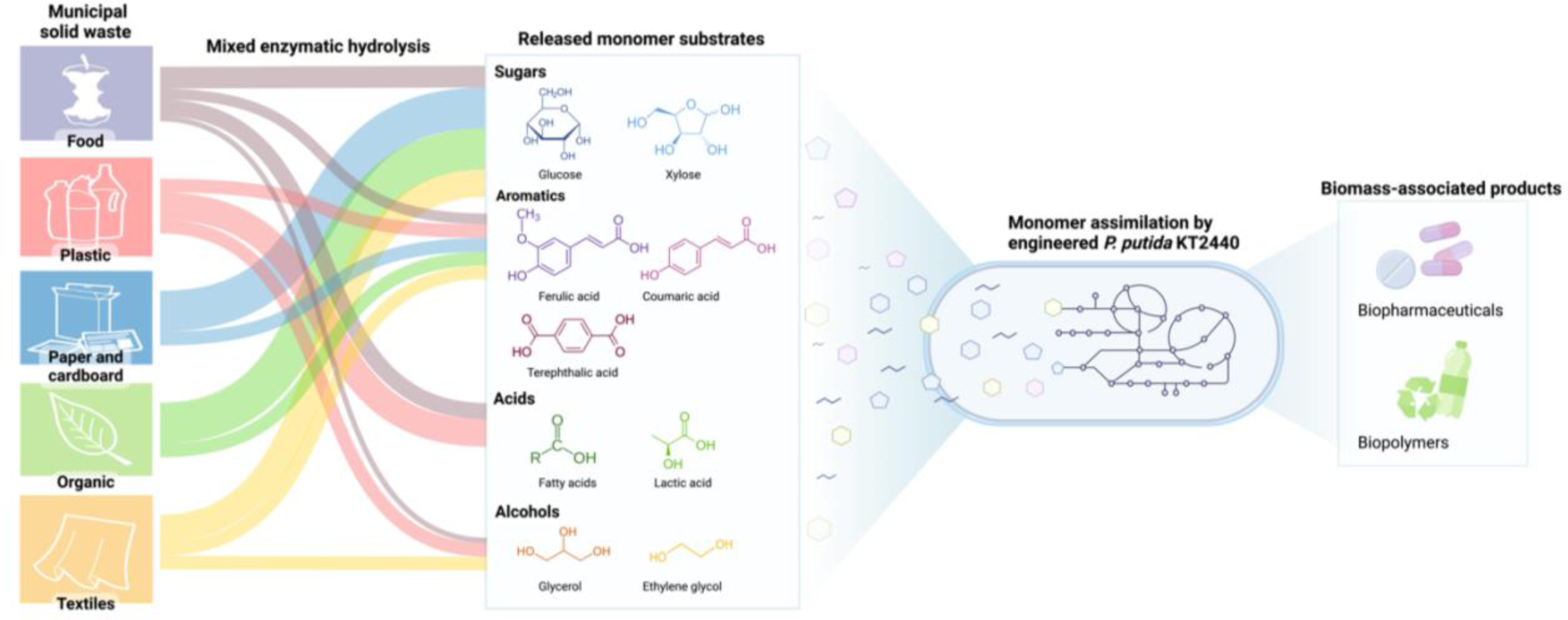
Concept for the degradation and valorisation of mixed municipal solid waste streams into valuable products. The most common hydrolytically labile carbon-based municipal solid waste fractions are enzymatically hydrolysed in a one-pot process, resulting in the release of different ratios of monomeric substrates. These monomers are subsequently co-utilised as the sole carbon and energy sources by an engineered *Pseudomonas putida* capable of synthesising biomass-associated products, including biotherapeutics and biopolymers.

## 2. Material and methods

### 2.1 Plasmid construction

All plasmid construction and sequence details can be found in Tables S1-4. All DNA synthesis and oligonucleotide primer synthesis was performed by Integrated DNA Technologies. The *XylABE* operon was amplified directly from the genome of *E. coli* BL21(DE3). DNA amplification for plasmid construction was performed using Q5 Hot Start High-Fidelity 2x Master Mix (NEB), and colony PCR was performed using Phire Green Hot Start II PCR Master Mix (Thermo Fisher Scientific). Plasmid assembly was performed using NEBuilder HiFi DNA Assembly Master Mix (NEB), followed by transformation into NEB 5-alpha competent *E. coli* cells. To validate correct plasmid assembly, Sanger sequencing was performed (Eurofins Genomic, Germany).

### 2.2 Strain construction

*P. putida* KT2440 genomic knockout and integration strains were created using the sucrose counter-selection method as described previously^26^. Briefly, electrocompetent cells were transformed with 1µg of the relevant pK18mobsacB-derived plasmid (Supplemental Table 2) and plated on LB+ Kan_20_ and incubated at 30℃. Individual colonies were streaked onto YT+25% sucrose (10 g/L yeast extract, 20 g/L tryptone, 250 g/L sucrose, 18 g/L agar) for counter selection. Individual colonies were selected and streaked once more onto YT+25% sucrose and LB+ Kan_20_ to validate genomic exchange and plasmid loss. Successful modification was further validated by cPCR and Sanger sequencing (Eurofins Genomic, Germany).

### 2.3 Expression and purification of the leaf-branch compost cutinase (LCC) mutant

The LCC (ICCG)^27^ gene was expressed in BL21(DE3) cells in a Type NLF 22, 30 L BioEngineering fermenter containing 20 L enhanced 2xYT medium (20 g/L yeast extract, 32 g/L tryptone, 17g/L NaCl, 2 g/L glucose, 17.6 mM Na_2_PO_4_ (pH 7.4)) at 37°C. Once at an OD_600_ of 0.8 was reached 0.25 mM isopropyl-β-D-1-thiogalactopyranoside (IPTG) was added and the temperature was reduced to 20°C for overnight incubation under a continuous compressed air gas flow of 30 L min^-1^. Cells were harvested by centrifugation (4 °C, 6000 *g* for 10 minutes) and cell pellets (∼100g) were resuspended in buffer A (200 mM NaCl, 50 mM Tris pH 7.5) supplemented with DNase, RNase (Sigma) and Complete EDTA-free protease inhibitor cocktail (Roche). Cells were lysed using a cell disruptor (Constant Systems Ltd.) at 20,000 psi and the lysate was clarified by centrifugation at 125,000 *g* for 90 minutes. The supernatant was applied to a Ni-NTA agarose drip column (Qiagen) and eluted with buffer A supplemented with 250 mM imidazole. Imidazole was removed from the eluted protein using a 10-DG desalting column (Bio-Rad) and purified to homogeneity with a 5 mL HiTrap HP column (GE healthcare) attached to a ÄKTA pure purification system using a linear gradient of 0-250 mM imidazole in buffer A over 35 CV. Fractions were subjected to SDS-PAGE analysis (BioRad, Mini-Protean TGX stain free precast SDS-gel with a 4-20% gradient) and those found to contain the purified protein were pooled and concentrated using a 10 kDa molecular weight cut-off Vivaspin spin column centrifugal concentrator (GE Healthcare) at 3894 x g and 4°C. Imidazole was removed using a 10-DG desalting column (Bio-Rad) and aliquoted, flash frozen in liquid nitrogen and stored at - 80 °C until required.

### 2.4 Mixed waste enzyme hydrolysis

All waste materials and enzymes used are listed in Table S5. Mixed waste hydrolysis was carried out at 25% solid loading (w/w) in 100 mM sodium phosphate buffer pH 7.0 in 500 mL baffled bottles at 50℃ and 125 rpm for 72h. The substrate composition of high- and low-income waste is described in Table S7. Mixed waste slurry was incubated at 50℃ for 1 hour prior to the addition of enzyme: 50 mg CTEC-2/g lignocellulosic substrate, 0.75 mg LCC/g PET substate, 1 mg CE1/g lignocellulosic substrate, 9 U CAL-B/g palm oil, 10 U α-amylase/g starch, 5 U maltase/g starch, and 500 µg/mL carbenicillin were added to the slurry. Separately, 0.5 mg Proteinase K/g PLA substrate and PLA was incubated at 50℃ to promote absorption before addition to the mixed waste slurry. Hydrolysate samples were taken at 8, 24, 48, and 72 hours, crudely filtered, ultracentrifuged at 40,000 g for 30 minutes, and passed through a 0.22 µm filter for fine filtration and sterilisation. Samples were taken for product quantification pre and post 0.22 µm filtration. Hydrolysates were stored at -20℃. Individual substrate hydrolyses were carried out as described above with variable solid loadings (1-10 % w/w) and 0.1% sodium azide (NaN_3_) as an antimicrobial agent instead of carbenicillin. Hydrolysis of post-consumer waste was carried out as described above at 10% (w/v) solid loading and at variable temperature and pH. Those hydrolyses carried out at pH 5.0 were done in 75 mM citrate phosphate buffer, and those at pH 9 were done in 100 mM Bis-Tris propane buffer.

### 2.5 *P. putida* growth and consumption on individual and mixed substrates

Precultures of wild type and engineered *P. putida* KT2440 were incubated in M9 (6.78 g/L Na_2_HPO_4_, 3 g/L KH_2_PO_4_, 0.5 g/L NaCl, 1 g/L NH_4_Cl, 2 mM MgSO_4_, 100 μM CaCl_2_, 1x trace elements) +/- antibiotic and 5 mM carbon source(s) at 30 ℃ and 180 rpm for 12-24h. Flasks containing 100 mL M9 +/- 500 ug/L carbenicillin and 10 mM of respective carbon source(s) were inoculated in triplicate from the preculture to an OD_600_ of 0.05 and incubated at 30 ℃ and 180 rpm. Intermittent samples were taken to determine OD_600_, cell dry weight (CDW), and substrate consumption. For the latter, 2 mL of culture was centrifuged at 4000 rpm and 4 ℃ for 10 minutes before the supernatant was filtered through a 0.22 µm filter and stored at -20 ℃. For those cultures in which free fatty acid (C12:0-C18:0) was used as a carbon source, solid substrate precipitation prevented quantification of consumption. To evaluate the free fatty acid consumption rate, linoleic acid (C18:2) – which is an oil at room temperature, was used. M9+ 10 mM linoleic acid was incubated shaking at 180 rpm for 2 hours to create an emulsion prior to inoculation with *P. putida*, and samples were taken while cultures remained shaking. CDW was determined by centrifuging 1 mL of culture at 4000 rpm for 2 minutes in a pre-weighed microcentrifuge tube, followed by discarding the culture supernatant, washing the cell pellet in 1 mL of M9 once, and incubating the cell pellet for 16 h at 50 ℃ before determining the dry pellet weight.

### 2.6 *P. putida* growth, consumption, and production on mock High-Income and Low-Income mixed waste hydrolysate

*P. putida* Δ*gcd*::*XylABE* harbouring either tphKAB or tphKAB-IFNα2a was pre-cultured as described above, followed by inoculation in triplicate into 0.9 mL of corresponding hydrolysate and incubated in 48-well flower plates in a BioLector® at 30 ℃ for 48 h (m2p-labs GmbH, Germany). For flasks cultivations, pre-cultures were inoculated in triplicate into 400 mL of M9+ 500 µg/L carbenicillin plus mock high Income hydrolysate (37 mM glucose, 5.4 mM xylose, 2.2 mM TPA, 2.2 mM ethylene glycol, 0.23 mM ferulic acid, 0.37 mM coumaric acid, 1 mM lactic acid, 1.7 mM glycerol, 0.1 mM C12:0, 0.15 mM C14:0, 2.3 mM C16:0, 0.25 mM C18:0, 1.8 mM C18:1; 0.5 mM C18:2) or low Income hydrolysate (30 mM glucose, 4 mM xylose, 1 mM TPA, 1 mM ethylene glycol, 0.5 mM ferulic acid, 0.73 mM coumaric acid, 0.5 mM lactic acid, 3 mM glycerol, 0.18 mM C12:0, 0.27 mM C14:0, 4 mM C16:0, 0.45 mM C18:0, 3.15 mM C18:1; 0.9 mM C18:2) to an OD_600_ of 0.05 and incubated at 30 ℃ and 180 rpm. Intermittent samples were taken to determine OD_600_, CDW, substrate consumption and product formation as described above, with the following exceptions: (i) no free fatty acid quantification was performed at they initially formed a solid precipitate, (ii) at intermittent time points 5 mL of culture pellet was harvested and stored at -20 ℃ for product quantification.

### 2.7 Analytics

#### 2.7.1 Quantification of Glucose, Xylose, Glycerol and Lactic acid by HPLC

All chemicals and reagents are listed in Table S5. 5 µL of neat 0.22 µm filtered culture supernatant or enzyme hydrolysate containing monomeric sugar (D-glucose and D-xylose), glycerol and/or L-lactic acid was injected and separated by a Supelcogel™ C–610H (6% Crosslinked) column operating at 30 ℃ with an isocratic 0.1% (v/v) H_3_PO_4_ mobile phase run at a flow rate of 0.5 mL/min. Sugars and glycerol were quantified from the RID, and lactic acid was quantified from the UV at 210 nm. Product identification and quantification was carried out using authentic standards.

#### 2.7.2 Quantification of TPA, Ferulic acid, Coumaric acid, and relevant metabolic acid intermediates by HPLC

300 µL of 0.22 µm filtered culture supernatant or enzyme hydrolysate was appropriately diluted and combined with 300 µl methanol and 13 µl of 3M HCl prior to injection of 10 µL onto a Agilent 1200 Infinity Series HPLC with a Kinetex 5 µm C18 100 Å column, 250 x 4.6 mM column (Phenomenex^®^) operating at 25 ℃ with a flow rate of 0.8 mL/min and a linear gradient of 5% acetonitrile:95% H_2_O to 30% acetonitrile:70% H_2_O over 20 minutes with a 4 minutes post run hold at 5% acetonitrile:95% H_2_O. All solvent contained 0.1 % TFA. All acids were quantified from the UV at 260 nm. Product identification and quantification was carried out using authentic standards.

#### 2.7.3 Quantification of Ethylene glycol by HPLC

300 µL of 0.22 µm filtered culture supernatant or enzyme hydrolysate was appropriately diluted and combined with 350 µL of 40% NaOH (w/v) and mixed thoroughly before the addition of 10 µL benzoyl chloride and incubation at 40 ℃ and 4000 rpm for 20 minutes. Samples were extracted into 1 mL of hexane, of which 500 µL was put into a 1.5 mL glass vial and dried fully under nitrogen before resuspension in 1 mL of H_2_O. 10 µL of derivatized sample was injected onto a Agilent 1200 Infinity Series HPLC with a Kinetex 5 µm C18 100 Å column, 250 x 4.6 mM column (Phenomenex^®^) operating at 25 ℃ with an isocratic 55% acetonitrile (0.1% TFA): 45% H_2_O (0.1% TFA) mobile phase at a flow rate of 1 mL/min. Derivatized ethylene glycol was quantified from the UV at 230 nm. Product identification and quantification was carried out using authentic standards.

#### 2.7.4 Quantification of free fatty acid by GC-MS

500 µL of 0.22 µm filtered culture supernatant or enzyme hydrolysate spiked with 10 µg of pentadecanoic acid (C15:0) as an internal standard before being extracted with 1 mL of 2:1 chloroform/toluene. The organic phase was washed with 1 mL of 0.9% NaCl (w/v) before being dried down under nitrogen gas. Dried sample was resuspended in 3 mL of 1 M MeOH-HCl and incubated at 50 ℃ for 60 minutes in a sealed vessel. Once cooled to room temperature, 0.9% NaCl (w/v) was added before extraction with a variable volume of hexane. Fatty acid methyl esters (FAMEs) were quantified using an Agilent Technologies 5975 MSD coupled to a 7890B GC with a PAL RSI 85 autosampler and an Agilent VT-5HP column (30 m x 0.25 mM x 0.1 µM). Injector temperature was set to 240 ℃ with a split ratio of 1:100 (1 µL injection). Column temperature was held at 140 ℃ for 5 minutes before ramping at 10 ℃/min to 300 ℃ and being held for 7 min. Ion source temperature of the mass spectrometer (MS) was set to 230 ℃ and spectra were recorded from m/z 50 to m/z 550. Compound identification was carried out using authentic standards and comparison to the reference spectra in the NIST library.

#### 2.7.5 Quantification of PHA monomers by GC-MS

At discrete timepoints during the time course of PHA production by a relevant strain of *P. putida*, 2 mL of culture was centrifuged at 4000 rpm for 10 minutes at 4 ℃, the culture supernatant was discarded, and the cell pellet was washed twice in 5 mL of 0.9% NaCl (w/v) before being lyophilized. Methanolysis was then carried out by resuspending the lyophilized cell pellet in 3 mL of 1 M MeOH-HCl with 25 µg of benzoic acid as an internal standard and incubating at 100 ℃ for 150 min in a sealed vessel. Once cooled to room temperature, samples were washed with 0.9% NaCl (w/v) before being extracted with 2 mL of hexane. Quantification of hydroxyacid methyl esters (HAMEs) was carried out by GC-MS as described in the section above with the following exceptions: split ratio of 1:10, column temperature was initially held at 50 ℃ or 5 minutes before ramping at 10 ℃/min to 280 ℃ at being held for 5 min. Compound identification was carried out using authentic standards and comparison to reference spectra in the NIST library.

### 2.8 Western blot and quantification of heterologous IFNα2 production

At either a final time point or discrete timepoints throughout a growth course with the relevant strain of *P. putida*, 5 mL of culture was harvested, centrifuged, and the cell pellet was resuspended in lysing buffer (1mL benzonase nuclease, 1 tablet protease inhibitor cocktail in 50ml 1xPBS) before sonication. Samples for comparison of protein production by *P. putida* tphKAB to strain tphKAB-EG12 were resuspended to an according volume for normalisation to OD_600_ = 10. Samples for IFNα2a quantification were resuspended to a normalising volume to OD_600_ = 20. Sonicated samples were resuspended in SDS-PAGE loading buffer (ThermoFisher), supplemented with 75mM dithiothreitol and heated at 95 ℃ for 10 minutes. Equal amounts of whole cell protein were separated on a 15% SDS-polyacrylamide gel (Bio-Rad) by SDS-PAGE. The SDS gel was then transferred to a nitrocellulose membrane and blocked with 5% skimmed milk in PBS. Membranes were incubated overnight (50 rpm, 4°C) with either mouse anti-His monoclonal (Thermo Fisher; 1:3000 in 5% skimmed milk) or anti-IFNa2 primary antibody (abcam, 193055, 1:2000 in 5% skimmed milk). Membranes were washed with PBS, followed by incubation with an IRDye anti-mouse or anti-rabbit IgG secondary antibody (LI-COR; 1:30000 in 5% skimmed milk). IFNα2a samples were quantified based on densitometry analysis using Li-Cor Image Studio 5.0 against pure IFNα2a protein standards of known concentrations prepared in the same way.

### 2.9 Data analyses, model construction, and simulation

The most recent genome-scale metabolic model of *P. putida* KT2440, iJN1463^28^ was extended to incorporate modifications for TPA and xylose metabolism. For EG, the glyoxylate carboligase reaction, GLXCL, was eliminated to simulate operon repression. The final model was subjected to flux balance analysis (FBA) using the COBRA toolbox ^29^ with the linear programming solver Gurobi (www.gurobi.com) in MATLAB 2020a (Mathworks Inc.). To predict flux variation from the equimolar experiment, the model was constrained with uptake rates obtained experimentally and the biomass reaction was set as the objective function. ATP maintenance requirements were fixed at 0.92 mmol gCDW^-1^ h^-1^. A prediction of the total CO_2_ produced was calculated by multiplying the resulting CO_2_ production rate against the total growth time for the specified period.

Statistical significance or correlation analysis were performed by with Prism 9 (GraphPad Software, Inc). All experiments were performed in at least three biological triplicates. Manual linear regression analysis was performed during exponential growth phase to calculate growth rate (μ) and specific uptake rate on the corresponding substrate(s) (*q*_s_). The biomass yield when using mixed substrates (Y_S/X_) was calculated as a lumped value. The total amount of carbon (C-mol) for biomass was based on the elemental carbon composition of *P. putida* KT2440^30^, calculated to be 24.5 g/mol.

### 2.10 Life cycle assessment and Life cycle costing

This prospective assessment estimates the climate change (CC) impact and life cycle costs of the valorisation of MSW to PHA via the scaled-up proposed bioprocess, and to compare with three conventional treatment methods - incineration with energy recovery, landfill and open burning. Life cycle assessment (LCA) and Life cycle costing (LCC) are the methods of choice to perform such analysis ^31–33^, and we applied consistently. The scope of the study is cradle to gate and includes i) an enzymatic hydrolysis of MSW for both high- and low-income countries compositions, ii) microbial fermentation and PHA bioproduction on the MSW-derived substrates, and iii) a PHA extraction step. The MSW feedstock is assumed burden free, and the system is credited with the production of PHA from corn^34^. The functional unit of the study is defined as “the treatment and valorisation of 1 tonne of MSW”. The primary data (raw materials, energy consumption, waste and product yield) was obtained directly from the experiments carried out in this study, combined with data from literature for the PHA extraction step^35^ and other relevant sources for scaling up^35,36^, whilst secondary data was obtained from the ecoinvent database v3.8^37^. The waste generated throughout the process is assumed to be treated either by incineration or sent to wastewater treatment. A full description of the inventory data and assumptions for both LCA and LCC studies is included in **Tables S8-10**. Whilst the LCA study excluded the plant infrastructure due to negligible contributions^38^, the LCC work considered the costs of major equipment required (**Table S11**). All costs are based on 2023 prices and were estimated based on literature, market prices and industry reports as detailed in the Supporting Information. The LCA modelling was carried out in LCA for Experts Software v9.1^39^ and the mid-point CC impact was estimated using the ReCiPe method ^40^. The environmental impacts and LCC values of the conventional treatments were sourced from the ecoinvent database v3.8^37^ and the study by Slorach, et al.^41^, respectively using an established LCC methodology^33^.

The LCC has been estimated according to the following equation:

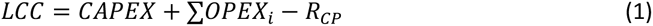

where:

LCC Life cycle costs of the valorisation of MSW to PHA via proposed bioprocess (£/t MSW)
CAPEX Capital expenditure including cost of equipment, service facilities costs, and maintenance costs (£/t MSW)
OPEX_i_ Operating expenses including costs of raw materials, utilities consumption, waste management costs (£/t MSW) of a given stage in process *i*
R_CP_ Revenue from PHA produced (£/t MSW).

The cost of equipment has been estimated based on literature (see **S11**). The capacity of the equipment reported in the relevant literature has been scaled to match the proposed capacity in this study for the production of 1 tonne of PHA using the equation below:

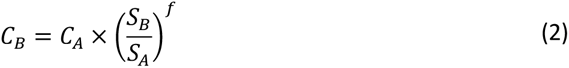

C_B_ Cost of larger equipment (£)
C_A_ Cost of smaller equipment (£)
S_A_ Capacity of smaller equipment (m^3^)
S_B_ Capacity of larger equipment (m^3^)

f Scaling factors: 0.78 for storage tank, hydrolysis tank, and extraction reactor; 0.75 for fermenter, 0.6 for crusher, mixer, filter, centrifuge, dryer, and evaporator.

The resulting equipment costs are provided in **Table S11**. The capital expenditure has been updated using the Chemical Engineering Plant Cost Index for 2023 ^42^.

## 3. Results and Discussion

### 3.1 Enzymatic hydrolysis of mixed municipal solid waste-like feedstock

Investigation into the enzymatic pre-treatment of mixed MSW began by selecting an assortment of commercial and post-consumer materials representative of the five most prevalent MSW fractions that can be amenable to enzymatic hydrolysis (**Table S6**). Post-consumer municipal paper pulp was used to directly exemplify the paper and cardboard fraction. Wheat straw served as a lignocellulosic example of organic waste. For plastic waste, we selected an amorphous PET film and poly lactic acid (PLA) beads. For food waste, we utilised palm oil, corn cob, barley, apple, and beetroot. Finally, a cotton-polyester blended fabric was used to represent textile waste. Due to varied waste management practices, the composition of municipal waste varies greatly across geographical zones^22^. A region’s GDP can accurately reflect its consumption practices, ultimately leading to distinctive waste compositions. As such, we replicated two waste scenarios using the selected feedstocks to represent (i) high and upper-middle income countries (high) and (ii) lower-middle- and low-income countries (low) (**Fig. 2a**). In the resulting formulations, paper and cardboard constituted the predominant fraction in the high-income representation (34%), followed by food (22%), garden (21%) and plastic (18%) waste. Low-income countries comprised predominantly of food and garden waste (38% each), with only 9% of both plastics and paper and cardboard waste. Textiles contributed only a small fraction of the waste for both high- and low-income countries (5% and 6%, respectively) (**Fig. 2a**).

**Figure 2.**
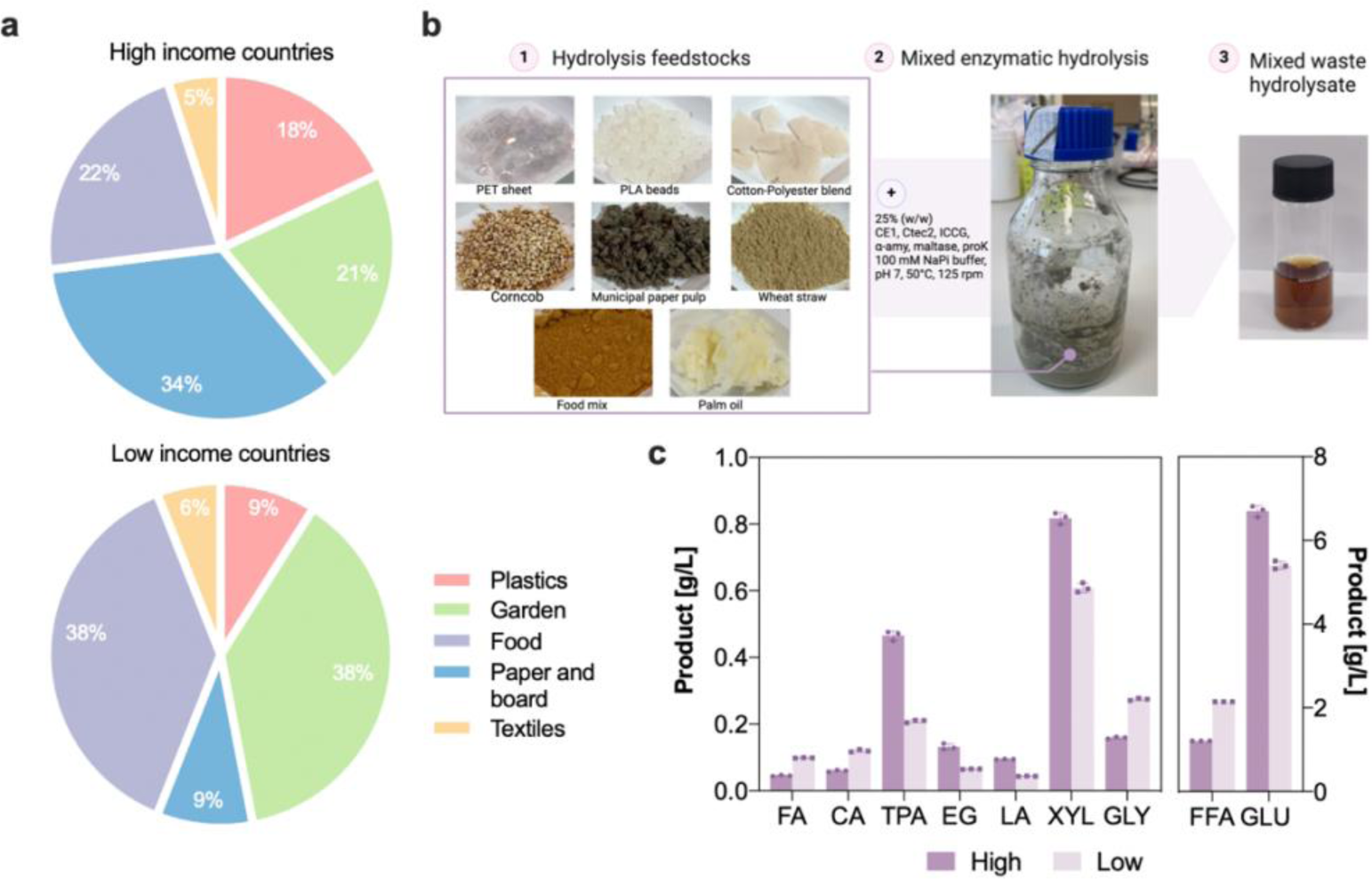
Enzymatic hydrolysis of different mixed MSW representations. (**a**) Adjusted waste composition from high- and upper-middle income countries (up) and lower-middle- and low-income countries (down). (**b**). Enzymatic hydrolysis of both high and low-income waste compositions. The selected waste fractions are prepared for hydrolysis according to proportions in (a) at a 25% solid loading (w/w) and enzymatically hydrolysed at 50°C for 72 hours (2). (**c**) Microbioreactor growth assays with tphKAB-XylABE in both waste hydrolysates and waste-derived (synthetic) substrates. For **c** and **d**, all data points are mean ± SD of n = 3 biological replicates.

To minimise complexity, cost, and time, we established a common enzymatic pre-treatment condition that would strike a compromise between enzymes with disparate environmental optima and support the mixed enzymatic hydrolysis of these feedstocks in one pot (**Fig. 2**). The two representative feedstocks were prepared to the relevant ratios for the high and low-income formulas (as illustrated in **Table S6**) to a final solid loading of 25% (w/w), and were treated with a tailored cocktail of degradative enzymes (results for individual feedstock hydrolysis can be found in **Fig. S1-S8**). We then quantified a selection of the most abundant monomers that were expected to be released from the selected polymers, with both mixed waste hydrolyses affording release of the corresponding sugars, acids, alcohols, and aromatic monomers (**Fig. 2b & Fig. S9**). The major hydrolysis product obtained from both mixed feedstock compositions was glucose. Fatty acids were the next most abundant, with a mixture of C_12:0_, C_14:0_, C_16:0_, C_18:0_, C_18:1_ and C_18:2_ acids being recovered (**Fig. S4**). Here the low-income formula yielded a higher fatty acid titre, concordant with the higher food fraction in this composition. The lowest yields were obtained for lactic acid, ferulic and coumaric acid. A total of 9.7 g L^-1^ and 9.0 g L^-1^ of combined products were obtained for the high and low-income feedstocks, respectively, corresponding to 0.365 and 0.405 mol of total carbon (C-mol) (**Fig. 2c**).

While the proposed enzymatic pre-treatment provides abundant carbon for subsequent bioproduction exploration, it’s important to note that the illustrative feedstocks are limited to those compatible with enzymatic hydrolysis – with the exclusion of a protein fraction as the required proteinase concentration would compromise the efficacy of the other enzymes within the cocktail. We conducted comparative analyses to benchmark the one-pot mixed-fraction hydrolysis against the hydrolysis of single stream synthetic and true post-consumer MSW waste fractions (**Fig. S10**). In most cases, under optimal conditions, single stream hydrolysis resulted in significantly improved monomer release compared to the one-pot process conducted under ‘common’ hydrolysis conditions (**Fig. 2b**). In addition, hydrolysis of post-consumer paper pulp yielded substantially more glucose and xylose, while hydrolysis of a post-consumer PET bottle resulted in lower TPA release compared to low-crystalline PET film (**Fig. S3, S5, and S10**). These results indicate the need to improve this one-pot enzyme pre-treatment to address diverse enzyme requirements and variable feedstock recalcitrance in order to enhance efficiency in future applications.

### 3.2 Strain engineering of *Pseudomonas putida* KT2440 to enable assimilation of MSW-like derived carbon substrates

*P. putida* KT2440 can natively metabolize a wide variety of waste-derived substrates including aromatics, acids, alcohols, and some sugars^17,43,44^. However, this strain cannot efficiently utilise some of the compounds that were recovered in high proportion from the enzyme pre-treatments, namely TPA, EG and xylose. Thus this necessitated genetic engineering of *P. putida* to incorporate the corresponding catabolic pathways.

#### 3.2.1 Establishing dynamic TPA metabolism in *Pseudomonas putida* KT2440

As a first step towards engineering a robust microbial strain capable of co-assimilating all the main waste monomeric substrates, we integrated a terephthalate (TPA) utilisation pathway into *P. putida* KT2440 – the aromatic monomer of polyethylene terephthalate (PET) plastic. To achieve this, we developed a genetic construct expressing a TPA *tph* catabolic operon (*tphA2_II_A3_II_B_II_A1_I_*_I_) from *Comamonas* sp. E6^45,46^ and a TPA transporter, *tphK,* from *P. mandelli* JR1^47^. A dynamic genetic feedback loop was used to fine-tune the expression of the *tph* operon according to available TPA concentrations. The *tph* catabolic operon and the *tphK* transporter were placed under the control of a protocatechuate (PCA)-inducible module^48^ (**Fig. 3a**). The resulting p131CB10-tphKAB plasmid was introduced into *P. putida,* generating strain **tphKAB**. This strain could utilise 10 mM of TPA as a sole carbon and energy source within 24 h (**Fig. 3b**, **Fig. S11a**) at a growth rate of 0.227 ± 0.021 h^-1^ (**Fig. 3c**), reaching a maximum biomass yield of Y_X/S_ of 0.33 g_CDW_ g ^-1^. These rates are consistent with previous reports of engineered *P. putida* equipped with the heterologous *tph* operon from *Comamonas* sp. E6^45,46,49^, reported to afford growth rates on TPA in the range of 0.20 and 0.55^46,49,50^.

**Figure 3.**
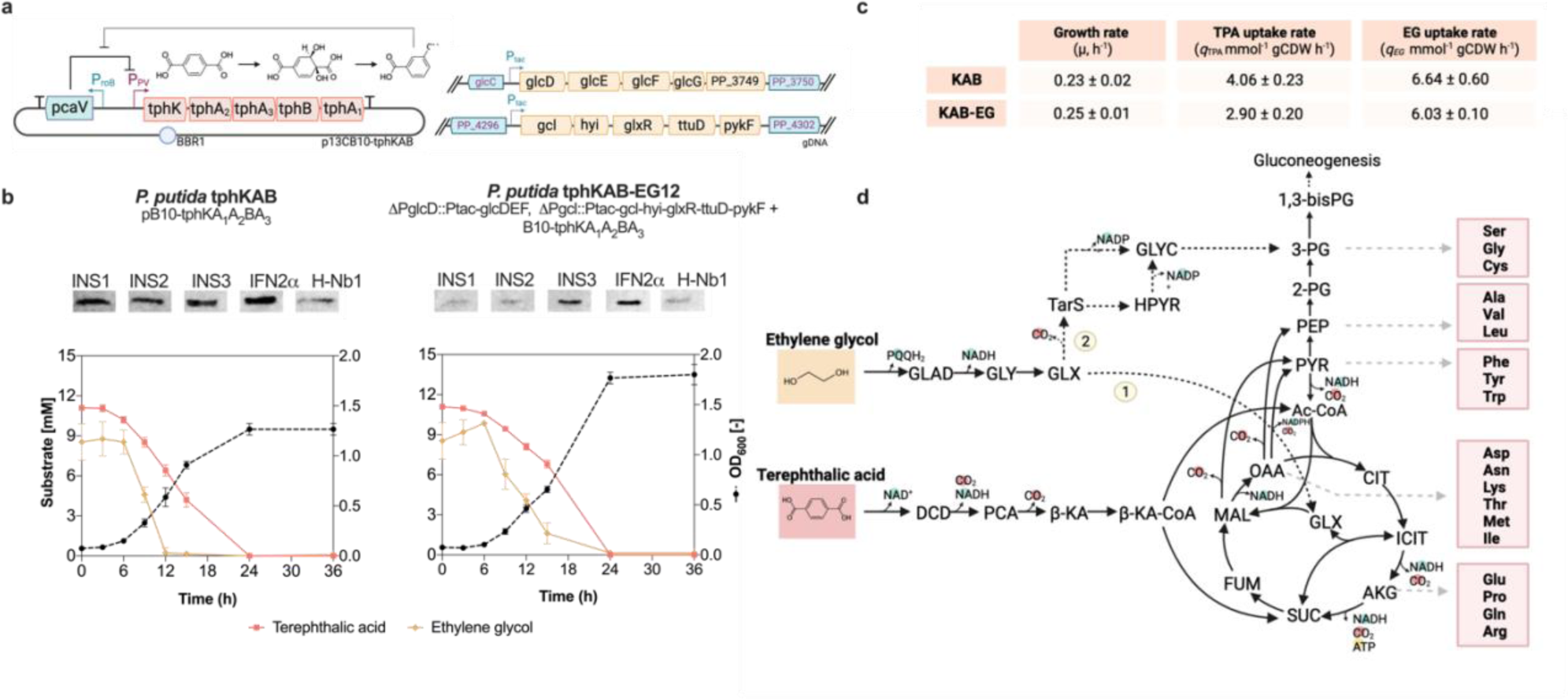
Engineered metabolism and therapeutic protein production from TPA and EG by engineered *P. putida*. (**a**) Schematic of catabolic modules enabling TPA (left) and EG (right) metabolism in *P. putida*. The *tph* operon (*tphA2*_II_*A3*_II_*B*_II_*A1*_II)_ from *Comamonas* sp. E6 and the tphK transporter from *P. mandelii* are placed under the control of a PCA-sensing module^48^. The *glcDEF* (P*_tac_*:*glcDEFG:PP_3750*) and *gcl (gcl*:*hyi*:*glxR*:*ttuD*:*pykF*) operons are overexpressed by downstream introduction of a P_tac_ constitutive promoter. (**b**) Growth, substrate utilisation and production of a panel of long-acting human insulin analogues (INS1, INS2, INS3), human interferon α2a (IFNα2a), and a synthetic HEL4 nanobody (Nb1), from strains tphKAB (harbouring TPA metabolism only) and tphKAB-EG (harbouring TPA and EG metabolism) grown in shake flasks containing 10mM of both TPA and EG. (**c**) Growth rates (μ) and substrate uptake rates (q_SUB_) calculated from flask growth and consumption curves of both strains. (**d**) Metabolic pathways for the assimilation of TPA and EG in the engineered *P. putida* KT2440 strains. In the tphKAB strain, EG is natively funnelled into the glyoxylate shunt (1), while in the tphKAB-EG strain, EG is metabolized to biomass directly from 3-PG (2). For **b** and **c**, all data points are mean ± standard deviation (SD) of n = 3 biological replicates.

#### 3.2.2 Ethylene glycol enhances recombinant therapeutic protein production in engineered *P. putida* KT2440 grown on terephthalic acid

We next set out to further modify this strain to assimilate ethylene glycol (EG), the diol monomer of PET plastic. *P. putida* KT2440 utilises EG for biomass accumulation suboptimally, partly due to complex regulatory mechanisms surrounding C_2_ metabolic pathways^51,52^. One such mechanism involves repression of the glyoxylate carboligase (*gcl*) pathway by GclR. After the conversion of ethylene glycol to glyoxylate by the glycolate oxidase (*glcDEF*) operon, *gcl* can allow EG assimilation into biomass via glycerate metabolism^53^ (**Fig. 2d**). Most engineering efforts to date have thus focused on the deletion of *gclR* or the overexpression of glyoxylate carboligase (*gcl*), which ultimately allow for improved biomass formation via the glycerate pathway^46,49,53,54^. Here we employed a similar approach, whereby both native promoters driving expression of the *gcl* and *glcDEF* operons were exchanged for the strong constitutive P_tac_ promoter in the tphKAB strain. The resulting strain, **tphKAB-EG12** (*P. putida* KT2440 p13CB10:*tphKA2_II_A3_II_B_II_A1_II_*Ptac::*glcDEFG* Ptac::*gcl*), could utilise 10 mM of both TPA and/or EG as the sole carbon and energy source (**Fig. S11b-c**). Moreover, growth in the presence of both TPA and EG was additive, reflected by an increased cell density by almost 30% over that achieved on TPA alone, (OD_600_ of 1.9 and 1.35, respectively), suggesting both monomers are simultaneously funneled into biomass (**Fig. 3**). Interestingly, when grown on both substrates, the parental tphKAB strain could also fully consume EG, albeit without it leading to biomass accumulation (**Fig. 3b**). Though both the tphKAB and tphKAB-EG12 strains exhibited similar growth rates when co-fed TPA and EG (0.227 ± 0.021 h^-1^ and 0.250 ± 0.012 h^-1^, respectively), tphKAB could more efficiently uptake both substrates, while tphKAB-EG12 was able to better assimilate them into biomass, reflected by its lower substrate uptake rates and higher biomass yield (**Fig. 3c**). This is consistent with expectations as natively, the *gcl* pathway is repressed in *P. putida*, and so EG is funneled directly into the glyoxylate shunt, without leading to biomass formation. This shunting has instead been shown to result in a surplus of reducing equivalents when used as an acetate co-feed^51^. We therefore hypothesized that likewise utilising EG as a co-substrate, with the gluconeogenic substrate TPA, could provide elevated concentrations of reducing equivalents as well as increased flux through amino acid biosynthesis precursors, favorable properties for recombinant protein production from PET monomers.

To empirically determine which genetic system (tphKAB or tphKAB-EG12) offered the optimal flux configuration for bioproduction from EG, we compared the ability of both strains to produce a panel of recombinant therapeutic proteins (RTP) when grown on TPA and EG as sole carbon and energy sources. These included several tagged long-acting human insulin analogues (INS1, INS2, INS3)^55^, human interferon α2a (IFNα2a), and a synthetic HEL4 nanobody (Nb1). Both strains were able to produce all recombinant proteins when grown on TPA and EG as the sole carbon and energy sources (**Fig. 3b**). Despite lower biomass accumulation, recombinant protein production titres were on average 7-fold higher in the tphKAB strain, as observed by western blot densitometry (**Fig. 3**). This increase was only observed with EG co-feeding in the tphKAB strain, as yields obtained with this strain grown solely on TPA were similar to those from tphKAB- EG12 (**Fig. S11d**). Due to the observed protein titre enhancement, we utilised tphKAB as the base strain for all further engineering. These results support the notion that additional reducing equivalents, via assimilation of EG through the native pathways rather than the engineered pathways, provided an enhancement in recombinant protein titres. Additionally, the production of therapeutic proteins shown here further broadens the range of products derived from PET plastic valorization ^20,54,56,57^.

#### 3.2.3 Establishing xylose metabolism and obtaining the final engineered *Pseudomonas putida* KT2440 strain

*P. putida* cannot utilise xylose as this strain is not able to metabolize C_5_ sugars. For this purpose, we integrated the D-xylose isomerase pathway from *E. coli* into the tphKAB strain, under the control of the strong P_tac_ promoter. Here, a constitutive promoter was used to avoid the incorporation of catabolite repression elements, that would impede xylose utilisation in the presence of glucose^58^. In this pathway, xylose is transported by the D-xylose:H^+^ symporter (*xylE*) and metabolized via the D-xylose isomerase (*xylA*) and xylulokinase (*xylB*) to xylulose-5-phosphate (X5P) where it can then enter central metabolism. We additionally deleted glucose dehydrogenase (*gcd*) to avoid conversion of xylose to the dead-end product xylonate^59^. As expected, the final strain **tphKAB-XylABE** was able to grow on xylose as a sole carbon and energy source (**Fig. S12**)^19,59^. The deletion of *gcd* resulted in a slight detrimental impact on the growth and uptake rate of tphKAB-XylABE on glucose alone, partly due to ATP-costly glucose uptake via an ABC transporter (GtsABCD)^59^ and the inability to feed into the electron transport chain, although there was no difference in the total biomass achieved (both OD_600_ = 1.7, **Fig. S12**).

Together, the final **tphKAB-XylABE** strain is able to individually assimilate terephthalic acid, ethylene glycol, lactic acid, glucose, xylose, ferulic acid, *p*-coumaric acid, glycerol, and fatty acids as the sole carbon and energy source (**Fig. S13-14**). Although no evidence of toxic compound accumulation or microbial inhibition was observed when growing this final strain directly in the enzymatic hydrolyses (**Fig. S15**), localized concentrations of additives and inhibitors in post-consumer waste streams could potentially interfere with growth^60,61^. To assess the strain’s resilience against potential MSW-derived inhibitory compounds, growth assays were conducted in their presence (**Fig. S16**). The strain displayed good growth profiles in the presence of a variety of commonly found inhibitors at millimolar concentrations. For example, limonene, a common component of clean products reported to be present in landfill at 5-235nM^62^, displayed no inhibitory effect at or below 20mM. Moreover, the observed impact on growth when cultivated with furfural or HMF only above 5mM, suggest that these inhibitors were not present in high millimolar concentrations within our hydrolysates (Fig. 3d). We thus next aimed to explore the growth, carbon efficiency and co-utilization dynamics of the final engineered strain within a mixture of all the major substrates derived from the selected MSW-like fractions.

### 3.3 Engineered *P. putida tphKAB-XylABE* co-utilises a mixture of different substrates derived from mixed municipal solid waste-like feedstocks

First, to gain in-depth understanding on the final strain’s carbon efficiency for each substrate, we constrained a modified genome-scale metabolic model (GEM) of *P. putida* KT2440 with the experimental data and performed flux balance analysis (FBA). The modified model was able to accurately predict the growth rate on each compound (R^2^ = 0.9524, **Fig. S17**), and so we used FBA to obtain predictions of CO_2_ production rates for each substrate, and ultimately their carbon balancing profiles. High yields of biomass per substrate were obtained in descending order with fatty acids, lactic acid, and glycerol (1.34 g_CDW_ g^-1^, 0.69 g_CDW_ g^-1^ and 0.58 g_CDW_ g^-1^, respectively). When using these preferential substrates alone, *P. putida* funneled more than 60% of the substrate’s C-mol content into biomass (**Fig. 4a**). The lowest biomass yield was obtained with TPA and EG, partly due to EG serving as an energy co-substrate only (0.26 g_CDW_ g ^-1^), with a predicted loss of 74% of the total carbon to CO_2_ (**Fig. 4a**). The remaining substrates afforded moderate biomass yields, similar to that of glucose (0.43 g_CDW_ g_GLU_^-1^, 0.48 g_CDW_ g ^-1^, and 0.44 g_CDW_ g ^-1^), channeling 33 to 53% of their C-mol content to biomass (**Fig. 4a**). In terms of growth rate, fatty acids and lactic acid provided the fastest growth (0.466 ± 0.05 h^-1^ and 0.489 ± 0.006 h^-1^ respectively). Moderate growth rates were observed for glucose, TPA and EG, coumaric acid and glycerol, while growth on ferulic acid and xylose were slowest (0.102 ± 0.003 h^-1^ and 0.04 ± 0.002 h^-1^, respectively). These results demonstrate that more reduced substrates – fatty acids, glycerol, and lactic acid – promote higher biomass yields, here showing that less than 35% of their carbon was lost to CO_2,_ consistent with previous modelling studies ^63^. This indicates that using feedstocks composed primarily of reduced monomers, such as triglyceride-enriched waste, could result in more sustainable and efficient processes than those currently performed using carbohydrates, as a higher proportion of carbon is funnelled into biomass accumulation and product formation. Indeed, glycerol and fatty acids have shown promising results for the cost-efficient and sustainable production of polyhydroxyalkanoate (PHA) in *P. putida* ^64–67^.

**Figure 4.**
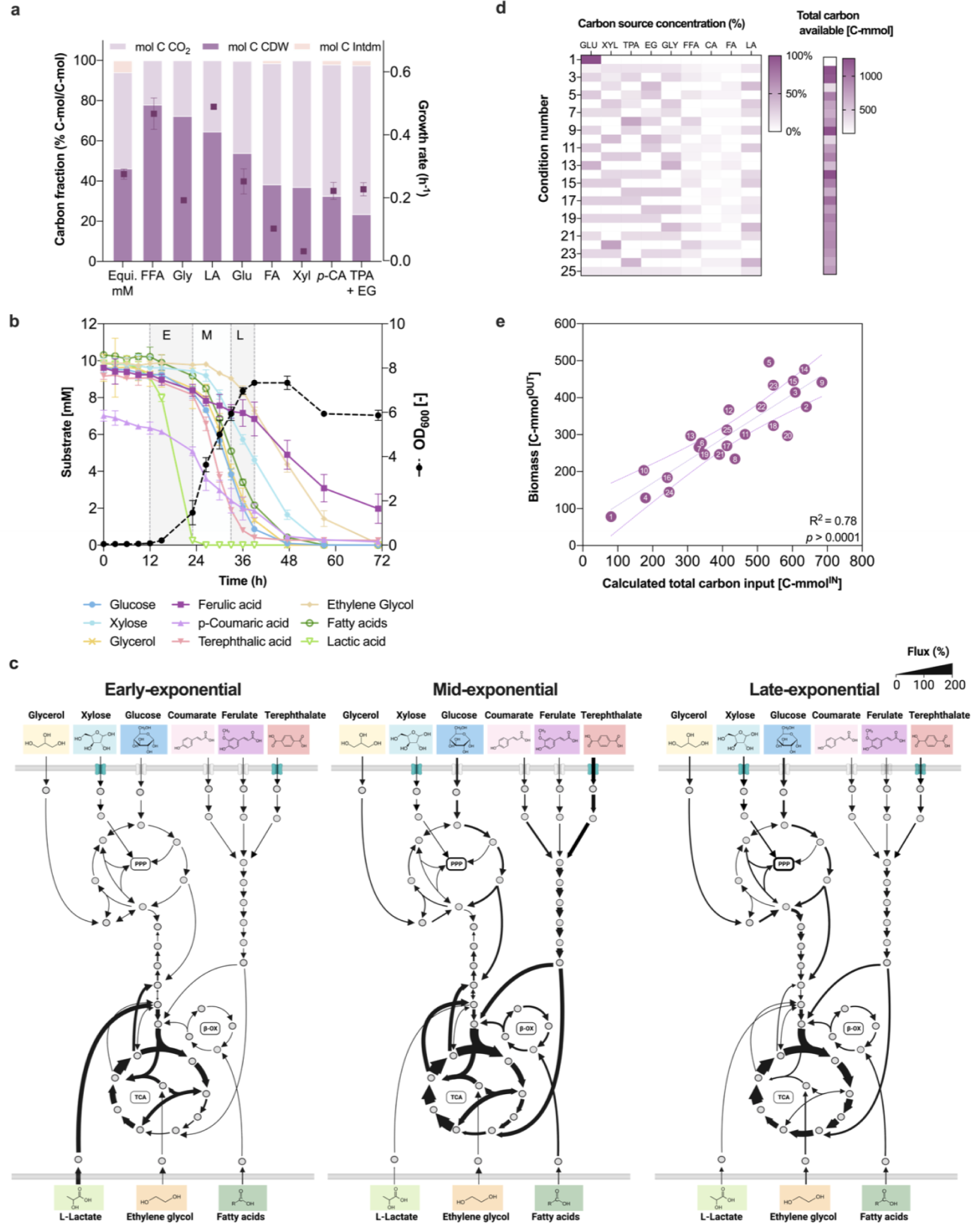
Engineered P. putida KT2440 efficiently assimilates mixtures of the nine major substrates in mixed hydrolysable MSW. (**a**) Growth rates and carbon balance for flask cultivations with individual substrates. Shown are all carbon-containing substrates measured at the end of exponential phase for each monomer, where the carbon fraction represents their relative abundance to the overall input of carbon in C-mol. The amount of CO_2_ produced was calculated from FBA-predicted CO_2_ production rates over the whole growth period. (**b**) Flask cell cultivation profile of tphKAB-XylABE in M9 medium containing equimolar (10 mM) amounts of the nine monomeric substrates (1.74 g L-1 glucose, 1.49 g L-1 xylose, 1.86 g L-1 ferulic acid, 1.45 g L-1 p-coumaric acid, 0.89 g L-1 lactic acid, 1.93 g L-1 terephthalic acid, 0.61 g L-1 ethylene glycol, 0.88 g L-1 glycerol and 2.47 g L-1 fatty acid). (**c**) Predicted flux distributions for the assimilation of the nine monomers in the equimolar cultivation during early, mid- and late-exponential phase. (**d**) Layout of the 25 selected treatments containing different concentrations of each substrate as proposed by design of experiments, and the total carbon available from these substrates in C-mol. (**e**) Correlation between the achieved biomass in C-mmol and the calculated total input of biomass-producing carbon. *OD_600_* represents the optical density measured at 600 nm. For **a**, **b**, and **e**, all data points are mean ± SD of n = 3 biological replicates.

We then assessed the simultaneous utilization of all nine monomers by conducting flask growth assays in minimal medium containing equimolar (∼10 mM) concentrations of each carbon source (**Fig. 4b**). This cultivation afforded a high maximum OD_600_ of 7.4, reaching a final combined biomass yield of 0.47 g_CDW_ g_SUB_^-1^ at a growth rate of 0.275 ± 0.016 h^-1^. Lactic acid was preferentially consumed during early exponential phase (12-23 h), along with small amounts of glucose, fatty acids, and aromatics (**Fig. 4b**). Lactic acid contributed roughly the same amount to biomass accumulation as when present as a sole carbon source, albeit at a slower uptake rate (10.5 ± 1.2 vs 2.3 ± 0.2 mmol g_CDW_^-1^ h^-1^). During this early phase, our model predicts an active glyoxylate shunt in the TCA cycle, and a central gluconeogenic configuration predominantly driven by lactic acid metabolism and from minor additional acetyl-CoA and intermediate organic acid influx from β-oxidation and aromatic degradation, respectively^68^ (**Fig. 4c & Fig. S18**). During mid-(23-30 h) and late-exponential (30-39h) phase, the remaining carbon sources were utilised simultaneously (**Fig. 4b & Fig. S18**), and TPA, glycerol, glucose and fatty acids were depleted by 48 hours. This period drove most biomass growth, reaching a maximum OD_600_ of 7.4. Flux predictions during mid-exponential phase, indicate all substrates were being co-utilised, although gluconeogenic substrates were used preferentially, resulting in especially strong flux through TCA and glyoxylate shunt reactions (**Fig. 4c**). In late exponential phase however, the central nodes were predicted to shift to a glycolytic directionality, likely due to the later onset of xylose consumption (**Fig. 4c**), which was assimilated more rapidly than when provided alone (**Fig. 4c & Fig. S13**). No further cell biomass accumulated beyond 39 h, even though substrates capable of contributing to growth remained, including 6.8 mM of ferulic acid and 1.86 mM of coumaric acid. Although these were partially depleted by mid-exponential phase, they had not completely journeyed through the lower aromatic catabolic pathway, and we observed a build-up of the respective pathway intermediates vanillic acid and 4-hydroxybenzoic acid (**Fig. S19**). Only after TPA had been completely depleted did their consumption begin, being completely depleted by 57 h and 72 h. The apparent lack of significant lag or diauxic interruption to growth (**Fig. 4b**), can likely be attributed to the reverse carbon catabolite repression (rCCR) mechanism present in *P. putida*, which operates at the mRNA level – contrary to transcriptional CCR in microorganisms like *E. coli* and *B. subtilis*^69–71^. This post-transcriptional regulation has evident physicochemical advantages, which allow for an extremely rapid and sensitive response to substrate availability. In *P. putida,* this mechanism confers a preference for gluconeogenic substrates^18,71,72^, although studies have previously demonstrated that in nutritionally complex environments, glycolytic and gluconeogenic co-utilisation can coexist^73–76^. Previous strategies to allow for substrate co-utilisation have consisted of *crc* elimination^19,20^, however this has shown to be detrimental in some instances^68^.

This metabolic analysis demonstrates how when using complex mixed substrates, *P. putida* can quickly and dynamically change the configuration of its central and peripheral metabolic pathways in response to changing carbon and energy availability. This trait is especially pertinent in the context of changing substrate ratios and concentrations from varying mixed waste compositions. To further illustrate this, we explored the growth of the tphKAB-XylABE strain on a variety of possible combinations of the nine monomers. We employed a design of experiments (DoE) approach to construct a set of treatments that would explore the full combinatorial space of substrate loadings (**Table S7**). Each substrate was provided in the respective treatment at either 0, 25 or 50 mM, except for ferulic and coumaric acid which were supplied at 0, 5 or 10 mM and fatty acids at 0, 12.5 or 25 mM (low solubility at higher concentrations), and grown in minimal media for 72 h. The final set of 25 conditions tested allowed us to investigate the vast number of possible scenarios at which each substrate might be found in process relevant conditions (**Fig. 4d**). Robust biomass yields from the combined substrates were obtained consistently across all conditions after 72h, ranging from 0.21g_CDW_ g ^-1^ to 0.61g g ^-1^. When accounting for the percentage of C-mmol that each substrate contributes to biomass, calculated from the individual growth experiments, we observed a correlation between the amount of calculated carbon input (C-mmol^IN^) contributing to biomass from each substrate and the final biomass obtained (C-mmol^OUT^) across the 25 conditions (R^2^ = 0.788, p-value < 0.001) (**Fig. 4e**). This reiterates the fact that, regardless of the amount and concentration of each available substrate, the engineered *P. putida* strain tphKAB-XylABE can efficiently and consistently convert the available carbon into biomass.

### 3.4 Mixed MSW-like derived substrates afford sufficient carbon for efficient RTP and PHA production

To demonstrate a proof-of-concept for bioproduction from complex feedstocks, we assessed the ability of the engineered tphKAB-XylABE strain to produce valuable products, including a recombinant therapeutic protein (RTP) and endogenously produced polyhydroxyalkanoates (PHA), from mixed MSW-derived substrate compositions. Working on the knowledge obtained from earlier in this work, we investigated production of a human therapeutic protein, IFNα2a, as a high-value illustrative product. For this, we utilised the previously generated p131CB10-tphKAB-IFN2a plasmid, and transformed it into the tphKAB-XylABE strain. *P. putida* KT2440 natively synthesizes PHAs from acetyl-CoA under nitrogen-limiting conditions^44^. We therefore sought to separately produce PHAs as a low-value illustrative product without any additional engineering.

From enzymatic hydrolysis of the MSW-like feedstocks (**Fig. 2**), the resulting liquor was directly used for microbial cultivation in microbioreactors. Engineered tphKAB-XylABE grew efficiently on both hydrolysis liquor and synthetic media with equivalent substrate concentrations (mocks) (**Fig. S15**), suggesting minimal release of additional metabolized substrates or inhibitory compounds, and no performance loss. We therefore performed the proof-of-concept bioproduction using synthetic mocks as a proxy for the real MSW hydrolysate. Scale-up flask cell growth assays were performed with both the high and low-income waste-derived substrate compositions for both products. Growth and consumption dynamics were similar for both products in each waste composition, and after 60h all biomass-producing substrates were completely depleted (**Fig. 5**). Higher biomass yields were obtained in the low-income cultivations, yielding 0.52 g_CDW_/g_SUB_ for IFNa2a and 0.56 g_CDW_/g_SUB_ for the PHA, whereas 0.47 g_CDW_/g_SUB_ and 0.49 g_CDW_/g_SUB_ were obtained in the high-income (IFNa2a and PHAs, respectively).

**Figure 5.**
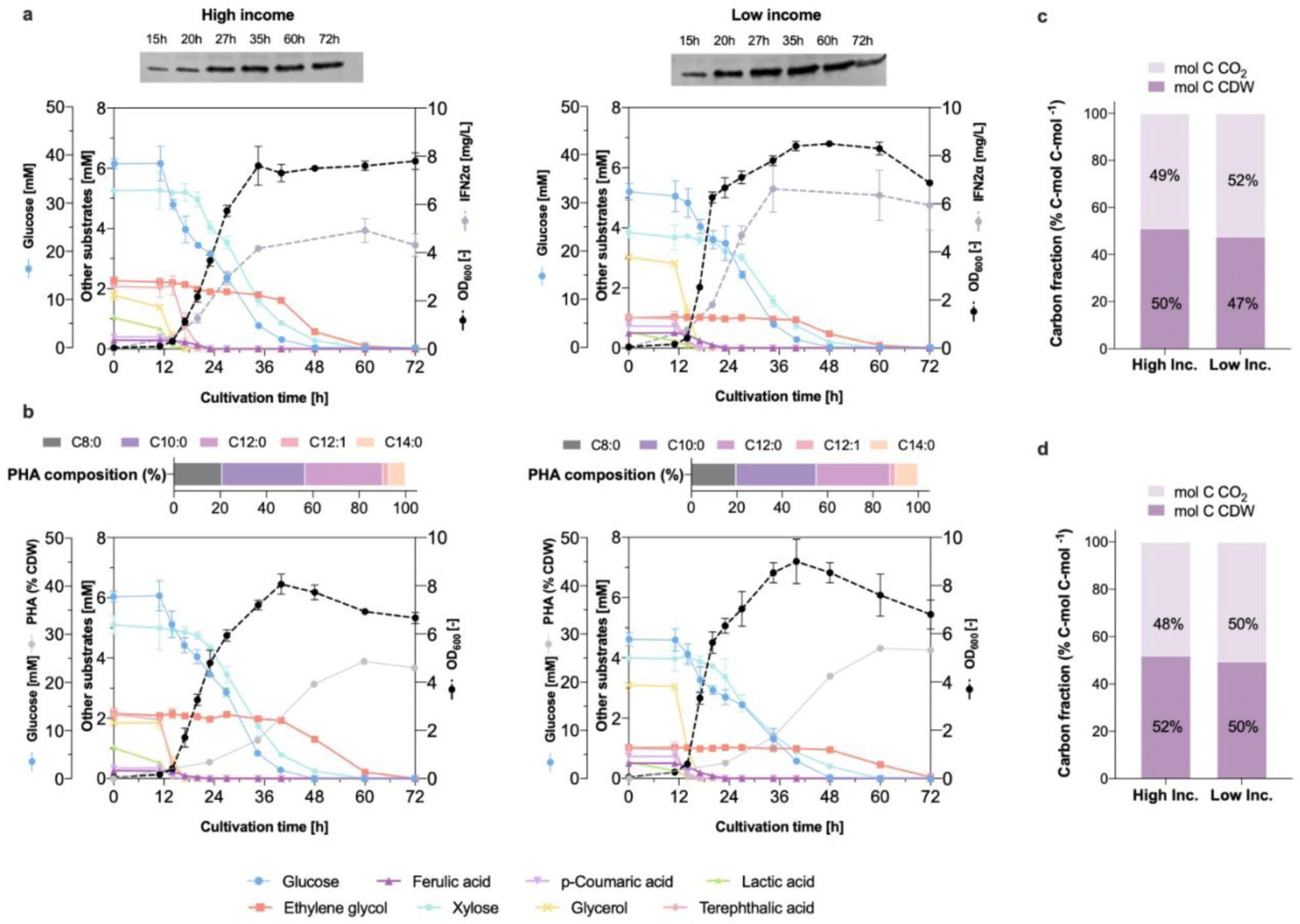
Bioconversion of mixed municipal solid waste derived substrates compositions into biotherapeutics and biopolymer. (**a**) Bacterial growth, substrate utilisation and IFNα2a accumulation from tphKAB-XylABE in high-(left) and low-income (right) waste derived substrate compositions in flask cultivations. Western blots show total cell samples taken at periodic time intervals. (**b**) Bacterial growth, substrate utilisation and PHA accumulation from tphKAB-XylABE in high-(left) and low-income (right) waste derived substrate compositions in flask cultivations. Shown above are compositions of PHA produced from both hydrolysates. (**c**) Carbon balancing profiles for IFNα2a producing cultivations. (**d**) Carbon balancing profiles for PHA-producing cultivations. For **a** and **b**, all data points are mean ± SD of n = 3 biological replicates.

Expression of IFNα2a was successfully observed in both feedstock compositions, although to a slightly higher concentration in the low-income hydrolysate (3.9 mg L^-1^ and 5.3 mg L^-1^, respectively) (**Fig. 5a**). In both mock hydrolysates, RTP production, as well as biomass, reached a maximum titre by about 36 h, even though some carbon was still available. However, protein yield remained stable, and no significant protein degradation was seen after a further 36h, concurring with the onset of ethylene glycol metabolism. Production of PHA was likewise successfully observed in both mock hydrolysates, reaching maximum yields of 23.06 ± 0.95% and 26.67 ± 0.65%of the cell dry weight (%CDW) after 60 h, consistent with reported yields for PHA production in the native host^44,54,66^. The PHAs produced from both substrate compositions primarily consisted of 3-hydroxydecanoic acid and 3-hydroxydodecanoic acid (**Fig. 5b**). The delayed onset of PHA production compared to RTP production is explained by the slow depletion of available nitrogen and differences in carbon contribution to each product (**Fig. S20**). Together, although the total carbon amount in the low-income substrate composition was higher, both high- and low-income cultivations efficiently funnelled on average a total of 50% of the available carbon into biomass (**Fig. 5c-d**). This again suggests that the engineered strain is able to efficiently utilise the available carbon both growth and bioproduction. Genetic stability is a key metric for bioprocess robustness, ensuring scalability and productivity. As the recombinant protein production was plasmid encoded, we assessed the long-term stability of the biosynthetic plasmid (**Fig. S21**). All strains retained the plasmid after 27 generations, whereas after 65 generations growth functionality was impeded in the absence of any plasmid selection, and plasmid selection with antibiotic-only led to partially impeded growth. This trend was furthered at later timepoints. This would suggest the need for microbial re-dosing strategies and/or use of engineered strains with biosynthetic loci stably integrated in the genome. Furthermore, the product exemplars used here serve only as illustrative high- and low-value products that could be produced using mixed MSW as feedstock, and it is worth noting that further optimisation work will access higher RTP and PHA production titres in an effort to reach greater economic viability ^44,77–79^. Whilst there are clearly several orders of magnitude difference between the annual tonnage of MSW produced and the requirement for very high-value products, such as therapeutic RTPs, these results suggest the potential to achieve both economic and environmental sustainability, by producing an array of products with different value and volume profiles in tandem.

### 3.5 Combined prospective life cycle assessment, life cycle costing and hotspot analysis demonstrates the environmental and economic sustainability benefits of using MSW as feedstock for biomanufacturing of PHA

To examine the environmental and economic sustainability of the proposed process, we conducted prospective life cycle assessment (LCA) and life cycle costing (LCC) studies of a scaled-up bioprocess. The climate change (CC) impact and economic feasibility of the MSW valorisation for both high- and low-income country waste compositions were compared to conventional treatment options (**Fig. 6**). LCA results were compared with incineration with energy recovery, landfill, and open burning, whilst LCC comparisons excluded the latter due to lack of data^41^. We focused on the production of PHA because of scale-up data availability and volume profile of bioplastics production and consumption. The total CC impact and LCC were estimated to be 0.31 kg CO_2_ eq./t and $111.4/t for the high-income and 0.28 kg CO_2_ eq./t and $96.3/t for the low-income MSW scenarios (**Fig. 6c**). The enzymatic hydrolysis step was identified as the main hotspot of this system for both CC impact and LCC, as it contributes around 90% of the total CC impacts (**Table S9-10**, **Fig. S22**) and around 70% of the costs, largely due to the consumption of enzymes and buffers. The microbial growth / PHA bioproduction stage is responsible for 14% and 21% of the total CC impact and LCC, respectively, largely due to the buffers and media, e.g. sodium phosphate and potassium phosphate. The PHA extraction step, on the other hand, accounts for less than 5% for both CC and LCC results. In the case of LCC, it was found that CAPEX contributions for the whole process are less than 5%.

**Figure 6.**
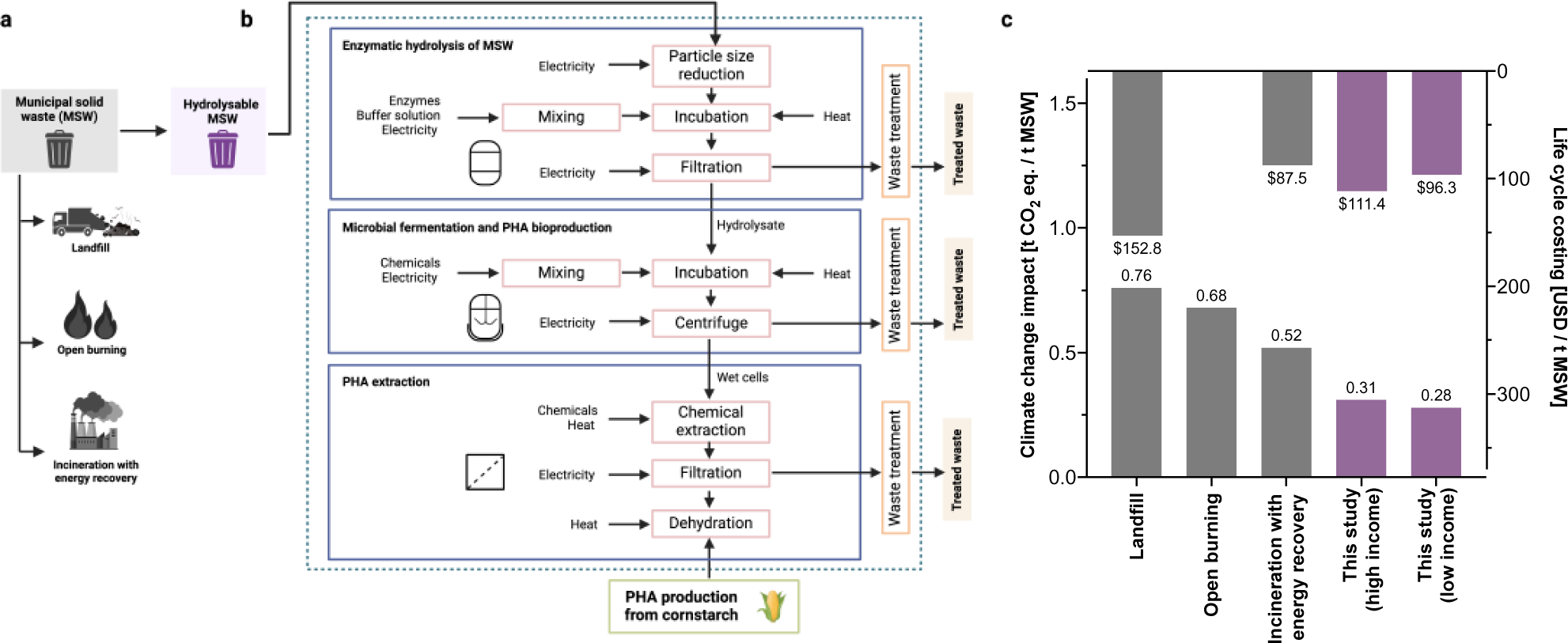
Comparison of current and proposed routes for MSW disposal and use. (**a**) Current dominant processes for MSW disposal and use. (**b**) Proposed use of complex hydrolysable MSW as feedstock for advanced biorefinery and system boundaries used for LCA and LCC. (**c**) Climate change and LCC results of current and proposed MSW disposal and uses, contributions from each production steps are provided in **Fig. S22**.

The LCA results on the proposed bioprocess show significant potential for reducing the carbon footprint of municipal solid waste (MSW) treatment, with a 41-62% reduction compared to conventional options. The LCC results are 27-37% lower when compared to landfill, whilst for incineration with energy recovery, these were found to be 10-27% higher as incineration will obtain higher revenues from selling energy. Although the LCA and LCC results indicated that the use of MSW as a feedstock for microbial fermentation and bioconversion is a promising alternative to the current disposal options, further process improvements are needed, particularly in the hydrolytic pre-treatment step. A wealth of studies aimed at developing efficient physiochemical and enzymatic deconstruction strategies for the individual polymers and macromolecules that comprise the constituent fractions of MSW waste exist^80–83^, and can be leveraged towards developing efficient mixed fraction hydrolysis. ^84,85^. As the engineered *P. putida* chassis can directly co-utilise complex substrate mixtures, any hydrolysis reaction that can generate monomers that can be fed into metabolism could be incorporated. For instance, a mixture of non-biodegradable plastics was recently depolymerised by metal-catalysed autoxidation^20^, and various alternative pre-treatment methods exist for lignocellulose and plastics, such as chemical hydrolysis, chemical glycolysis or thermolysis^57,86–93^. Further enzymatic optimisation efforts together with the discovery and engineering of novel enzymatic cocktails – especially those working under mild and mesophilic conditions, will aid the development of sustainable mixed MSW pre-treatment strategies^94^. Enhanced pre-treatment options coupled with combined LCA and LCC analysis, such as those reported here, will permit informed strategies to balance and maximise both environment and economic outcomes.

## 4. Conclusion

Collectively, the results of this study provide an illustration into the considerable potential of leveraging *P. putida* as a *super host* organism for bioproduction from variable and complex waste feedstocks, including MSW-like feedstocks. We hereby shed light on the inherent metabolic versatility and consistency of the engineered *P. putida* strain, laying a foundation for the consistent bioconversion of complex feedstocks – irrespective of their compositional variability – into added value bioproducts. While current global management efforts are being made to prevent, reduce, and recycle waste, its generation will undoubtedly never be eliminated. In a circular bio-based economy, this unavoidable waste can be utilised as a resource, and maintained for as long as possible in a closed loop. Under this philosophy, new biomanufacturing and advanced biorefinery approaches will therefore be critical to derive value from waste.

## Authors′ contributions

GAG, MC, AB, KF, planned and performed the experiments. PG and RCF performed the LCA and interpreted the results. GAG, MC, and ND analysed the data and wrote the manuscript. All authors read and approved the final version of the manuscript.

## Data availability

All data generated or analysed in this manuscript is available as supplementary data files. Data for all main text and supplementary figures are available in supplementary data file 1. The modified metabolic model is available in https://github.com/GuadalupeAlvarezGonzalez/KAB-XylABE. Any additional data can be made available from the authors upon reasonable request. Materials can be made available upon reasonable request under a material transfer agreement.

## Acknowledgements

We would like to acknowledge Rehana Sung and the MIB separation facility, Ross Kent for supporting early catabolic constructs assembly efforts, and Joseph Webb for supporting with fermentation.

## References

1. United Nations. The sustainable development goals report 2022. United Nations Publ. issued by Dep. Econ. Soc. Aff. 68 (2022).

2. Friedlingstein, P. et al. Global Carbon Budget 2021. Earth Syst. Sci. Data 14, 1917–2005 (2022).

3. WEF. The Global Risks Report *2023* 18th Edition. World Economic Forum www.weforum.org (2023).

4. UNFCC. *Adoption of the Paris Agreement*. *Conference of the Parties on its twenty-first session* vol. 21932 http://unfccc.int/resource/docs/2015/cop21/eng/l09r01.pdf (2015).

5. Intergovernmental Panel on Climate Change. Climate Change 2022 - Mitigation of Climate Change - Full Report. Cambridge Univ. Press 1–30 (2022).

6. Culaba, A. B. et al. Design of biorefineries towards carbon neutrality: A critical review. Bioresour. Technol. 369, 128256 (2022).

7. Calvo-Flores, F. G. & Martin-Martinez, F. J. Biorefineries: Achievements and challenges for a bio-based economy. Front. Chem. 10, 1329 (2022).

8. Mohr, A. & Raman, S. Lessons from first generation biofuels and implications for the sustainability appraisal of second generation biofuels. Effic. Sustain. Biofuel Prod. Environ. Land-Use Res. 63, 281– 310 (2015).

9. Magnan, A. K. et al. Sea level rise risks and societal adaptation benefits in low-lying coastal areas. Sci. Rep. 12, 1–22 (2022).

10. Kulp, S. A. & Strauss, B. H. New elevation data triple estimates of global vulnerability to sea-level rise and coastal flooding. Nat. Commun. 10, 1–12 (2019).

11. Zambare, V. P. & Christopher, L. P. Integrated biorefinery approach to utilization of pulp and paper mill sludge for value-added products. J. Clean. Prod. 274, 122791 (2020).

12. Gaspar, M. C. et al. Assessment of Agroforestry Residues: Their Potential within the Biorefinery Context. ACS Sustain. Chem. Eng. 7, 17154–17165 (2019).

13. Teigiserova, D. A., Hamelin, L. & Thomsen, M. Review of high-value food waste and food residues biorefineries with focus on unavoidable wastes from processing. Resour. Conserv. Recycl. 149, 413– 426 (2019).

14. Kaparaju, P., Serrano, M., Thomsen, A. B., Kongjan, P. & Angelidaki, I. Bioethanol, biohydrogen and biogas production from wheat straw in a biorefinery concept. Bioresour. Technol. 100, 2562–2568 (2009).

15. Wu, Y., Shen, X., Yuan, Q. & Yan, Y. Metabolic engineering strategies for co-utilization of carbon sources in microbes. Bioengineering 3, (2016).

16. Weimer, A., Kohlstedt, M., Volke, D. C., Nikel, P. I. & Wittmann, C. Industrial biotechnology of Pseudomonas putida: advances and prospects. Appl. Microbiol. Biotechnol. 104, 7745–7766 (2020).

17. Nikel, P. I. & de Lorenzo, V. Pseudomonas putida as a functional chassis for industrial biocatalysis: From native biochemistry to trans-metabolism. Metab. Eng. 50, 142–155 (2018).

18. Schink, S. J. et al. Glycolysis/gluconeogenesis specialization in microbes is driven by biochemical constraints of flux sensing. Mol. Syst. Biol. 18, 10704 (2022).

19. Elmore, J. R. et al. Engineered Pseudomonas putida simultaneously catabolizes five major components of corn stover lignocellulose: Glucose, xylose, arabinose, p-coumaric acid, and acetic acid. Metab. Eng. 62, 62–71 (2020).

20. Sullivan, K. P. et al. Mixed plastics waste valorization through tandem chemical oxidation and biological funneling. Science (80-. ). 378, (2022).

21. Vinti, G. et al. Municipal Solid Waste Management and Adverse Health Outcomes: A Systematic Review. Int. J. Environ. Res. Public Health 18, (2021).

22. Kaza, S., Yao, L. C., Bhada-Tata, P. & Van Woerden, F. *What a Waste 2.0: A Global Snapshot of Solid Waste Management to* 2050. World Bank Publications (2018) doi:10.1596/978-1-4648-1329-0.

23. Abbasi, S. A. The myth and the reality of energy recovery from municipal solid waste. Energy. Sustain. Soc. 8, 1–15 (2018).

24. Nizami, A. S. et al. Waste biorefineries: Enabling circular economies in developing countries. Bioresour. Technol. 241, 1101–1117 (2017).

25. Vergara, S. E. & Tchobanoglous, G. Municipal solid waste and the environment: A global perspective. Annu. Rev. Environ. Resour. 37, 277–309 (2012).

26. Marx, C. J. Development of a broad-host-range sacB-based vector for unmarked allelic exchange. BMC Res. Notes 1, 1–8 (2008).

27. Tournier, V. et al. An engineered PET depolymerase to break down and recycle plastic bottles. Nature 580, 216–219 (2020).

28. Nogales, J. et al. High-quality genome-scale metabolic modelling of Pseudomonas putida highlights its broad metabolic capabilities. Environ. Microbiol. 22, 255–269 (2020).

29. Heirendt, L. et al. Creation and analysis of biochemical constraint-based models using the COBRA Toolbox v.3.0. Nat. Protoc. 14, 639–702 (2019).

30. van Duuren, J. B. J. H. et al. Reconciling in vivo and in silico key biological parameters of Pseudomonas putida KT2440 during growth on glucose under carbon-limited condition. BMC Biotechnol. 13, 1–13 (2013).

31. ISO. Environmental management - Life cycle assessment - Requirements and guidelines (ISO 14044:2006). Iso 14044:2006 https://www.iso.org/standard/38498.html (2020).

32. ISO. Environmental Management - Life Cycle Assessment - Principles and Framework (ISO 14040:2006). British Standard vol. 3 https://www.iso.org/standard/37456.html (2004).

33. Swarr, T. E. et al. Environmental life-cycle costing: A code of practice. Int. J. Life Cycle Assess. 16, 389–391 (2011).

34. Zhong, Z. W., Song, B. & Huang, C. X. Environmental impacts of three polyhydroxyalkanoate (pha) manufacturing processes. Mater. Manuf. Process. 24, 519–523 (2009).

35. Andreasi Bassi, S., Boldrin, A., Frenna, G. & Astrup, T. F. An environmental and economic assessment of bioplastic from urban biowaste. The example of polyhydroxyalkanoate. Bioresour. Technol. 327, 124813 (2021).

36. Moreno, J. et al. Life-cycle sustainability of biomass-derived sorbitol: Proposing technological alternatives for improving the environmental profile of a bio-refinery platform molecule. J. Clean. Prod. 250, 119568 (2020).

37. 37. ecoQuery - EcoInvent. https://ecoquery.ecoinvent.org/3.8/apos/search.

38. Castellanos-Beltran, I. J., de Medeiros, F. G. M., Bensebaa, F. & De Vasconcelos, B. R. Novel bottom-up methodology to build the lifecycle inventory of unit operations: the impact of macroscopic components. Int. J. Life Cycle Assess. 28, 669–683 (2023).

39. Sphera. Life Cycle Assessment Software (GaBi). https://sphera.com/life-cycle-assessment-lca-software/ (2023).

40. Huijbregts, M. A. J. et al. ReCiPe2016: a harmonised life cycle impact assessment method at midpoint and endpoint level. Int. J. Life Cycle Assess. 22, 138–147 (2017).

41. Slorach, P. C., Jeswani, H. K., Cuéllar-Franca, R. & Azapagic, A. Environmental and economic implications of recovering resources from food waste in a circular economy. Sci. Total Environ. 693, 133516 (2019).

42. ChemEngOnline. Chemical Engineering Plant Cost Index. Chemical Engineering 182 https://www.chemengonline.com/site/plant-cost-index/ (2020).

43. Linger, J. G. et al. Lignin valorization through integrated biological funneling and chemical catalysis. Proc. Natl. Acad. Sci. U. S. A. 111, 12013–12018 (2014).

44. Mezzina, M. P., Manoli, M. T., Prieto, M. A. & Nikel, P. I. Engineering Native and Synthetic Pathways in Pseudomonas putida for the Production of Tailored Polyhydroxyalkanoates. Biotechnol. J. 16, (2021).

45. Sasoh, M. et al. Characterization of the terephthalate degradation genes of Comamonas sp. strain E6. Appl. Environ. Microbiol. 72, 1825–1832 (2006).

46. Werner, A. Z. et al. Tandem chemical deconstruction and biological upcycling of poly(ethylene terephthalate) to β-ketoadipic acid by Pseudomonas putida KT2440. Metab. Eng. 67, 250–261 (2021).

47. Salvador, M. et al. Microbial genes for a circular and sustainable bio-PET economy. Genes vol. 10 (2019).

48. Berepiki, A., Kent, R., MacHado, L. F. M. & Dixon, N. Development of High-Performance Whole Cell Biosensors Aided by Statistical Modeling. ACS Synth. Biol. 9, 576–589 (2020).

49. Brandenberg, O. F., Schubert, O. T. & Kruglyak, L. Towards synthetic PETtrophy: Engineering Pseudomonas putida for concurrent polyethylene terephthalate (PET) monomer metabolism and PET hydrolase expression. Microb. Cell Fact. 21, 1–31 (2022).

50. Liu, P. et al. Valorization of Polyethylene Terephthalate to Muconic Acid by Engineering Pseudomonas Putida. Int. J. Mol. Sci. 23, (2022).

51. Li, W. J. et al. Laboratory evolution reveals the metabolic and regulatory basis of ethylene glycol metabolism by Pseudomonas putida KT2440. Environ. Microbiol. 21, 3669–3682 (2019).

52. Wirth, N. T. et al. A synthetic C2 auxotroph of Pseudomonas putida for evolutionary engineering of alternative sugar catabolic routes. Metab. Eng. 74, 83–97 (2022).

53. Franden, M. A. et al. Engineering Pseudomonas putida KT2440 for efficient ethylene glycol utilization. Metab. Eng. 48, 197–207 (2018).

54. Tiso, T. et al. Towards bio-upcycling of polyethylene terephthalate. Metab. Eng. 66, 167–178 (2021).

55. Mikiewicz, D. et al. Soluble insulin analogs combining rapid- and long-acting hypoglycemic properties from an efficient E. coli expression system to a pharmaceutical formulation. PLoS One 12, e0172600 (2017).

56. Sadler, J. C. & Wallace, S. Microbial synthesis of vanillin from waste poly(ethylene terephthalate). Green Chem. 23, 4665–4672 (2021).

57. Diao, J., Hu, Y., Tian, Y., Carr, R. & Moon, T. S. Upcycling of poly(ethylene terephthalate) to produce high-value bio-products. Cell Rep. 42, 111908 (2023).

58. Kaplan, N. A. et al. Simultaneous glucose and xylose utilization by an Escherichia coli catabolite repression mutant. Appl. Environ. Microbiol. 90, (2024).

59. Dvořák, P. & de Lorenzo, V. Refactoring the upper sugar metabolism of Pseudomonas putida for co-utilization of cellobiose, xylose, and glucose. Metab. Eng. 48, 94–108 (2018).

60. Rao, P. & Rathod, V. Valorization of Food and Agricultural Waste: A Step towards Greener Future. Chem. Rec. 19, 1858–1871 (2019).

61. Ügdüler, S., Van Geem, K. M., Roosen, M., Delbeke, E. I. P. & De Meester, S. Challenges and opportunities of solvent-based additive extraction methods for plastic recycling. Waste Manag. 104, 148–182 (2020).

62. Masoner, J. R. et al. Contaminants of emerging concern in fresh leachate from landfills in the conterminous United States. Environ. Sci. Process. Impacts 16, 2335–2354 (2014).

63. Tiso, T. et al. The metabolic potential of plastics as biotechnological carbon sources – Review and targets for the future. Metab. Eng. 71, 77–98 (2022).

64. Beckers, V., Poblete-Castro, I., Tomasch, J. & Wittmann, C. Integrated analysis of gene expression and metabolic fluxes in PHA-producing Pseudomonas putida grown on glycerol. Microb. Cell Fact. 15, 1–18 (2016).

65. Prieto, A. et al. A holistic view of polyhydroxyalkanoate metabolism in Pseudomonas putida. Environ. Microbiol. 18, 341–357 (2016).

66. Kang, D. K. et al. Production of polyhydroxyalkanoates from sludge palm oil using pseudomonas putida S12. J. Microbiol. Biotechnol. 27, 990–994 (2017).

67. Kellerhals, M. B., Kessler, B., Witholt, B., Tchouboukov, A. & Brandl, H. Renewable long-chain fatty acids for production of biodegradable medium-chain-length polyhydroxyalkanoates (mcl-PHAs) at laboratory and pilot plant scales. Macromolecules 33, 4690–4698 (2000).

68. Molina, L., Rosa, R. La, Nogales, J. & Rojo, F. Pseudomonas putida KT2440 metabolism undergoes sequential modifications during exponential growth in a complete medium as compounds are gradually consumed. Environ. Microbiol. 21, 2375–2390 (2019).

69. Lv, F. et al. Regulation of hierarchical carbon substrate utilization, nitrogen fixation, and root colonization by the Hfq/Crc/CrcZY genes in Pseudomonas stutzeri. iScience 25, 105663 (2022).

70. Green, J. et al. Cyclic-AMP and bacterial cyclic-AMP receptor proteins revisited: adaptation for different ecological niches. Curr. Opin. Microbiol. 18, 1–7 (2014).

71. Park, H., McGill, S. L., Arnold, A. D. & Carlson, R. P. Pseudomonad reverse carbon catabolite repression, interspecies metabolite exchange, and consortial division of labor. Cell. Mol. Life Sci. 2019 773 **77**, 395–413 (2019).

72. Wilkes, R. A., Waldbauer, J. & Aristilde, L. Analogous Metabolic Decoupling in Pseudomonas putida and Comamonas testosteroni Implies Energetic Bypass to Facilitate Gluconeogenic Growth. MBio 12, (2021).

73. Kukurugya, M. A. et al. Multi-omics analysis unravels a segregated metabolic flux network that tunes co-utilization of sugar and aromatic carbons in Pseudomonas putida. J. Biol. Chem. 294, 8464–8479 (2019).

74. Perrin, E. et al. Diauxie and co-utilization of carbon sources can coexist during bacterial growth in nutritionally complex environments. Nat. Commun. 11, 1–16 (2020).

75. La Rosa, R., Nogales, J. & Rojo, F. The Crc/CrcZ-CrcY global regulatory system helps the integration of gluconeogenic and glycolytic metabolism in Pseudomonas putida. Environ. Microbiol. 17, 3362–3378 (2015).

76. Wilkes, R. A. et al. Complex regulation in a Comamonas platform for diverse aromatic carbon metabolism. Nat. Chem. Biol. 1–12 (2023) doi:10.1038/S41589-022-01237-7.

77. Meyers, A., Furtmann, C., Gesing, K., Tozakidis, I. E. P. & Jose, J. Cell density-dependent auto-inducible promoters for expression of recombinant proteins in Pseudomonas putida. Microb. Biotechnol. 12, 1003–1013 (2019).

78. Liang, T. et al. Construction of T7-Like Expression System in Pseudomonas putida KT2440 to Enhance the Heterologous Expression Level. Front. Chem. 9, 556 (2021).

79. Lieder, S., Nikel, P. I., de Lorenzo, V. & Takors, R. Genome reduction boosts heterologous gene expression in Pseudomonas putida. Microb. Cell Fact. 14, 1–14 (2015).

80. Blazek, J. & Gilbert, E. P. Effect of enzymatic hydrolysis on native starch granule structure. Biomacromolecules 11, 3275–3289 (2010).

81. Van Dyk, J. S. & Pletschke, B. I. A review of lignocellulose bioconversion using enzymatic hydrolysis and synergistic cooperation between enzymes-Factors affecting enzymes, conversion and synergy. Biotechnol. Adv. 30, 1458–1480 (2012).

82. Chandra, P., Enespa, Singh, R. & Arora, P. K. Microbial lipases and their industrial applications: A comprehensive review. Microb. Cell Fact. 19, 1–42 (2020).

83. Mora, L. & Toldrá, F. Advanced enzymatic hydrolysis of food proteins for the production of bioactive peptides. Curr. Opin. Food Sci. 49, 100973 (2023).

84. Pleissner, D. & Lin, C. S. K. Valorisation of food waste in biotechnological processes. *Sustain*. Chem. Process. 1, 1–6 (2013).

85. Iqbal, H. M. N., Kyazze, G. & Keshavarz, T. Advances in the valorization of lignocellulosic materials by biotechnology: An overview. BioResources 8, 3157–3176 (2013).

86. Linger, J. G. et al. Lignin valorization through integrated biological funneling and chemical catalysis. Proc. Natl. Acad. Sci. 111, 12013–12018 (2014).

87. Borchert, A. J., Henson, W. R. & Beckham, G. T. Challenges and opportunities in biological funneling of heterogeneous and toxic substrates beyond lignin. Curr. Opin. Biotechnol. 73, 1–13 (2022).

88. Ragaert, K., Delva, L. & Van Geem, K. Mechanical and chemical recycling of solid plastic waste. Waste Management vol. 69 24–58 (2017).

89. Vora, N. et al. Leveling the cost and carbon footprint of circular polymers that are chemically recycled to monomer. Sci. Adv. 7, (2021).

90. Arnold, S., Moss, K., Henkel, M. & Hausmann, R. Biotechnological Perspectives of Pyrolysis Oil for a Bio-Based Economy. Trends Biotechnol. 35, 925–936 (2017).

91. Zakzeski, J., Bruijnincx, P. C. A., Jongerius, A. L. & Weckhuysen, B. M. The catalytic valorization of lignin for the production of renewable chemicals. Chem. Rev. 110, 3552–3599 (2010).

92. Xu, C., Nasrollahzadeh, M., Selva, M., Issaabadi, Z. & Luque, R. Waste-to-wealth: biowaste valorization into valuable bio(nano)materials. Chem. Soc. Rev. 48, 4791–4822 (2019).

93. Bhatt, M. et al. Valorization of solid waste using advanced thermo-chemical process: A review. J. Environ. Chem. Eng. 9, 105434 (2021).

94. Chen, C. C., Dai, L., Ma, L. & Guo, R. T. Enzymatic degradation of plant biomass and synthetic polymers. Nat. Rev. Chem. 4, 114–126 (2020).

