## Supplementary Information for "Complex waste stream valorisation through combined enzymatic hydrolysis and catabolic assimilation by *Pseudomonas putida*"

### Contents


| | |
|---|---|
| Table S1 | p2 |
| Table S2 | p3 |
| Table S3 | p7 |
| Table S4 | p10 |
| Table S5 | p11 |
| Table S6 | p12 |
| Table S7 | p13 |
| Table S8 | p14 |
| Table S9 | p15 |
| Table S10 | p16 |
| Table S11 | p17 |
| Figure S1 | p18 |
| Figure S2 | p19 |
| Figure S3 | P20 |
| Figure S4 | p21 |
| Figure S5 | p22 |
| Figure S6 | p23 |
| Figure S7 | p24 |
| Figure S8 | p25 |
| Figure S9 | p26 |
| Figure S10 | p27 |
| Figure S11 | p28 |
| Figure S12 | p29 |
| Figure S13 | p30 |
| Figure S14 | p31 |
| Figure S15 | p32 |
| Figure S16 | p33 |
| Figure S17 | p34 |
| Figure S18 | p35 |
| Figure S19 | p36 |
| Figure S20 | p37 |
| Figure S21 | p38 |
| Figure S22 | p39 |
| References | p40 |

**Table S1.** Plasmids used in this study.

| Name | Details |
|---|---|
| pK18mobsac-$P_{glcD}$::$P_{tac}$ | Replacement of the native *P. putida* KT2440 *glcD* promoter with the constitutive $P_{tac}$ promoter. |
| pK18mobsac-$P_{gcl}$::$P_{tac}$ | Replacement of the native *P. putida* KT2440 *gcl* promoter with the constitutive $P_{tac}$ promoter. |
| pK18mobsac-*gcd*::$P_{tac}$:*XylABE* | Integration of the *E. coli XylE*, *XylA*, and *XylB* genes, under the control of the constitutive $P_{tac}$ promoter, into the genome of *P. putida* KT2440 at the *gcd* locus for simultaneous gene deletion. |
| ptphKAB ($P_{proB}$:*pcaV* $P_{ppv}$:*TphK*, *TphA2*$_{II}$, *TphA3*$_{II}$, *TphB*$_{II}$, *TphA1*$_{II}$) | Operon for the import of TPA and intracellular conversion to PCA by *P. putida*. *TphK* from *P. mandelii*, *TphA2*$_{II}$, *TphA3*$_{II}$, *TphB*$_{II}$, and *TphA1*$_{II}$ from *Comamonas sp.* strain E6 were placed under the control of the *S. coelicolor* PCA inducible promoter $P_{ppv.}$ The *pcaV* transcriptional repressor from *S. coelicolor* was placed under the constitutive $P_{proB}$ promoter. |
| ptphKAB-*Nb1* | HEL4 nanobody gene with N-terminal His6 and TEV tag under the control of the constitutive $P_{pem7}$ promoter for *P. putida*. |
| ptphKAB-*IFNα2* | Interferon-α2 with N-terminal His6 and TEV tag under the control of the constitutive $P_{pem7}$ promoter for *P. putida*. |
| ptphKAB-UbiQ-*grk-ins* | Insulin analogue GRK with N-terminal His6, TEV tag and ubiquitin leader sequence under the control of the constitutive $P_{pem7}$ promoter for *P. putida*. |
| ptphKAB-UbiQl-*gekr-ins* | Insulin analogue GERK with N-terminal His6 and ubiquitin leader sequence under the control of the constitutive $P_{pem7}$ promoter for *P. putida*. |
| ptphKAB-Trxa-*gekr-ins* | Insulin analogue GERK with N-terminal His6, TEV tag and thioredoxin leader sequence under the control of the constitutive $P_{pem7}$ promoter for *P. putida*. |

**Table S2.** Sequence of genomic modifications.

| Name | Sequence 5'à3' | Notes |
|---|---|---|
| $P_{glcD}$::$P_{tac}$ | cgatacaaggccgcttcacccccagccatccatacgcatggtcgccctgaccaggcgctgctcttcc<br>tgctcgatctcccgcaaactgtcgcgtatgccgttgatatgctcacgcgccgcccgctgcgcctgctc<br>gggcagttgttccatgactgcgcggtacaaccgcgcatgctgccgatcaatctggcgtttctgcgct<br>ggccggcagtacaagttgttgaccgacgcaaacaccgtgctcaacgtcaggtcactgagcgattgc<br>agggtatgcaccagcactgggttatgcgacgcctcactgatggcccggtgaaaggcatggtcacg<br>gcgggcatgctcacgggcatcgagcgcctgcgcagcctcgtgcgcagccagcatttcttcgtagcg<br>ccggcggattagcaggcggtcgacgtcggtagcccgcaacgccgccagccgcgccgactcagcct<br>ccagcaacgcccggacctccagcaggtcgaacagagtgcgcggttgcgaactgaacaggtgcatc<br>aaaggcgtagcacccgcctgcccggtgagatcggcgacgaacgaaccccgcccctgttcggtgtc<br>gatgatgccgcgcccacgcagaacgcgcaggccctcgcgcaaggccgaacgtgaacagccaagc<br>ttttccaccagccgccgctccgacggcagtgcctggcccaccttgagcacgccttcgacgataagcc<br>gctcaacccgttcggccacctggtcggcgaccttggccttgccttcagtacccactgcgcacactcct<br>gctggtaggaccactttgacttatatcgccaatctagccagacagaaccgtgaatagacagtactg<br>cccgaagaaactggtaggaccagtcaacatctcactcgaccccaaacactagcacgcgcacgctg<br>catcggtgaccgcttcgccaacaacaacaaaaaccgttgcgagtgagcggcgcgccagacctaac<br>taatgctctatttaaatttgacaattaatcatcggctcgtataatgtgtggaattgtgagcggataac<br>aattgattaactttataaggaggaaaaacatatgaatatcctgtacgacgaacgcgtcgatggcgc<br>gctgcccaacgtggacctggccgccctgttgcaggcgctgcgcgatgccctgccggatcttgaaat<br>cctgcaccgcgatgaagacctcaaaccgtacgaatgcgacggcctgtcggcctaccgcaccgtgc<br>cactgctggtggcgctgcccgagcgcctggagcaggtgcagacgctgttgaagctttgccaccagc<br>gcggcgtaccggtggttgcgcgtggcgccggtaccggcctgtctggtggtgccctgccgctggcca<br>agggcatcctgctggtgatggcgcgcttcaaccgtatcctcgaggtcaacccgcaggggcgttacg<br>cccgcgtgcaaccgggcgtacgcaacctggccatctcccaggccgccgcaccccatggtatgtact<br>acgcacccgacccttcctcgcaaattgcctgctcgatcggtggcaacgtcgccgaaaacgccggtg<br>gcgtgcactgcctcaagtacggcctgaccgtacacaatgtgctcaaggtggacatccttacggtcg<br>agggcgaacgcctgagcctgggcagcgatgccctggacagccccggcttcgacctgctggcattgt<br>tcatcggctccgaaggcatgctcggtatcgtcaccgaagtcaccgtcaagctgctgcccaagcccc<br>aggtggcgcgggtgatactggccagtttcgacagcgtcgaggacgccggccgggcagtcgccgac<br>atcatcgctgccggcatcattcccggcggcctggagatgatggacaacctggcgatccgcgccgct<br>gaagacttcatccatgccggctacccggtggacgcggcggcgatcctgctgtgcgaactggatggc<br>gtggaagccgatgtttacgacgactgcgagcgcgtcgccgccgtgctgacgcaagccggggcccg<br>cgaggtgcacctggcctgcgacgaagccgagcgcgctcgtttctgggccgggcgcaagaacgcct<br>tcccggccgtgggacgcatctccccggactactac | Upstream and downstream homology arms of *glcD*, $P_{tac}$ promoter, RBS |
| $P_{gcl}$::$P_{tac}$ | cccagacctatgccgacttcgactggcagatttctgcggtgaccggtaaggccaagctggccgaca<br>gtgattatgacgcggcggccgagaccatcattcgtatggtcttgaaaggttgtgagcctgaggcag<br>cctgaaccatctcgaagggccattcgcgggcaagcccgctcccacaggatcaccacatgcctgaa<br>ggttgtggagtccctgtgggagcgggcttgcccgcgaatgggccagaacaggcaatagagattta<br>ggccgctaccccggcatctgcccgcagccccacctcctcgatcgcactgatcgcacactgctcgtcg<br>atatccgacgtatcgccgctgatccccacggcacccagcaccttgccgtcctgatcacgaatcagc<br>accccacccggcgccggcaccaccggccgctcacccatcccgttcaacgcggcaaaaaaagccg<br>gccgctgctgcgcatccagcgccagcaagcgcgagcccttgcccagggcaatcgcaccccaggct<br>ttgccagtggccacttcaggccgaatcaggctagcgccatcctcgcgctgcagcgccagcaggtgc<br>ccgccggcatccagcaccgccacggtcagcggcgcggcattgatcttgcggcccgccaccagtgc<br>ggcattcaccaggctgacggcgactttcaggttaagaacgttcatggaaaggtcctcttcttgttgt<br>gagaagccctaagggcagttgaaaaccgatcgaaaaaatagaacacaacgagcgatatttttgta<br>tacaatattttgaaacgatcgtatgcgatggcgcaaacctccgttttcacgggcgctctgacgaaag<br>ccagcttagccgatgaaaacccattgacctaagccgtcaggcgtgaatacactctgtcgcaaagca<br>agttgtggcgcgccagacctaactaatgctctatttaaatttgacaattaatcatcggctcgtataat<br>gtgtggaattgtgagcggataacaattgattaactttataaggaggaaaaacatatgagcaaaat | Upstream and downstream homology arms of *gcl*, $P_{tac}$ promoter, RBS |

| | | |
|---|---|---|
| | gagagcaatcgatgcagccgttctggtcatgcgccgtgaaggtgtagataccgcgttcggcatccc<br>gggggctgccatcaacccgttgtattcggccctgaaaaaagtcggtggcatcgatcacgtcctcgct<br>cgtcacgtcgaaggtgcctcgcacatggccgagggctacacccgcgccaacccgggcaacatcgg<br>tgtgtgcatcggcacttccggccctgccggcaccgacatggtcaccggcctgtacagtgcctcggcc<br>gactccatcccgattctgtgcatcactggccaggcgccacgtgcccgtctgcacaaggaagacttcc<br>aggctgtcgacatcaccaacatcgtcaagccagtgaccaagtgggcgaccaccgttctggagcca<br>ggccaggtgccttacgccttccagaaggccttctatgaaatgcgtaccggccgcccaggcccggtg<br>ctgatcgacctgccgttcgacgtgcagatggccgaaatcgaattcgacatcgacgcctacgaaccg<br>ctgcccgtgcacaaaccgtccgccacacgcgtacaggccgaaaaagccctggccctgctcaatga<br>cgccgagcgcccactgctggtagccggtggcggcatcatcaacgccgacgccagtgacaagctgg<br>tcgaattcgccgaactgaccggcgtacccgtgatcccgaccctgatgggctggggcaccatcccgg<br>acgaccacgcacagatggtcggcatggtcggcctgcagacctcgcaccgctatggcaacgcaacc<br>ctgctgaaatccgacctggtgttcggtatcggtaaccgctgggccaaccgccacaccggttccgtcg<br>atgtctacaccgaaggccgcaagttcgtgcacgtcgacatcgaaccgacccagatcggccgcgtgt<br>tcaccccggacctgggcatcgtttccgatgctggtaaggc | |
| *Δgcd*::P$_{tac}$*XylABE* | ttggtgaaggcgaggacttcaacagcttcccttgcgacttccagaacctggctttctgcggctcgca<br>ggtgggcaactgggtgggtggcatctggtacaactggccagtcagccagtgggccctgcgcgtgc<br>gctacaaccttaccccgagctgtacgcccaggtcggcgtcttcgagcagaacccttccaacctcg<br>aatccggcaatggtttcaagctcagcggcagcggcacccagggtgcggtaatgccgttcgaactg<br>gtatggaccccacgtatccaaggcttgaaaggggaatatcgtgccggctactactacagtaatgcc<br>aaggcacaagatgttctcaaggacagcaacggtcagccggccgccctcagcggcgccgcctaccg<br>cagcagttcgagcaagcacggcttgtggattggcgcccagcagcaggtcacctcgctggcgtccga<br>ccagtcgcgcggcttgagcgtgttcgccaacgccacggtgcatgacaaaaagaccaatgccatcg<br>acaactatgtgcaggcaggactggtattcaaagggcctttcgatgcccgcgccaaggacgacatc<br>ggtttcgccctggcccgcgtgcacgtcaaccctgcctatcgcaagaacgcccgcctggtcaaccag<br>gccgccggcctctatgactacgacaacccgggcttcctgccagtgcaggacaccgagtacagcgc<br>cgagctgtattacggcattcacttggccgactggctcacggtacgccccaacctgcagtacatccgc<br>cacccgggcggggtgtcgcaggtcgatggcgccctgatcggcggcctgaagatccagagcagttt<br>ctaaccccttttttgacctgacggagaacctacgttgacaattaatcatcggctcgtataatgtgtgg<br>aattgtgagcggataacaattgattaactttataaggaggaaaaacatatgaatacccagtataat<br>tccagttatatattttcgattaccttagtcgctacattaggtggtttattatttggctacgacaccgcc<br>gttatttccggtactgttgagtcactcaataccgtctttgttgctccacaaaacttaagtgaatccgct<br>gccaactccctgttagggttttgcgtggccagcgctctgattggttgcatcatcggcggtgccctcgg<br>tggttattgcagtaaccgcttcggtcgtcgtgattcacttaagattgctgctgtcctgtttttatttctg<br>gtgtaggttctgcctggccagaacttggttttacctctataaacccggacaacacagtgcctgtttat<br>ctggcaggttatgtcccggaatttgttatttatcgcattattggcggtattggcgttggtttagcctca<br>atgctctcgccaatgtatattgcggaactggctccagctcatattcgcgggaaactggtctcttttaa<br>ccagtttgcgattattttcgggcaacttttagtttactgcgtaaactattttattgcccgttccggtgat<br>gccagctggctgaatactgacggctggcgttatatgtttgcctcggaatgtatccctgcactgctgtt<br>cttaatgctgctgtataccgtgccagaaagtcctcgctggctgatgtcgcgcggcaagcaagaaca<br>ggcggaaggtatcctgcgcaaaattatgggcaacacgcttgcaactcaggcagtacaggaaatta<br>aacactccctggatcatggccgcaaaaccggtggtcgtctgctgatgtttggcgtgggcgtgattgt<br>aatcggcgtaatgctctccatcttccagcaatttgtcggcatcaatgtggtgctgtactacgcgccgg<br>aagtgttcaaaacgctgggggccagcacggatatcgcgctgttgcagaccattattgtcggagtta<br>tcaacctcaccttcaccgttctggcaattatgacggtggataaatttggtcgtaagccactgcaaatt<br>atcggcgcactcggaatggcaatcggtatgtttagcctcggtaccgcgttttacactcaggcaccgg<br>gtattgtggcgctactgtcgatgctgttctatgttgccgcctttgccatgtcctggggtccggtatgct<br>gggtactgctgtcggaaatcttcccgaatgctattcgtggtaaagcgctggcaatcgcggtggcgg<br>cccagtggctggcgaactacttcgtctcctggaccttcccgatgatggacaaaaactcctggctggt<br>ggcccatttccacaacggtttctcctactggatttacggttgtatgggcgttctggcagcactgtttat<br>gtggaaatttgtcccggaaaccaaaggtaaaacccttgaggagctggaagcgctctgggaaccgg<br>aaacgaagaaaacacaacaaactgctacgctgtaatttaactttaagaaggagatatacatatgc | Upstream and downstream homology arms of *gcd*, P$_{tac}$ promoter, RBS, *XylE* gene, *XylA* gene, *XylB* gene, rrnC terminator. |

aagcctattttgaccagctcgatcgcgttcgttatgaaggctcaaaatcctcaaacccgttagcattc
cgtcactacaatcccgacgaactggtgttgggtaagcgtatggaagagcacttgcgttttgccgcct
gctactggcacaccttctgctggaacggggcggatatgtttggtgtgggggcgtttaatcgtccgtg
gcagcagcctggtgaggcactggcgttggcgaagcgtaaagcagatgtcgcatttgagtttttcca
caagttacatgtgccattttattgcttccacgatgtggatgtttcccctgagggcgcgtcgttaaaag
agtacatcaataattttgcgcaaatggttgatgtcctggcaggcaagcaagaagagagcggcgtg
aagctgctgtggggaaccgccaactgctttacaaaccctcgctacggcgcgggtgcggcgacgaa
cccagatcctgaagtcttcagctgggcggcaacgcaagttgttacagcgatggaagcaacccata
aattgggcggtgaaaactatgtcctgtggggcggtcgtgaaggttacgaaacgctgttaaataccg
acttgcgtcaggagcgtgaacaactgggccgctttatgcagatggtggttgagcataaacataaaa
tcggtttccagggcacgttgcttatcgaaccgaaaccgcaagaaccgaccaaacatcaatatgatt
acgatgccgcgacggtctatggcttcctgaaacagtttggtctggaaaaagagattaaactgaaca
ttgaagctaaccacgcgacgctggcaggtcactctttccatcatgaaatagccaccgccattgcgct
tggcctgttcggttctgtcgacgccaaccgtggcgatgcgcaactgggctgggacaccgaccagtt
cccgaacagtgtggaagagaatgcgctggtgatgtatgaaattctcaaagcaggcggtttcaccac
cggtggtctgaacttcgatgccaaagtacgtcgtcaaagtactgataaatatgatctgttttacggtc
atatcggcgcgatggatacgatggcactggcgctgaaaattgcagcgcgcatgattgaagatggc
gagctggataaacgcatcgcgcagcgttattccggctggaatagcgaattgggccagcaaatcctg
aaaggccaaatgtcactggcagatttagccaaatatgctcaggaacataatttgtctccggtgcatc
agagtggtcgccaggagcaactggaaaatctggtaaatcattatctgttcgacaaataacggctaa
ctgtgcagtccgttggcccggttatcggtagcgataccgggcatttttttaaggaacgatcgatatgt
atatcgggatagatcttggcacctcgggcgtaaaagttattttgctcaacgagcagggtgaggtgg
ttgcttcgcaaacggaaaagctgaccgtttcgcgcccgcatccactctggtcggaacaagacccgg
aacagtggtggcaggcaactgatcgcgcaatgaaagctctgggcgatcagcattctctgcaggac
gttaaagcattgggtattgccggccagatgcatggagcaaccttactggatgctcaacaacgggta
ttgcgccctgccattttgtggaacgacgggcgctgtgcgcaagagtgcactttgctggaagcgaga
gttccgcaatcacgagtgattaccggcaacctgatgatgcccggatttactgcgcctaaattgctat
gggttcagcggcatgagccggagatattccgtcaaatcgacaaagtattattaccgaaagattact
tgcgtctgcgtatgacgggggagtttgccagcgatatgtctgacgcagctggcaccatgtggctgg
atgtcgcaaagcgtgactggagtgacgtcatgctgcaggcttgcgacttatctcgtgaccagatgcc
cgcattatacgaaggcagcgaaattactggtgctttgttacctgaagttgcgaaagcgtggggtat
ggcgacggtgccagttgtcgcaggcggtggcgacaatgcagctggtgcagttggtgtgggaatggt
tgatgctaatcaggcaatgttatcgctggggacgtcggggtctattttgctgtcagcgaagggttct
taagcaagccagaaagcgccgtacatagcttttgccatgcgctaccgcaacgttggcatttaatgtc
tgtgatgctgagtgcagcgtcgtgtctggattgggccgcgaaattaaccggcctgagcaatgtccc
agctttaatcgctgcagctcaacaggctgatgaaagtgccgagccagtttggtttctgccttatcttt
ccggcgagcgtacgccacacaataatccccaggcgaaggggttttctttggtttgactcatcaac
atggccccaatgaactggcgcgagcagtgctggaaggcgtgggttatgcgctggcagatggcatg
gatgtcgtgcatgcctgcggtattaaaccgcaaagtgttacgttgattgggggcggggcgcgtagt
gagtactggcgtcagatgctggcggatatcagcggtcagcagctcgattaccgtacgggaggga
tgtggggccagcactgggcgcagcaaggctggcgcagatcgcggcgaatccagagaaatcgctc
attgaattgttgccgcaactaccgttagaacagtcgcatctaccagatgcgcagcgttatgccgctt
atcagccacgacgagaaacgttccgtcgcctctatcagcaacttctgccattaatggcgtaacaga
taaaaaaaatccttagctttcgctaaggatgtcgctgggcaccaagatgggcgattacatcattgct
tacaaattagccgagtaagcgacaccgctcccgcaggtgcgtttgagcatgggcacgcagtcgag
cacttcctgcaactgattgccggtggccgaggtgagcgcggcggcgaagtcggtgagcggcgaag
ggtcgttggggctgaagcggttgaggtagttcttcagctcttcgctgtacagcgggttgagcggcag
tagctctttgatcgcgttgatcagcgccatgccgtaggctttgacctcgtcggtcggctcggcgggct
ggcgcgggtattcgacttcgaccaggtagggcgggcggtggtgcttgagccagttgcggatgcgca
cccgggacaggccctgggcgacgaactgcaatttgccgttttcgcggctggcatggtgcaccttga
ccagcgtgccgtactgcggcagtgccgaggtgtcgaagtggcggtggtcttctggcggggtgtcca
tgaagaacagcgccaggcagtggtccggtgacttggctaccaggtcgagcgtttcggcccagggct

| | |
|---|---|
| | cttcattgacgatcaccggcagtacctgggccgggaagaacgggcggttgtggatcgggatgacg<br>tagaccttgtccggcagctgttgaccgggcagggcgagggcgtggctgacttcggcctggagggtg<br>tgttcgacttcgctgtgttcgtcagggtgttcggggaaatcctgctggtcgctcatggggcacctgcg<br>tgaatgactatggtgcttagatggggcggggagggatgggtttcaatggttcgggagcagccttgt<br>gtcgcgattgggctgcgcagcagccccagaatccatccagcattctagatcgttggggccgctgcg<br>cgaccgaaacgcgacacaaggccgctccctcagggttgag |

**Table S3.** Sequence of plasmid expressed genes.

| Name | Sequence 5’à3’ | Notes |
|---|---|---|
| $P_{proB}$:*pcaV*<br>$P_{ppv}$:*TphK*,<br>*TphA2*$_{II}$,<br>*TphA3*$_{II}$,<br>*TphB*$_{II}$, *TphA1*$_{II}$ | atttgtcctactcaggagagcgttcaccgacaaacaacagataaaacgaaaggcccagtctttcgactgagcc<br>tttcgttttatttgatgcctttaattaaagcggataacaatttcacacaggaggccgcctaggcctcaacccggt<br>gcaactgccggttcagccggattacgcaggccttctgctgcatctgcaacacgacgaatcagatcaaaaaaa<br>ctgcctgttcatctgctgccagcggtgccagaaaaacctgattcatacgtgcaatacgaacacccagacgacg<br>atgaacacgcagaccttcatcggtcagacgcagcagtgaacgacgaccatcctgcggatcacgaactttatcc<br>agcagaccacgacgacccagacggctaacaacttctgcaatggtgctacgatccaggccaacacgttcacca<br>acggtacgctgatccagacccggttctgcaaccagtgcattcagaactgcatactgcggactggtggtttcttcg<br>ctaaccattgtattccacagcagataatgtgcctgttgcagacgacgtgccagatgacccggatgggttgccag<br>atcaactgctgccatttgtattacctccttatttcaaagttaaacaaaattatttgtagagggaaaccgttgtggt<br>ctccctgaatatatcgtacgagccttatgcatgcccgtaaagttatccagcaaccactcatagacctagggcag<br>cagatagggacgacgtggtgttagctgtgggcaaaaaacattatccagaacgggagtgcgccttgagcgaca<br>cgaattatgcagtgatttacgacctgcacagccataccacagcttccgatggctgcctgacgccagaagcattg<br>gtgcaccgtgcagtcgatgataagctgtcaaacgcatgcaaaatttatcaaaaagagtgttgactatactcagt<br>gccctgactatgatacttagattcatactcagtgccctgactattttaactttaagaaggagatatacatatgctc<br>ccggagtccaagccgctgcagctcccacagtttgtcgatgagcgtcgcgtgtccccttttcaatatggggtcatc<br>atcctgtgtggcttggtcatgttcctggatggcttcgatacccaagcaattagctatatggcgccttatatcgcgc<br>aggaatggtccctctcgaagcagatgttgggcccaattttctccagctccttggctggtctgatggtcgggtacc<br>tcgtcctgacgcctttgtcggagcgttttgggcataagaaggccattattgtgtcgacgatcgtgttttccctctg<br>caccctcgcttcggtgtgggcaacgaacgtcacggaattgatggtcttgcggttcattacgggtatcgggttggg<br>cacggccgccccttcggcaatcgcaatgaccggtgagtactccccgaagcggctgcgcgctaccttcgtcctgg<br>caatctattgcggtttctccctgggcttcatcgcagcagggttggccgcaggctggtttatcccgcattatggttg<br>gcggtcgatgttttgggtcggtgctctggccccgctgatcttggtgcccgtcctgtttcgttttctcccagagtcga<br>tggtgtttatgatcaaaaataaggtccccccacaacaaatcttgaacgtctttcgtaagatcgatgcaagcgtct<br>cgcagcagagcaaacctgtcttcgtggtcgaagcactcgaacagggccaacacggcgctctcgcgagcctgtt<br>cacccgtgaccgcatcatgggtacggtgctgttgtggttggtgttcgcaatcaacctgggtgaattttatgccctc<br>cagtcgtggctgccgagcatcatgacggggttgaattacccgatgagcacggtcgtcatggcgaccaccttga<br>ccacggtcgggggtattgcggcagctttcattacggggccaagcatggaccgcttgggtgcctataaaaccgtg<br>ggtattctctacttggtcggcttcgtctttgtggccctgaccggcatggcatttaactcgccgttgtgggtcttgtt<br>gagcgctaattttctggctgggtgctgcatttcgggtggtcagaaaagcctcattgcactcgctgcagtgttcta<br>cccggcgccaatgcggtccaccggtgtcggttgggctctgggtattggccggatcggcgggatcgcggggcct<br>attgtcgtgggcgcagctttgggtattggctggtcgccgagcgcggtgttttacgggatggcaatcccaatgctg<br>attgccggtttgatcgtgctctttctcggtcgccgctatgggaactccgataaggtgtaactcttaggaaccaac<br>atgcaagaatccatcatccagtggcatggggccactaatacgcgcgtgccttttggtatctataccgacacagc<br>caatgctgatcaggaacagcagcgcatctatcgcggcgaggtctggaactacttgtgcctggaatctgaaattc<br>ccggggccggtgatttccgcactacctttgccggtgaaacaccgatagttgtcgtacgggatgccgaccagga<br>aatctacgccttcgagaaccgctgcgcgcatcgcggcgctctcatcgctctggagaaatcgggccgtacggat<br>agtttccagtgcgtctatcacgcctggagctacaaccgacagggagatctgaccggcgttgccttcgagaaag<br>gtgtcaagggccagggtggcatgccggcctcattctgcaaagaagagcatggcccgcgcaagctccgcgtgg<br>ctgtcttttgcggtttggtctttggcagtttttccgaggacgtgcccagcattgaggattaccttggccctgagatt<br>tgcgagcgcatagagcgcgtgctgcacaagcccgtagaagtcatcggtcgcttcacgcaaaagctgcctaaca<br>actggaagctctacttcgagaacgtgaaggacagctatcacgccagcctcctgcatatgttcttcaccaccttcg<br>agctgaatcgcctctcacaaaaaggcggtgtcatcgtcgacgagtcgggtggccaccatgtgagctattccatg<br>atcgatcgcggcgccaaagacgactcgtacaaggaccaggccatccgctccgacaacgagcgttaccggctc<br>aaagatcctagccttctagagggctttgaggagttcgaggacggcgtgaccctgcagatcctttctgtgttccct<br>ggctttgtgctgcagcagattcagaacagcatcgccgtgcgtcagttgctgcccaagagcatctccagctcgga<br>actcaactggacctatcttggctatgcagatgacagtgccgagcaacgcaaggtcagactcaaacaggccaa<br>ccttatcggcccggccggattcatttccatggaagacggagctgtcggtggattcgtgcagcgtggcatcgcag<br>gcgctgccaaccttgatgcagtcatcgagatgggcggagaccacgaaggctctagcgagggccgcgccacgg<br>aaacctcggtacgcggcttttggaaggcctaccgcaagcatatgggacaggagatgcaagcatgatcaatga<br>aattcaaatcgcggccttcaatgccgcctacgcgaagaccatagacagtgatgcaatggagcaatggccaac<br>ctttttcaccaaggattgccactattgcgtcaccaatgtcgacaaccatgatgagggacttgctgccggcattgt<br>ctgggcggattcgcaggacatgctcaccgaccgaatttctgcgctgcgcgaagccaatatctacgagcgccac<br>cgctatcgccatatcctgggtctgccttcgatccagtcaggcgatgcaacacaggccagcgcttccactccgttc<br>atggtgctgcgcatcatgcatacaggggaaacagaggtctttgccagcggtgagtacctcgacaaattcacca<br>cgatcgatggcaagttacgtctgcaagaacgcatcgcggtttgcgacagcacggtgacggacacgctgatggc | rrnB T1 terminator,<br><br>pcaV transcriptional repressor (3’à5’ sequence),<br><br>RBS and<br><br>$P_{ProB}$ promoter, native *Comamonas* sp.<br><br>$P_{ppv}$ promoter,<br><br>TphK gene,<br><br>TphA2$_{II}$ gene,<br><br>TphA3$_{II}$ gene (underlined sequence shows the gene overlap with TphA2$_{II}$),<br><br>TphB$_{II}$ gene (underlined sequence shows the gene overlap with TphA3$_{II}$),<br><br>TphA1$_{II}$ gene,<br><br>rrnC terminator |

| | | |
|---|---|---|
| | attgccgctatgacaatagtgcaccgtagattggctttggccatcggcgatccccacggtattggcccagaaat<br>cgcactgaaagctctccagcagctgtctgtcaccgaaaggtctcttatcaaggtctatggaccttggagcgctct<br>cgagcaagccgcacgggtttgcgaaatggagccgcttcttcaagacatcgttcacgaggaagccggcacactt<br>acacaaccagttcaatggggagaaatcaccccgcaggctggtctatctacggtacaatccgcaacagcggcta<br>tccgagcgtgcgaaaacggcgaagtcgatgccgtcattgcctgccctcaccatgaaacggccattcaccgcgc<br>aggcatagcgttcagcggctacccatctttgctcgccaatgttcttggcatgaacgaagaccaggtattcctgat<br>gctggtgggggctggcctgcgcatagtgcatgtcactttgcatgaaagcgtgcgcagcgcattggagcggctct<br>ctcctcagttggtggtcaacgcggcgcaggctgccgtgcagacatgcaccttactcggagtgcctaaaccaaa<br>agtcgctgtattcgggatcaaccctcatgcatctgaaggacagttgttcggcctggaggactcccagatcaccg<br>ttcccgctgtcgagacactgcgcaagcgcggcctagcagtagacggccccatgggagctgacatggttctggc<br>acagcgcaagcacgacctgtatgtagccatgctgcacgatcagggccatatccccatcaagctgctggcacct<br>aacggagccagcgcactatctatcggtggcagggtggtgctttccagcgtgggccatggcagcgccatggaca<br>ttgccggccgtggcgtggctgacgcaacggcactcctacgcacaatagccctactcggagcccaaccggtctg<br>aggactctctatgaaccaccagatccatatccacgactccgatatcgcgttcccctgcgcgcccgggcaatccg<br>tactggatgcagctctgcaggccggcatcgagctgccctattcctgccgcaaaggtagctgtggcaactgtgcg<br>agtacgctgctcgacggaaatattgcctccttcaatggcatggccgtgcgaaacgaactctgcgcctcggaac<br>aagtgctgctgtgcggctgcactgcagccagcgatatacgtatccacccgagctcctttcgccgtctcgacccg<br>gaagcccgaaaacgttttacggccaaggtgtacagcaatacactggcggcacccgatgtctcgctgctgcgcc<br>tgcgcctgcctgtgggcaagcgcgccaaatttgaagccggccaatacctgctgattcacctcgacgacgggga<br>aagccgcagctactctatggccaatccaccccatgagagcgatggcatcacattgcatgtcaggcatgtacctg<br>gtggtcgcttcagcactatcgttcagcagttgaagtctggtgacacattggatatcgaactgccattcggcagca<br>tcgcactgaagcctgatgacgcaaggcccctgatttgcgttgcgggtggcacgggatttgcgcccattaaatcc<br>gttcttgatgacttagccaaacgcaaggttcagcgcgacatcacgctgatctgggggctcgcaacccctcggg<br>cctgtatcttcctagcgccatcgacaagtggcgcaaagtctggccacagtttcgctacattgcagccatcaccg<br>acctaggcgatatgcctgcggatgctcacgcaggtcgggtggatgacgcgctacgcactcactttggcaacctg<br>cacgatcatgtggtgcactgctgtggctcaccagctctggttcaatcagtgcgcacagccgcttccgatatgggc<br>ctgcttgcacaggacttccacgcggatgtttttgcgacaggcccgactggtcaccactagcagataaaaaaat<br>ccttagctttcgctaaggatg | |
| $P_{pem7}$:*Nb1* | tgttgacaattaatcatcggcatagtatatcggcatagtataatacgacaaggtgaggaactaaaccgattaa<br>ctttataaggaggaaaaacatatgcaccatcaccaccatcacggtgcaggggagaacctgtactttcagtccg<br>ctggggcagaagtgcaactgctggagagcggtggtggttccgtccaaccaggcggtagcctccggctcagctg<br>tgccgctagcggctttcgtattagcgacgaggacatgggctggtttcgccaggccccagggaaggaacgcgaa<br>atcgtcagctcgatttatggcccttcggggtccacgtattacgcggattccgtgaagggtcgcttcaccatttccc<br>gggataatagcaagaatacgctctacctccagatgaactccctgcgcgcggaagacacggctgtgtattattgt<br>gccagcgcattggaacctctgtcggaaccactgggttttcgcggccagggcaccttggtgaccgtgtcgtccta<br>aggtgactgggaaaaccctggcgactagtcttggactcctgttgatagatccagtaatgacctcagaactccat<br>ctggatttgttcagaacgctcggttgccgccgggcgtttttattggtgagaat | $P_{pem7}$ promoter,<br><br>RBS, His6-TEV-*Nb1* gene, lambda T0 terminator |
| $P_{pem7}$: *IFNα2* | tgttgacaattaatcatcggcatagtatatcggcatagtataatacgacaaggtgaggaactaaaccgattaa<br>ctttataaggaggaaaaacatatgcaccatcaccaccatcacggtgcaggggagaacctgtactttcagtccg<br>ctggggcatgtgatctgccgcagacccatagcctgggtagccgtcgtaccctgatgctgctggcacagatgcgt<br>cgtattagcctgtttagctgtctgaaagatcgtcacgattttggttttccgcaagaagaatttggcaaccagtttc<br>agaaagcagaaaccattccggttctgcatgaaatgattcagcagatctttaacctgttcagcaccaaagatag<br>cagcgcagcatgggatgaaaccctgctggataaattctataccgaactgtatcagcagctgaatgatctggaa<br>gcatgtgttattcagggtgttggtgttaccgaaacaccgctgatgaaagaagatagcattctggcagttcgcaa<br>atatttccagcgtattaccctgtacctgaaagagaaaaaatacagcccgtgtgcatgggaagttgttcgtgcag<br>aaattatgcgtagctttagcctgagcaccaatctgcaagaaagcctgcgtagcaaagaataggtgactgggaa<br>aaccctggcgactagtcttggactcctgttgatagatccagtaatgacctcagaactccatctggatttgttcag<br>aacgctcggttgccgccgggcgtttttattggtgagaat | $P_{pem7}$ promoter,<br><br>RBS, His6-TEV-*IFNα2*gene, lambda T0 terminator |
| $P_{pem7}$:ubiQ-*grk-ins* | tgttgacaattaatcatcggcatagtatatcggcatagtataatacgacaaggtgaggaactaaaccgattaa<br>ctttataaggaggaaaaacatatgcaccatcaccaccatcacggtgcaggggagaacctgtactttcagtccg<br>ctggggcacagattttcgtcaagacgctcaccgggaagaccattaccctggaagtggaatccagcgataccat<br>cgacaatgtgaaaagcaaaatccaagacaaagaaggtatcccgccggatcagcaggcactcatctttgcggg<br>taaacaactcgaagatggcgccacgctgtccgattataatatccaaaaggaatccacgttgcatctcgtcttgg<br>cgctcgcaggggggcgcttcgtcaatcaacatttgtgtgggagccatttggtcgaagccttgtatttggtctgcg<br>gggaacgcgggttcttctatacgccgaagaccaaacgggggattgtggagcaatgctgtacgtccatttgttcg | $P_{pem7}$ promoter,<br><br>RBS, His6-TEV-*gkr-ins* insulin analogue gene with ubiQ leader, lambda T0 terminator |

| | | |
|---|---|---|
| | ctctatcagctcgaaaactattgtaatggctaagactcctgttgatagatccagtaatgacctcagaactccatctggatttgttcagaacgctcggttgccgccgggcgttttttattggtgagaat | |
| $P_{pem7}$:ubiQ-*gekr-ins* | tgttgacaattaatcatcggcatagtatatcggcatagtataatacgacaaggtgaggaactaaaccgattaactttataaggaggaaaaacatatgcaccatcaccaccatcacggtgcaggggagaacctgtactttcagtccgctggggcacagattttcgtcaagacgctcaccgggaagaccattaccctggaagtggaatccagcgataccatcgacaatgtgaaaagcaaaatccaagacaaagaaggtatcccgccggatcagcaggcactcatctttgcgggtaaacaactcgaagatggcgccacgctgtccgattataatatccaaaaggaatccacgttgcatctcgtcttggcgctcgcagggggggcgcttcgtcgagcaacatttgtgtggtagccacttggtggaggcgctctacctggtctgtggtgagcgtggctttttttatacccccaaaacgaaacgtgggattgtcgaacaatgctgtacctcgatctgtagcctgtatcaactggaaaactactgcaatggttaagactcctgttgatagatccagtaatgacctcagaactccatctggatttgttcagaacgctcggttgccgccgggcgttttttattggtgagaat | $P_{pem7}$ promoter,<br><br>RBS, His6-*gekr-ins* insulin analogue gene with ubiQ fusion tag, lambda T0 terminator |
| $P_{pem7}$:Trxa-*gekr-ins* | tgttgacaattaatcatcggcatagtatatcggcatagtataatacgacaaggtgaggaactaaaccgattaactttataaggaggaaaaacatatgcaccatcaccaccatcacggtgcaggggagaacctgtactttcagtccgctggggcaagcagcgatctgatcaaacatgtcaccgacgcctccttcgaagccgatgtcctgaaggccgaaggcgcggtactggtcgactactgggctgaatggtgcggtccatgcaagatgatcgctccggttctggacgacatcgcttccacctacgagggcaaactgaccgtcgccaagctgaacatcgacgagaaccaggaaaccccggccaagcacggtgtgcgtggtatcccgacgctgatgctgttcaagaacggcaacgtcgaagccaccaaggttggcgcactgtccaaatcgcagctggccgcattcctcgacgcccacctgggcggtggcggcgggcgtttcgtcgagcaacatttgtgtggtagccacttggtggaggcgctctacctggtctgtggtgagcgtggctttttttatacccccaaaacgaaacgtgggattgtcgaacaatgctgtacctcgatctgtagcctgtatcaactggaaaactactgcaatggttaagactcctgttgatagatccagtaatgacctcagaactccatctggatttgttcagaacgctcggttgccgccgggcgttttttattggtgagaat | $P_{pem7}$ promoter,<br><br>RBS, His6-TEV-*gekr-ins* insulin analogue gene with TrxA leader sequence, lambda T0 terminator |

**Table S4.** Oligonucleotides used in this work.

| Name | Sequence (5'à3') |
|---|---|
| Therapopron | gcccgactggtcaccactagcagataaaaaaaatccttagctttcg |
| Therapopron | gccagggttttcccagtcacttagccattacaatagttttcga |
| GA50 | ctaaggattttttttatctgctagtggtgaccagtcggg |
| GA51 | aaaactattgtaatggctaagtgactgggaaaaccctgg |
| AB242 | atgaaccaccagatccatatccac |
| GA275 | cagataaaaaaaatccttagctttcg |
| GA209 | gagaacctgtactttcagtccgctggggcatgtgatctgccgcagacc |
| GA206 | atgccccagcggactgaaagta |
| GA210 | aagactagtcgccagggttttcccagtcacctattctttgctacgcaggcttt |
| KAB101 | tcaacccggtgcaactgc |
| KAB102 | ccggctgaaccggcagttgcaccgggttgaggcctaggcggcctcct |
| KAB103 | gtttaactttgaaataaggaggtaatacaaatggcagcagttgatctggc |
| KAB104 | ttgtattacctccttatttcaaagttaaacaaaat |
| KAB105 | tcaacactctttttgataaattttgcatgcgtttgacagcttatcatcgactgc |
| KAB106 | ctattttaactttaagaaggagatatacatatgctcccggagtccaag |
| KAB107 | gatggattcttgcatgttggttcctaagagttacaccttatcggagttcccatag |
| KAB108 | cgccgctatgggaactccgataaggtgtaactcttaggaaccaacatgcaaga |
| GA268 | agtgagcgcaacgcaattaatgtgagttagtgcgtttgagcatgggcac |
| GA278 | taatgcagctggcacgac |
| GA276 | ttagcgaaagctaaggattttttttatctgttacgccattaatggcagaag |
| GA275 | cagataaaaaaaatccttagctttcg |
| GA267 | aaaaaaaatccttagctttcgctaaggatgtcgctgggcaccaagatg |
| GA265 | tgctacgctgtaatttaactttaagaaggagatatacatatgcaagcctattttgaccag |
| GA263 | gcatatgtatatctccttcttaaagttaaattacagcgtagcagtttgttg |
| GA262 | attgattaactttataaggaggaaaaacatatgaatacccagtataattccag |
| GA272 | tatataactggaattatactgggtattcatatgtttttcctccttataaagttaatc |
| GA271 | cccctttttgacctgacggagaacctacgttgacaattaatcatcggctcg |
| GA269 | cgtaggttctccgtcaggtc |
| GA270 | ccctgagtgcttgcggcagcgtgaagctagttggtgaaggcgaggacttc |
| GA277 | ctttctacgtgttccgcttc |
| AB238 | atgcaagaatccatcatccagtg |
| AB257 | ctagcttcacgctgccgcaag |
| AB258 | ctaactcacattaattgcgttgcgctcactg |
| AB 259 | agtgagcgcaacgcaattaatgtgagttagcccagacctatgccgacttc |
| AB 260 | ttccacacattatacgagccgatgattaattgtcaaatttaaatagagcattagttaggtctggcgcgccacaacttgctttgcgaca |
| AB 261 | attaatcatcggctcgtataatgtgtggaattgtgagcggataacaattgattaactttataaggaggaaaaacatatgagcaaaat |
| AB 262 | ccctgagtgcttgcggcagcgtgaagctaggccttaccagcatcggaaac |
| AB 263 | agtgagcgcaacgcaattaatgtgagttagcgatacaaggccgcttcac |
| AB 264 | ttccacacattatacgagccgatgattaattgtcaaatttaaatagagcattagttaggtctggcgcgccgctcactcgcaacggtttt |
| AB 265 | attaatcatcggctcgtataatgtgtggaattgtgagcggataacaattgattaactttataaggaggaaaaacatatgaatatcct |

**Table S5.** Representation waste feedstocks, chemicals and enzymes used in this study.

| |
|---|
| ***Municipal waste composition*** |
| 3.0-5.0 mm Poly Lactic Acid beads (Goodfellow, ME346310) |
| 5.0µm Polyethylene Terephthalic acid film (Goodfellow, ES301005) |
| Micronised Beetroot (Biopower Technology Limited) |
| Micronised Apple (Biopower Technology Limited) |
| Micronised Wheat straw (Biopower Technology Limited) |
| Micronised Barley (Biopower Technology Limited) |
| 0.7-2.0 mm Crushed Corncob (Ebay, Craft and Design UK) |
| UK paper and card pulp (Fiberight Ltd.) |
| Palm oil (Supelco, 70905)) |
| 35%/65% Cotton-Polyester blend fabric (Leon's Fabric Store, Manchester) |
| ***Enzymes*** |
| Lipase B from *Candida antartica* (Sigma Aldrich, L3170; 3.1.1.3) |
| Proteinase K from *Tritirachium album* (Sigma Aldrich, 1.24568; 3.4.21.64) |
| Cellulase Blend from *Trichoderma reesei* (Sigma Aldrich, C2730; 3.2.1.4) |
| α Amylase from *Bacillus sp.* (Megazyme, E-BSTAA; 3.2.1.1) |
| Maltase from *Bacillus stearothermophilus* (Sigma Aldrich, G3651; 3.2.1.20) |
| CE1 Ferulic Acid Esterase from *Clostridia thermocellum* (Prozomix, PRO-E0355; 3.1.1.73) |
| ***Chemicals*** |
| ≥98% L-(+)-Lactic Acid (Sigma Aldrich, L1750; 79-33-4) |
| ≥99% Disodium Terephthalic Acid (Thermo Scientific, AA4294606; 10028-70-3) |
| 99.8% Ethylene Glycol (Sigma Aldrich, 102466; 107-21-1) |
| ≥99% D-(+)-Glucose (Sigma Aldrich, G8270; 50-99-7) |
| ≥99% D-(+)-Xylose (Sigma Aldrich, X1500; 58-86-6) |
| ≥99.5% Glycerol (Sigma Aldrich; G7893; 56-81-5) |
| 99% Pentadecanoic Acid (C15:0) (Sigma Aldrich, W433400, 1002-84-2) |
| 99% Lauric Acid (C12:0) (Fluorochem, 0988488; 143-07-7) |
| ≥98.5% Myristic Acid (C14:0) (Sigma Aldrich, W276413; 544-63-8) |
| 99% Pentadecanoic Acid (C15:0) (Sigma Aldrich, W433400, 1002-84-2) |
| 99% Palmitic Acid (C16:0) (Fluorochem, 995213; 57-10-3) |
| ≥95% Stearic Acid (C18:0) (Sigma Aldrich, W303518; 57-11-4) |
| 99% Oleic Acid (C18:1) (Fluorochem, 492471; 112-80-1) |
| >97% Linoleic Acid (C18:2) (Tokyo Chemical Industry Ltd, L0124; 60-33-3) |
| ≥99% *trans*-Ferulic Acid (Sigma Aldrich, W518301; 537-98-4) |
| ≥98% *p*-Coumaric Acid (Sigma Aldrich, C9008; 501-98-4) |
| ≥97% Vanillic Acid (Sigma Aldrich, H36001; 121-34-6) |
| 99% 4-Hydroxybenzoic Acid (Sigma Aldrich, H20059; 99-96-7) |
| 97% Protocatechuic Acid (Sigma Aldrich, E24859; 3943-89-3) |
| 3-Hydroxyhexanoic Acid (Toronto Research Chemicals, H825820; 10191-24-9) |
| 3-Hydroxyoctanoic Acid (Toronto Research Chemicals, S357998; 33796-86-0) |
| 3-Hydroxydecanoic Acid (Toronto Research Chemicals, H943233; 14292-26-3) |
| 99% 3-Hydroxydodecanoic Acid (Sigma Aldrich, H3398; 1883-13-2) |
| 3-Hydroxytetradecanoic Acid (Toronto Research Chemicals, H956780; 1961-72-4) |
| 98% 3-Hydroxyhexadecanoic Acid (Sigma Aldrich, H4398; 2389-34-7) |
| ≥99.5% Benzoic Acid (Sigma Aldrich, 242381; 65-85-0) |
| 3M Methanolic HCl (Supelco, 90964; 7647-01-1) |
| 99% Benzoyl Chloride (Sigma Aldrich, 259950; 98-88-4) |
| >98% Ethylene Glycol Dibenzoate (Tokyo Chemical Industry Ltd., E0300; 94-49-5) |
| Anti-interferon (IFN) alpha 2 antibody (Abcam, AB193055) |
| ≥95% SDS-PAGE Recombinant human IFN alpha protein (Abcam, AB285612) |

**Table S6.** Proportion of each waste component within the 25% (w/w) mixed waste hydrolysis for both high- and low-income waste.

| **Category** | **Component** | **Mixed Waste Composition (%)** | |
|---|---|---|---|
| | | **High Income** | **Low Income** |
| **Plastic** | PET film | 12.6 | 6.3 |
| | PLA beads | 5.4 | 2.7 |
| **Organic** | Wheat straw | 21.5 | 38 |
| **Paper and Board** | Municipal paper pulp | 34 | 9.5 |
| **Food** | Palm oil | 6.45 | 11.4 |
| | Corn cob | 10.75 | 19 |
| | Micronized barley | 2.6 | 4.6 |
| | Micronized apple | 0.9 | 1.5 |
| | Micronized beetroot | 0.9 | 1.5 |
| **Textiles** | Cotton-polyester blend | 5 | 5.5 |

**Table S7.** Treatment layout as dictated from design of experiments (DoE).

| | Substrate (in mM) | | | | | | | | | |
|---|---|---|---|---|---|---|---|---|---|---|
| **Treatment No.** | **Glucose** | **Xylose** | **TPA** | **EG** | **Glycerol** | **FFA** | **Coumaric** | **Ferulic** | **Lactic acid** | **Total** |
| **1** | 25 | 0 | 0 | 0 | 0 | 0 | 0 | 0 | 0 | 25 |
| **2** | 25 | 50 | 50 | 50 | 50 | 25 | 10 | 10 | 50 | 320 |
| **3** | 50 | 25 | 50 | 0 | 50 | 25 | 10 | 0 | 0 | 210 |
| **4** | 0 | 25 | 0 | 50 | 0 | 0 | 0 | 10 | 50 | 135 |
| **5** | 50 | 0 | 25 | 50 | 0 | 25 | 10 | 10 | 0 | 170 |
| **6** | 0 | 50 | 25 | 0 | 50 | 0 | 0 | 0 | 50 | 175 |
| **7** | 50 | 50 | 0 | 25 | 50 | 0 | 10 | 10 | 50 | 245 |
| **8** | 0 | 0 | 50 | 25 | 0 | 25 | 0 | 0 | 0 | 100 |
| **9** | 50 | 0 | 50 | 0 | 25 | 25 | 0 | 10 | 50 | 210 |
| **10** | 0 | 50 | 0 | 50 | 25 | 0 | 10 | 0 | 0 | 135 |
| **11** | 50 | 0 | 0 | 50 | 0 | 12.5 | 10 | 0 | 50 | 172.5 |
| **12** | 0 | 50 | 50 | 0 | 50 | 12.5 | 0 | 10 | 0 | 172.5 |
| **13** | 50 | 0 | 0 | 0 | 50 | 0 | 5 | 10 | 0 | 115 |
| **14** | 0 | 50 | 50 | 50 | 0 | 25 | 5 | 0 | 50 | 230 |
| **15** | 50 | 50 | 0 | 0 | 0 | 25 | 0 | 5 | 50 | 180 |
| **16** | 0 | 0 | 50 | 50 | 50 | 0 | 10 | 5 | 0 | 165 |
| **17** | 50 | 50 | 50 | 0 | 0 | 0 | 10 | 0 | 25 | 185 |
| **18** | 0 | 0 | 0 | 50 | 50 | 25 | 0 | 10 | 25 | 160 |
| **19** | 50 | 50 | 50 | 50 | 0 | 0 | 0 | 10 | 0 | 210 |
| **20** | 0 | 0 | 0 | 0 | 50 | 25 | 10 | 0 | 50 | 135 |
| **21** | 50 | 0 | 50 | 50 | 50 | 0 | 0 | 0 | 50 | 250 |
| **22** | 0 | 50 | 0 | 0 | 0 | 25 | 10 | 10 | 0 | 95 |
| **23** | 50 | 50 | 0 | 50 | 50 | 25 | 0 | 0 | 0 | 225 |
| **24** | 0 | 0 | 50 | 0 | 0 | 0 | 10 | 10 | 50 | 120 |
| **25** | 25 | 25 | 25 | 25 | 25 | 12.5 | 5 | 5 | 25 | 172.5 |

**Table S8.** Life cycle inventory data used for LCA and OPEX for LCC evaluations for the enzymatic hydrolysis of municipal solid waste (MSW) stage.

| Flows | Steps | Units | Amount | | | Cost (USD) | | |
|---|---|---|---|---|---|---|---|---|
| | | | High-income composition | Low-income composition | Source | High-income composition | Low-income composition | Source |
| **Inputs** | | | | | | | | |
| MSW | Hydrolysis | kg | 1000 | 1000 | Own experiments | | | |
| Enzymes | Hydrolysis | kg | 14.3 | 11.3 | Own experiments | 45.81 | 36.26 | Calculated based on [ref] |
| Buffer solution[b] | Hydrolysis | kg | 3106 | 3101 | Own experiments | 27.47 | 25.24 | Calculated based on [ref]for water and [ref] for sodium phosphate |
| Heat | Hydrolysis | MJ | 412 | 411 | Calculated based on [1] | 6.78 | 6.77 | Calculated based on [ref] |
| Electricity | Particle size reduction | MWh | 0.02 | 0.02 | | 4.3 | 4.3 | Calculated based on [ref] |
| Electricity | Filtration | MWh | 0.004 | 0.004 | Calculated based on [1] | 1.08 | 1.08 | Calculated based on [ref] |
| **Outputs** | | | | | | | | |
| Hydrolysate | Filtration | kg | 1728 | 1564 | Own experiments | | | |
| Waste[c] | Filtration | kg | 2392 | 2548 | Own calculations based on mass balances | 0.53[d] | 0.28[d] | Calculated based on [ref] |

Hydrolytic efficiency is 4% and 3.7% for high- and low-income compositions, respectively.

[b] The buffer solution is an aqueous solution of sodium phosphate, with an 2.4 wt% concentration for both high- and low-income compositions.

[c] Waste stream composition assumed at 80% solids and 20% liquid streams for high-income countries, and 90% solids and 10% liquid streams for low-income countries compositions.

[d] Only cost for wastewater treatment is considered as solid waste is assumed to be treated on-site via incineration.

**Table S9.** Life cycle inventory data used for LCA and OPEX for LCC evaluations for the microbial fermentation and PHA bioproduction stage.

| Flows | Steps | Units | Amount | | | Cost (USD) | | |
|---|---|---|---|---|---|---|---|---|
| | | | High-income composition | Low-income composition | Source | High-income composition | Low-income composition | Source |
| **Inputs** | | | | | | | | |
| Hydrolysate | Incubation | kg | 1728 | 1564 | Own experiments | | | |
| Sodium phosphate ($Na_2HPO_4$) | Incubation | kg | 11.8 | 10.7 | Own experiments | 0.005 | 0.004 | Calculated based on [ref] |
| Potassium phosphate ($KH_2PO_4$)[a] | Incubation | kg | 5.2 | 4.7 | Own experiments | 15 | 13.6 | Calculated based on [ref] |
| Sodium chloride (NaCl) | Incubation | kg | 0.9 | 0.8 | Own experiments | 2.84 | 2.57 | Calculated based on [ref] |
| Ammonium chloride ($NH_4Cl$) | Incubation | kg | 1.7 | 1.6 | Own experiments | 3.54 | 3.20 | Calculated based on [ref] |
| Heat | Incubation | MJ | 8.8 | 8.0 | Calculated based on [2] | 0.15 | 0.13 | Calculated based on [ref] |
| Electricity | Centrifugation | kWh | 7.5 | 6.8 | Calculated based on [2] | 2.02 | 1.83 | Calculated based on [ref] |
| **Outputs** | | | | | | | | |
| Wet cells | | kg | 26 | 26 | Own experiments | | | |
| Wastewater | | kg | 1722 | 1556 | Own calculations based on mass balances | 1.9 | 1.7 | Calculated based on [ref] |

Hydrolytic efficiency of base case is 4% and 3.7% for high- and low-income compositions, respectively.

[a] Potassium sulfate was used as proxy in the absence of data in commercial databases [3]

**Table S10.** Life cycle inventory data used for LCA, OPEX and revenues for LCC evaluations for the PHA extraction stage[a].

| **Flows** | **Steps** | **Units** | **Amount** | | | **Cost (USD)** | | |
|---|---|---|---|---|---|---|---|---|
| | | | **High-income composition** | **Low-income composition** | **Source** | **High-income composition** | **Low-income composition** | **Source** |
| **Inputs** | | | | | | | | |
| Wet cell | Chemical extraction | kg | 26 | 26 | Own experiments | | | |
| Sodium hydroxide (NaOH) | Chemical extraction | kg | 0.5 | 0.5 | Calculated based on [2] | 1.32 | 1.47 | Calculated based on [ref] |
| Sodium hypochlorite (NaOCl) | Chemical extraction | kg | 0.4 | 0.4 | Calculated based on [2] | 1.32 | 1.47 | Calculated based on [ref] |
| Hydrogen peroxide ($H_2O_2$) | Chemical extraction | kg | 0.5 | 0.5 | Calculated based on [2] | 1.95 | 2.18 | Calculated based on [ref] |
| Water | Chemical extraction | kg | 47 | 53 | Calculated based on [2] | 0.1 | 0.05 | Calculated based on [ref] |
| Heat | Chemical extraction | MJ | 0.08 | 0.10 | Calculated based on [2] | 0.001 | 0.002 | Calculated based on [ref] |
| | Dehydration | MJ | 0.06 | 0.06 | Calculated based on [2] | 0.001 | 0.001 | Calculated based on [ref] |
| Electricity | Filtration | kWh | 2.3 | 2.6 | Calculated based on [2] | 0.62 | 0.69 | Calculated based on [ref] |
| **Outputs** | | | | | | | | |
| PHA[b] | | kg | 1.2 | 1.3 | Own experiments | -10.61 | -11.82 | Calculated based on [ref] |
| Wastewater | | kg | 70 | 75 | Own calculations based on mass balances | 0.08 | 0.08 | Calculated based on [ref] |

[a] PHA is assumed to be chemically extracted using sodium hydroxide and sodium hypochlorite. The mixture is then filtered to remove 95% moisture content, and dehydrated to obtain the final product at commercial grade.

[b]The system was credited with the production of polyhydroxyalkanoate (PHA) from corn starch [4]

**Table S11.** Bioprocessing equipment costs provided per 1 t of valorised MSW.

| Step | Equipment | High income composition (USD) | Low income composition (USD) |
|---|---|---|---|
| Enzymatic hydrolysis[a] | Storage tank | 0.09 | 0.09 |
| | Crusher | 0.27 | 0.27 |
| | Mixer | 0.1 | 0.10 |
| | Hydrolysis tank | 0.87 | 0.87 |
| | Filter press[e] | 0.06 | 0.06 |
| Microbial fermentation and PHA bioproduction[a] | Mixer | 0.02 | 0.02 |
| | Fermenter | 1.85 | 1.71 |
| | Centrifuge | 0.19 | 0.18 |
| PHA extraction[b] | Extraction reactor | 0.08 | 0.08 |
| | Filter press[e] | - | - |
| | Evaporator | 0.11 | 0.12 |
| Service facilities[c] | | 1.09 | 1.05 |
| Maintenance[d] | | 0.36 | 0.35 |

Plant lifetime is assumed to be 25 years

[a] Estimated based on [ref]

[b] Estimated based on [ref]

[c] Assumed to be 30% of fixed cost [ref]

[d] Assumed to be 10% of fixed cost [ref]

[e]Assumed to be used in both processes

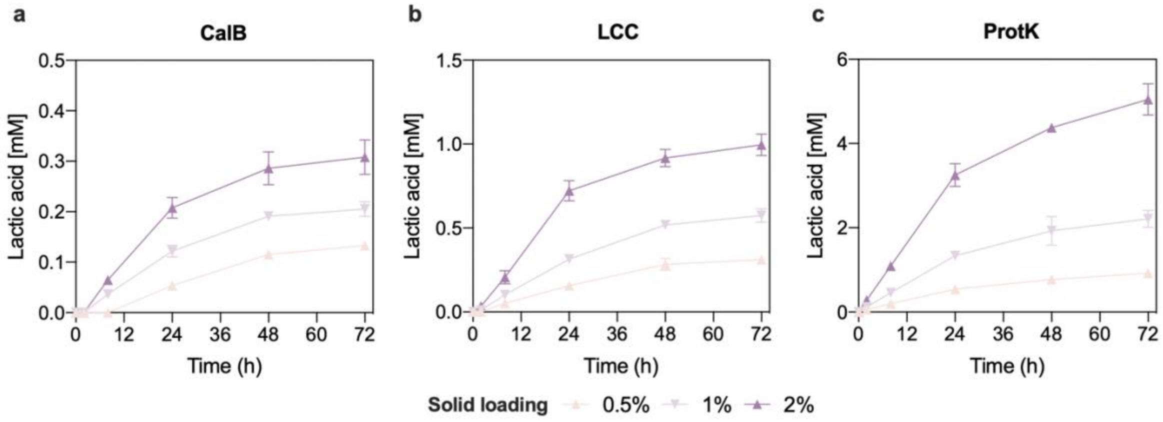


**Figure S1. Enzymatic hydrolysis of PLA plastic beads.** Hydrolysis of 0.5-2% PLA beads (DW) by either 9 U CAL-B/g PLA (**a**), 0.75 mg LCC/g PLA (**b**) or 0.5 mg proteinase K/g PLA (**c**) at 50 °C and pH 7.0 for 72 hours. Released lactic acid is plotted against time with all data points being mean ± SD of n = 3 biological replicates. CalB, Candida Antarctica Lipase B; LCC, Leaf-branch compost cutinase; ProtK, Proteinase K.

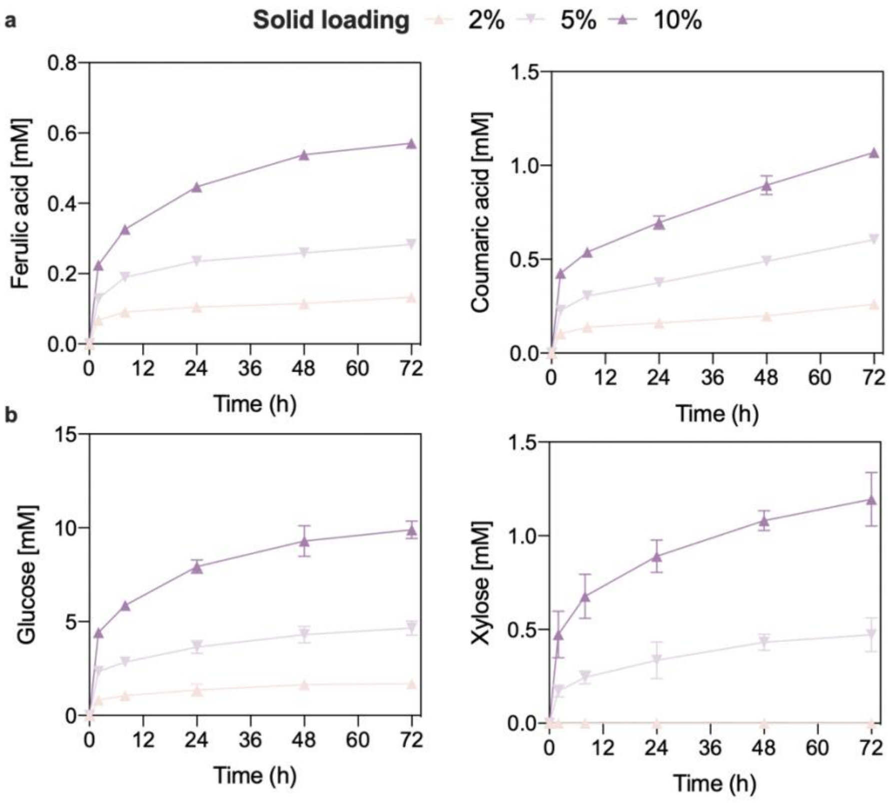


**Figure S2. Enzymatic hydrolysis of milled corncobs.** Hydrolysis of 2-10% milled corncob (DW) by 1 mg CE1/g corncob (**a**) or 50 mg CTEC-2/g corncob (**b**). Both hydrolyses were performed at 50 °C and pH 7.0 for 72 hours. Released ferulic acid and coumaric acid (**a**) and glucose and xylose (**b**) are plotted against time with all data points being mean ± SD of n = 3 biological replicates.

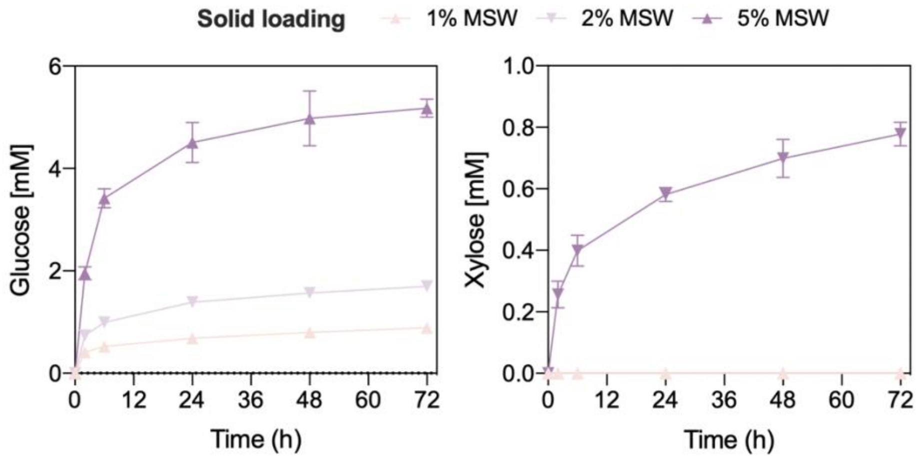


**Figure S3. Enzymatic hydrolysis of paper and cardboard pulp.** Hydrolysis of 1-5% MSW paper pulp (DW) by 50 mg CTEC-2/g paper pulp at 50 °C and pH 7.0 for 72 hours. Released glucose and xylose are plotted against time with all data points being mean ± SD of n = 3 biological replicates.

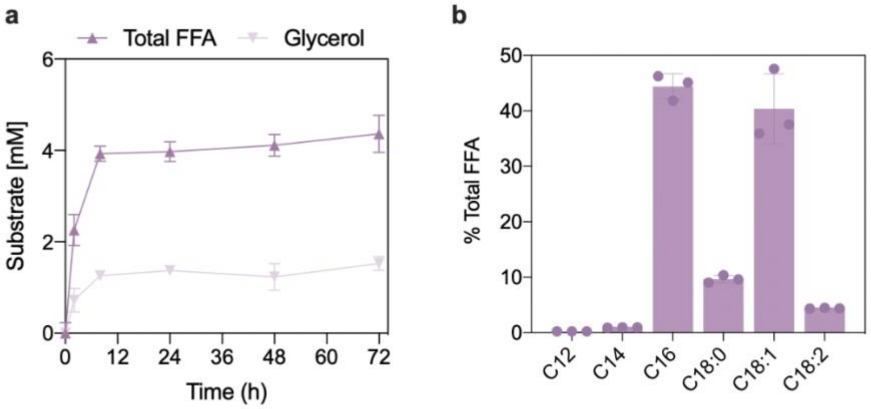


**Figure S4. Enzymatic hydrolysis of palm oil.** Hydrolysis of 1% (DW) palm oil by 9 U CAL-B/g palm oil at 50 °C and pH 7.0 for 72 hours. Released total free fatty acids ($C_{12:0-18:0}$) and glycerol are plotted against time (**a**), the ratio of fatty acid to glycerol was approximately 3:1. Proportion of the individual fatty acid constituents released from palm oil hydrolysis (**b**). All data points are mean ± SD of n = 3 biological replicates.

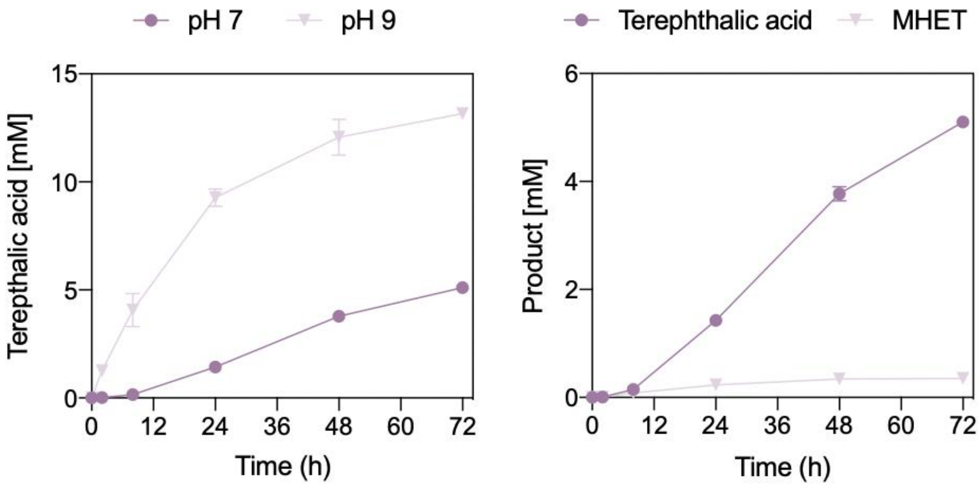


**Figure S5. Enzymatic hydrolysis of PET plastic film.** Hydrolysis of 4% PET plastic (DW) by 0.75 mg LCC/g PET at 50 °C and pH 7.0 or 70ºC and pH 9 (**a**) or at 50 °C pH 7.0 alone to see MHET release under these environmental conditions (**b**) for 72 hours. Released TPA and MHET are plotted against time with all data points being mean ± SD of n = 3 biological replicates.

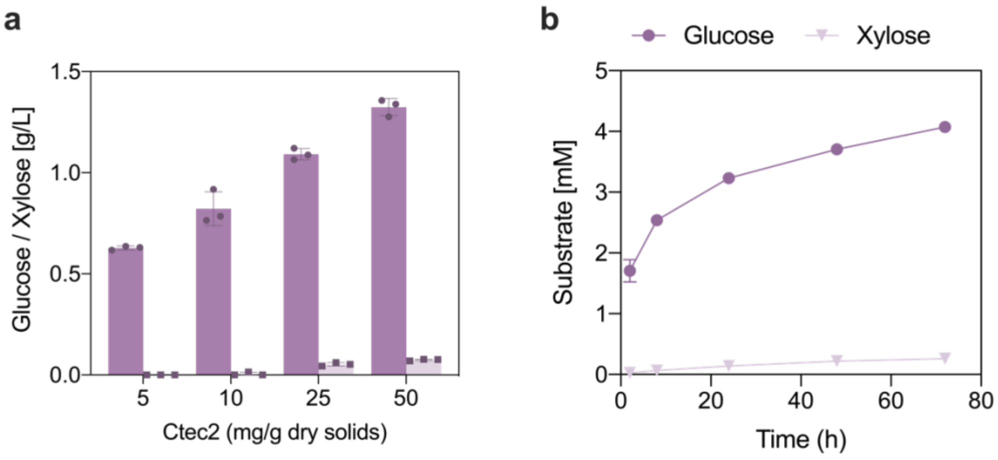


**Figure S6. Enzymatic hydrolysis of wheat straw.** Hydrolysis of 5% wheat straw (DW) by an enzyme loading ranging from 5-50 mg CTEC-2/g PET at 50 °C and pH 7.0 for 24 hours, released glucose and xylose are plotted (**a**). Time course of glucose and xylose release from the hydrolysis of 10% wheat straw (DW) by 50 mg CTEC-2/g wheat straw at 50 °C and pH 7.0 for 72 hours (**b**). All data points are mean ± SD of n = 3 biological replicates.

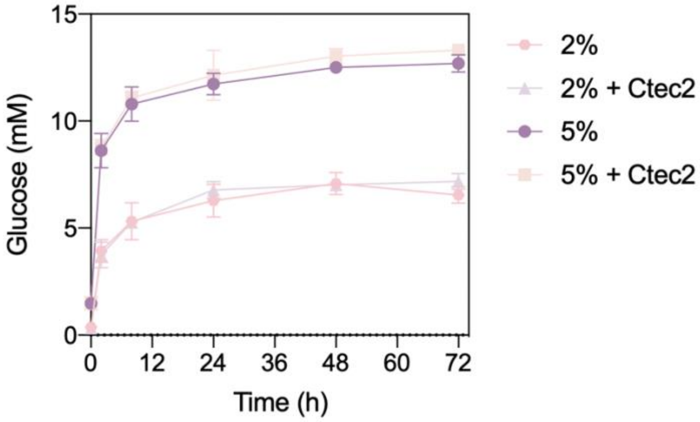


**Figure S7. Enzymatic hydrolysis of micronized food mixture.** Hydrolysis of 2% or 5% (DW) food waste mix (consisting of 2:1:1 barley, apple, and beetroot mix) by 10 U α-Amylase and 5 U maltase/g food ± 50 mg CTEC-2/g food at 50 °C and pH 7.0 for 72 hours. Released glucose is plotted against time with all data points being mean ± SD of n = 3 biological-replicates.

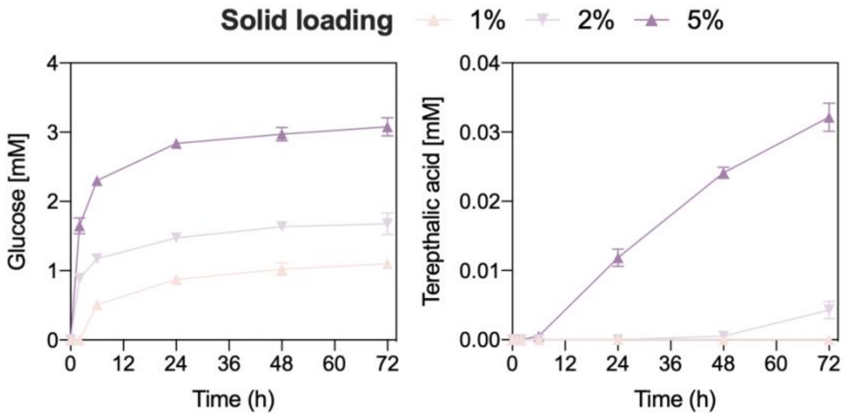


**Figure S8. Enzymatic hydrolysis of textile fabrics.** Hydrolysis of 1-5% (DW) cotton/polyester blend fabric (65:35) by and 50 mg CTEC-2/g fabric (**a**) or 0.75 mg LCC/g fabric (**b**) at 50 °C and pH 7.0 for 72 hours. Released glucose (**a**) and TPA (**b**) are plotted against time with all data points being mean ± SDs of n = 3 biological replicates.

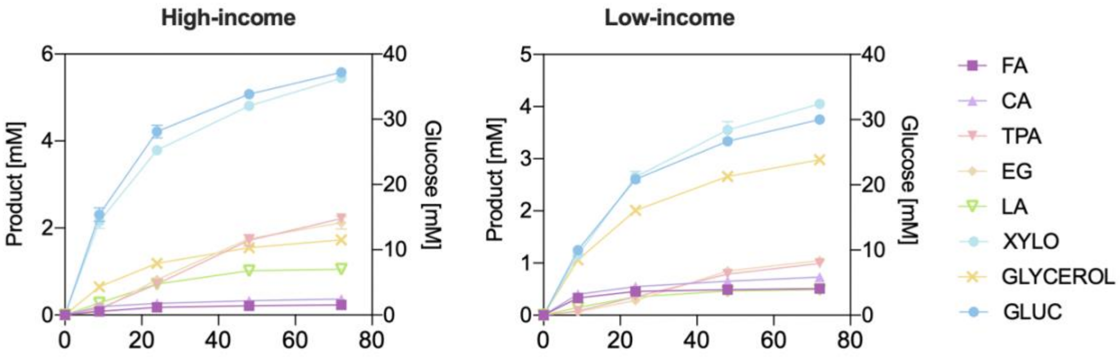


**Figure S9. Time-course of the enzymatic hydrolysis of MSW for high- and low-income representation.** Hydrolysis of 25% (DW) MSW for the high-income formula (**a**) and low-income formula (**b**) by 50 mg CTEC-2/g MSW, 0.75 mg LCC/g MSW, 10 U α-amylase/g MSW, 5 U maltase/g MSW, 1 mg CE1/g MSW, 0.75 mg LCC/g MSW and 0.5 mg proteinase K/g MSW at 50 °C and pH 7.0 for 72 hours. Released glucose (GLUC), xylose (XYLO), ferulic acid (FA), coumaric acid (CA), TPA, EG, lactic acid (LA) and glycerol are plotted against time with all data points being mean ± SD of n = 3 biological replicates.

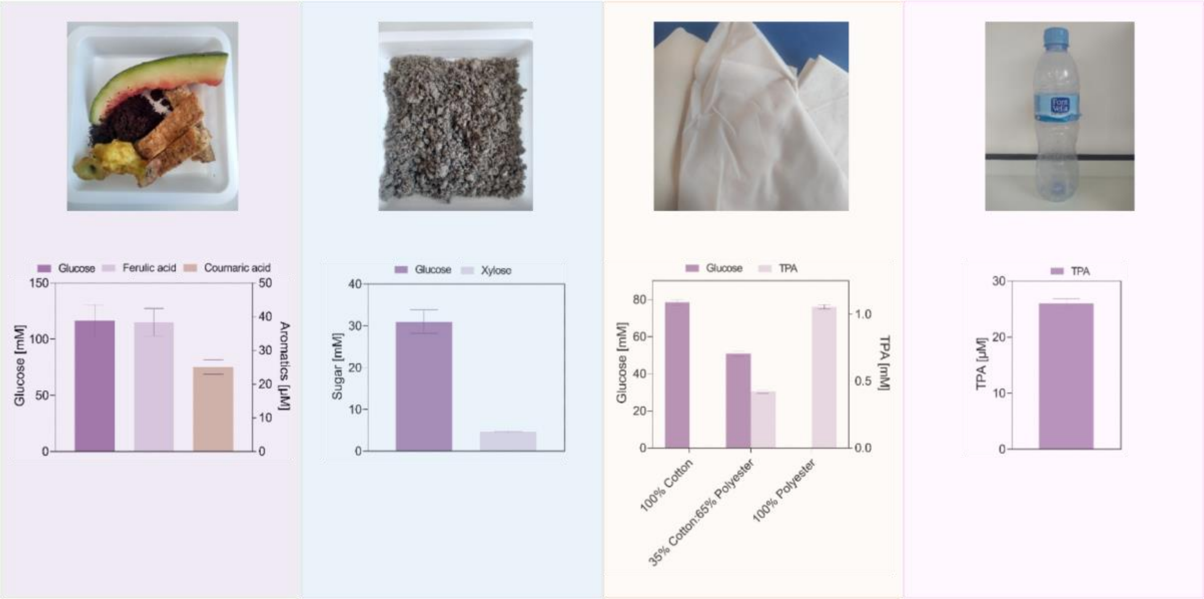


**Figure S10. Enzymatic hydrolysis of post-consumer waste.** (a) hydrolysis of 10% (DW) food waste (spent coffee, apple, watermelon rind, and bread crust at a ratio of 1:1:1:1) using 10 U/g α-amylase, 5 U/g maltase, 15 mg/g CTEC-2, and 1mg/g CE1. Hydrolysis was performed at pH 6 and 55 °C for 72 hours. (b) Hydrolysis of 10% (DW) post-consumer paper pulp using 15 mg/g CTEC-2. Hydrolysis was performed at pH 5 and 50 °C for 72 hours. (c) Hydrolysis of 100% cotton, 100% polyester or a 35% cotton:65% polyester cotton blended fabric using either 15 mg/g CTEC-2 at pH 5 and 50 °C or 0.75 mg/g LCC at pH 9 and 70 °C for 72 hours. (d) Hydrolysis of neck of a Font Vella PET plastic bottle using 0.75 mg/g LCC at pH 9 and 70 °C for 72 hours. All data points are mean $\pm$ SD of n = 3 biological replicates.

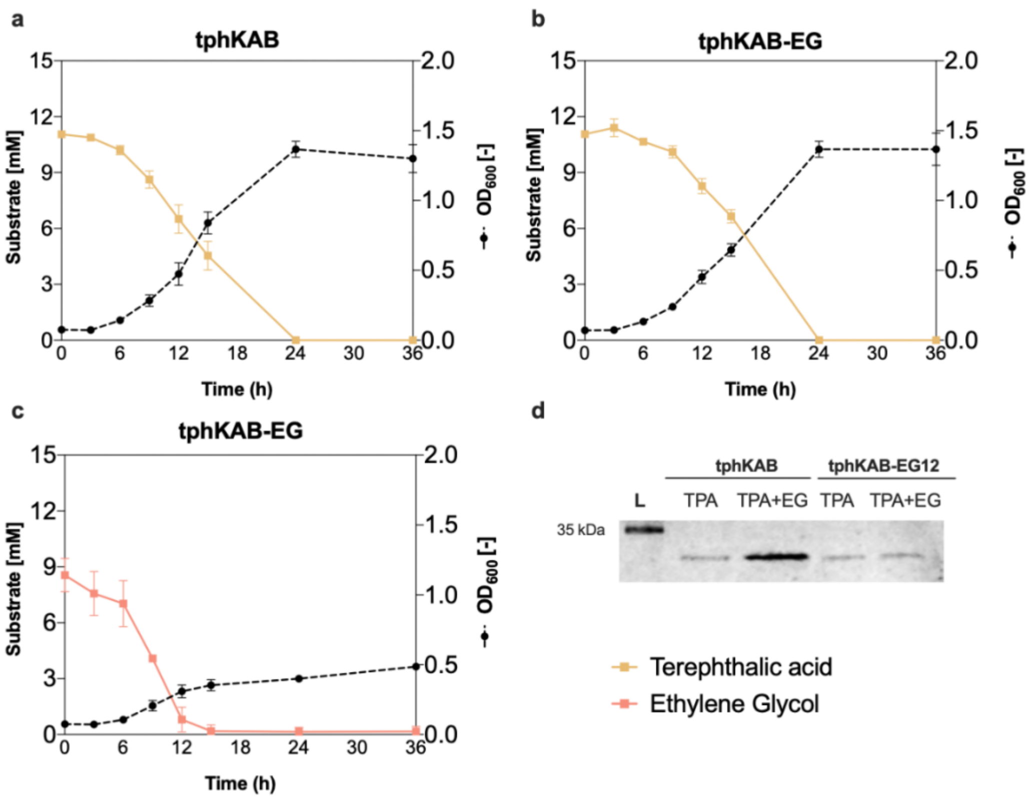


**Figure S11. Engineered catabolism of PET monomers and insulin production in *P. putida*. (a-c)** Growth and substrate consumption of TPA or EG as the sole carbon source by strain tphKAB (**a, b**) and tphKAB-EG (**c**) in shake flasks containing M9 media supplemented with 10 mM of the corresponding substrate. All data points are mean ± SD of n = 3 biological replicates. (**d**) Western blot of whole-cell samples of tphKAB or tphKAB-EG grown in shake-flasks containing either 20mM TPA or a 20 mM TPA:EG co-feed.

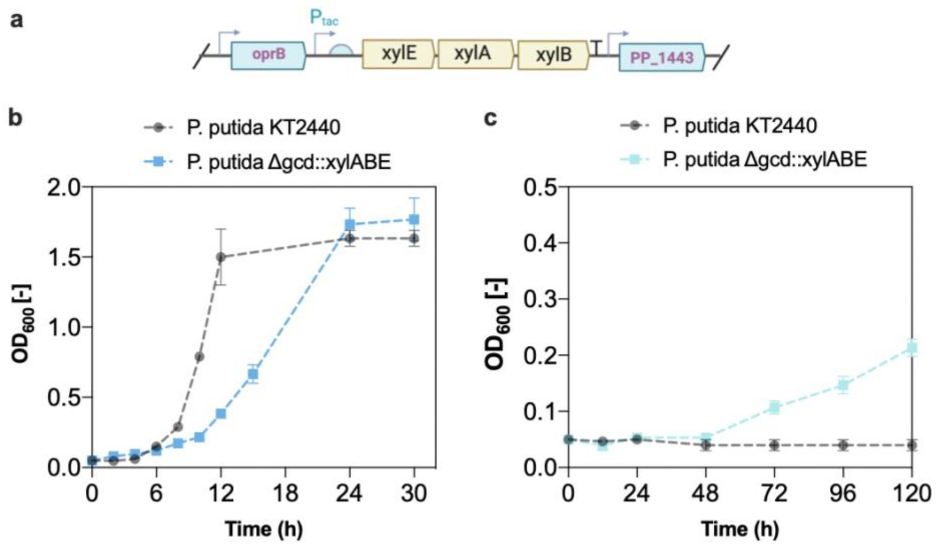


**Figure S12. Xylose catabolism in *P. putida.*** The XylAB catabolic operon from *E. coli* was integrated by replacement of glucose dehydrogenase (*gcd)* (**a**). The resulting strain, tphKAB-XylABE can grow in shake-flasks containing M9 media supplemented with 10 mM glucose (a) and xylose (b) as the sole carbon source. All data points are mean ± SD of n = 3 biological-replicates.

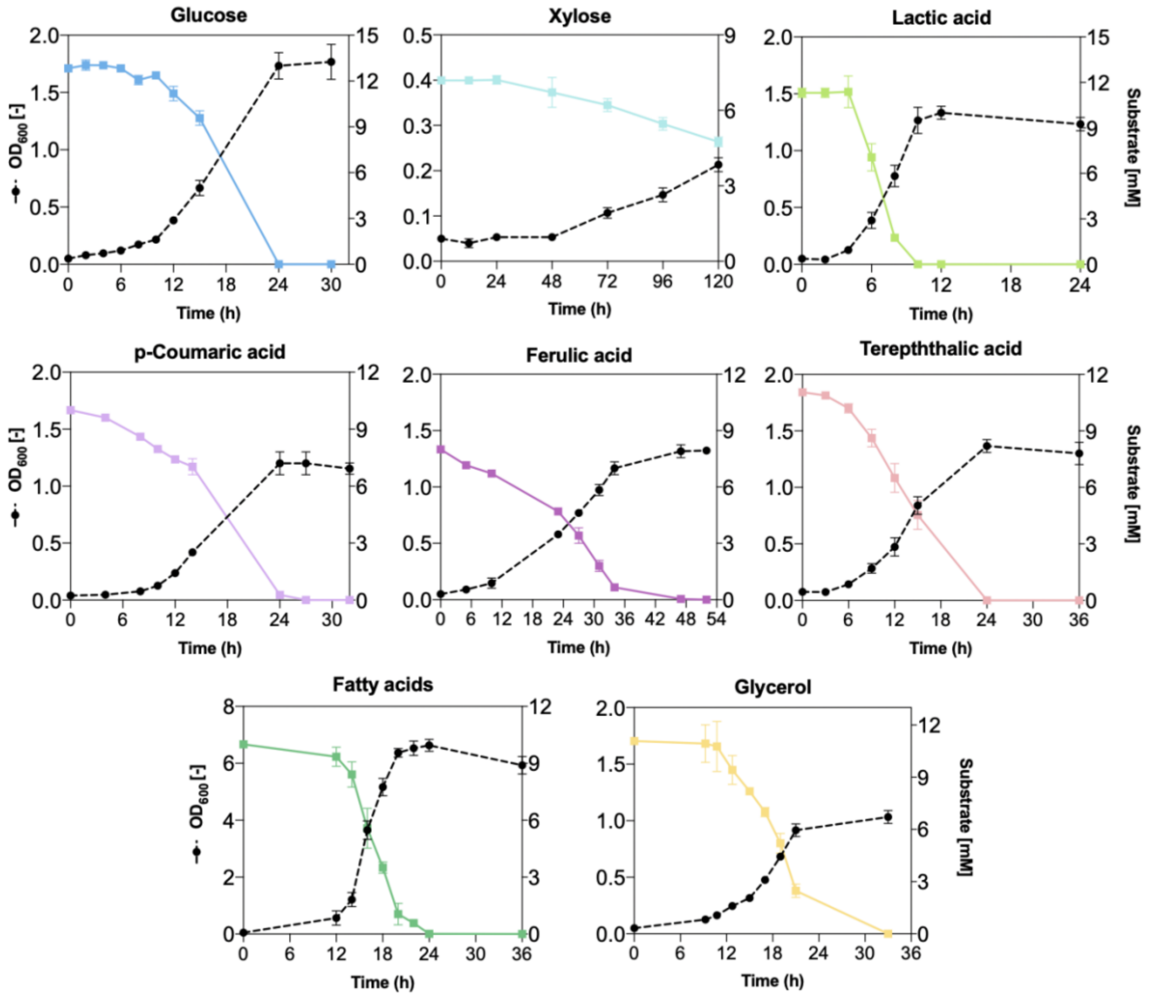


**Figure S13. Growth and consumption profiles of MSW-derived substrates by the engineered tphKAB-XylABE strain.** Strain tphKAB-XylABE was cultivated in shake-flasks containing M9 media supplemented with 10 mM of the corresponding MSW-derived substrate. All data points are mean ± SD of n = 3 biological replicates.

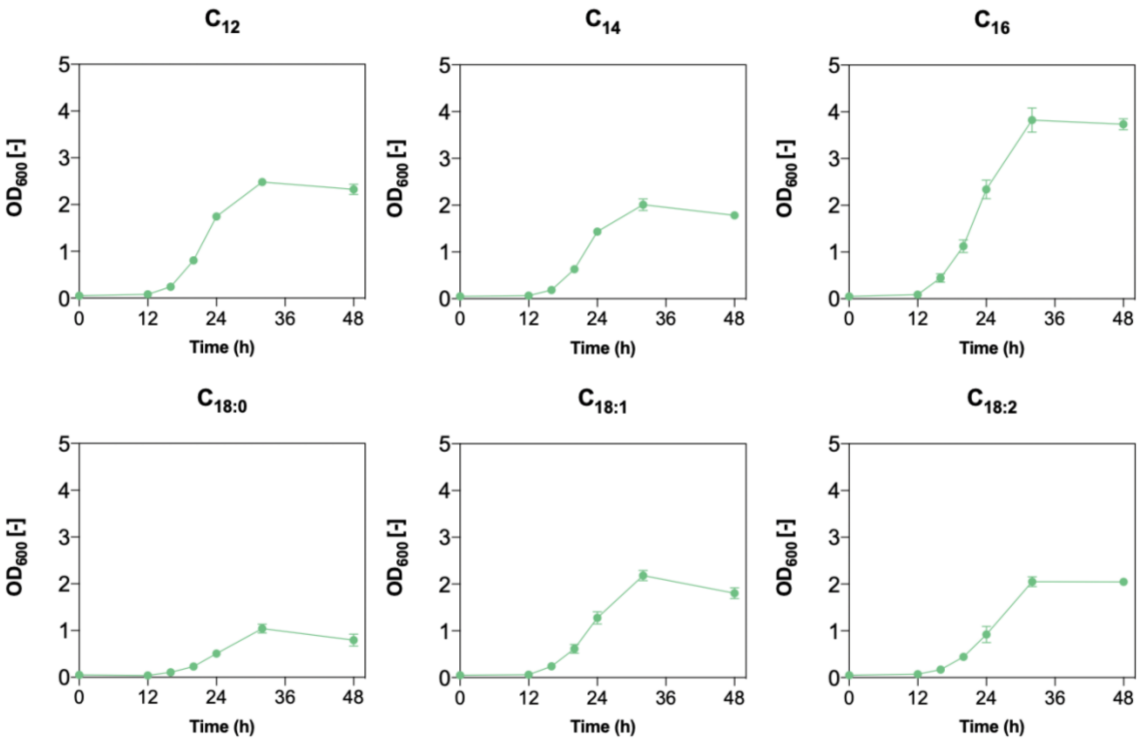


**Figure S14. Utilization of individual palm-oil representative fatty acids by *P. putida.*** Cultivation of tphKAB-XylABE in M9 media supplemented with 2.5 mM fatty acid of varying in chain length from $C_{12}$ through $C_{18}$. Cultivations were performed in shake-flasks, and all data points are mean ± SD of n = 3 biological replicates. $C_{12}$, lauric acid; $C_{14}$, myristic acid; $C_{16}$, palmitic acid; $C_{18:0}$, stearic acid; $C_{18:1}$, oleic acid; $C_{18:2}$, linoleic acid.

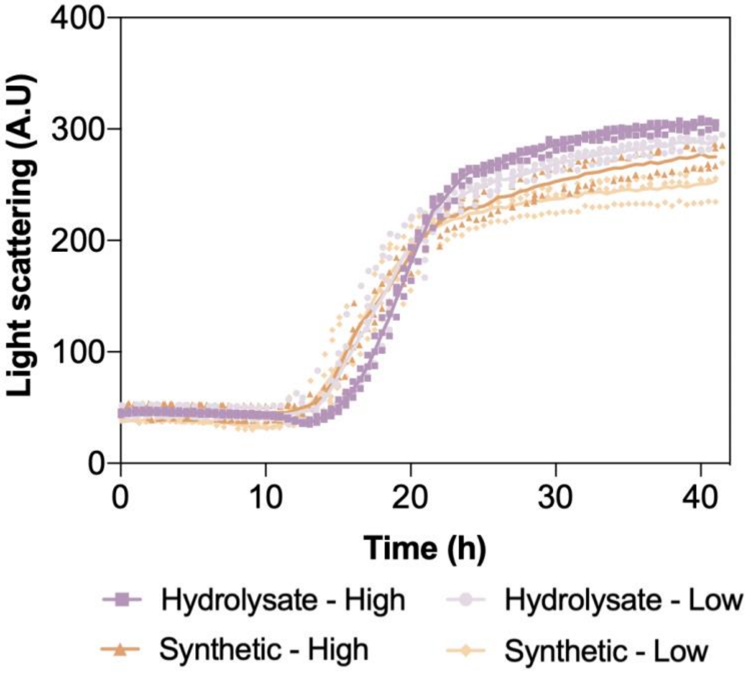


**Figure S15. Microbioreactor growth assays with tphKAB-XylABE.** The engineered strain was grown on both high- and low-income waste enzymatic hydrolysates (obtained from Fig. 2) and mock synthetic substrates.

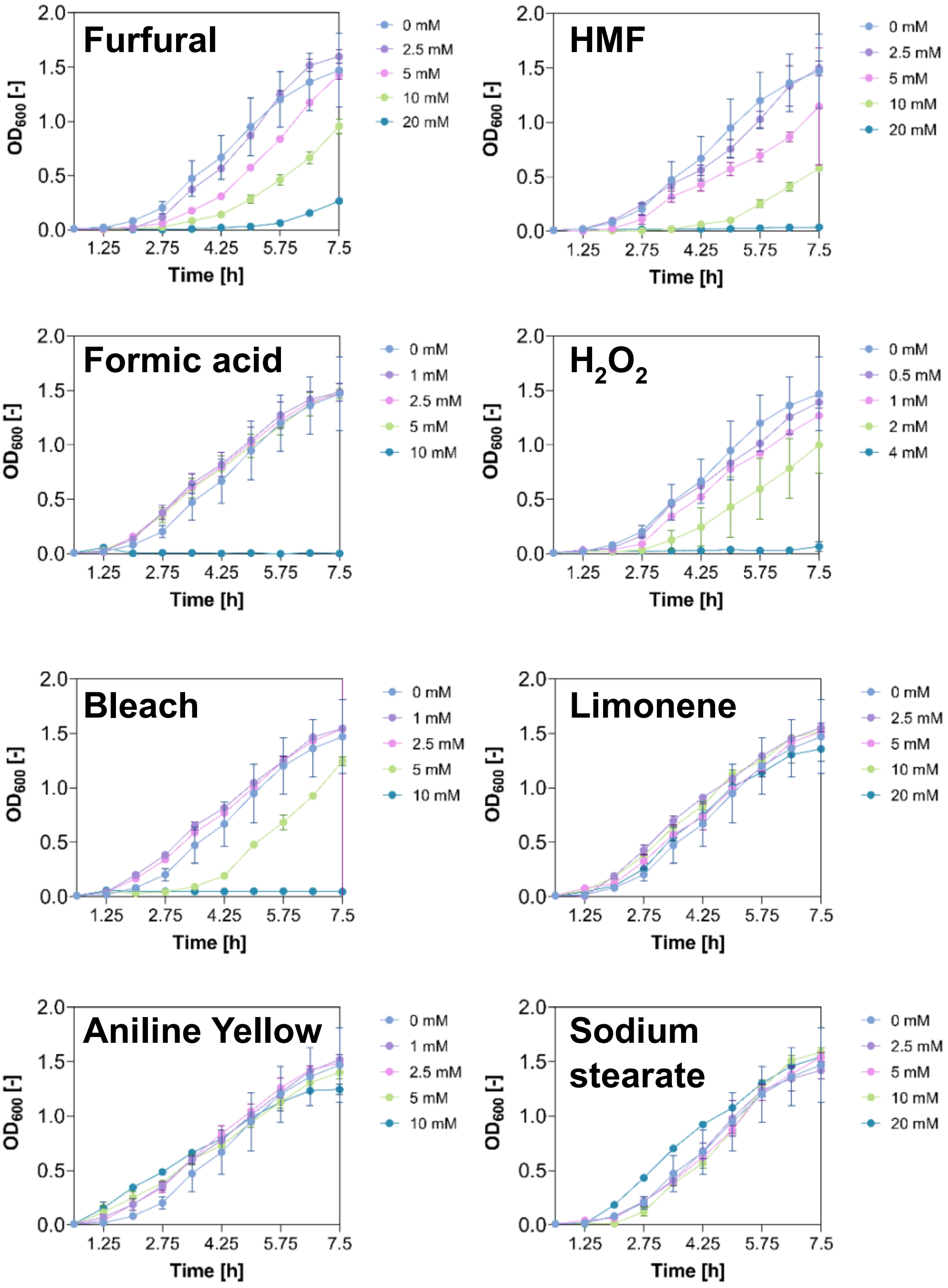


**Figure S16. Growth profile of wild type *P. putida* in the presence of potential MSW-derived inhibitors.** *P. putida* KT2440 was cultivated in LB media supplemented with variable concentrations of each inhibitor. All data points are mean + SD of n = 3 biological replicates.

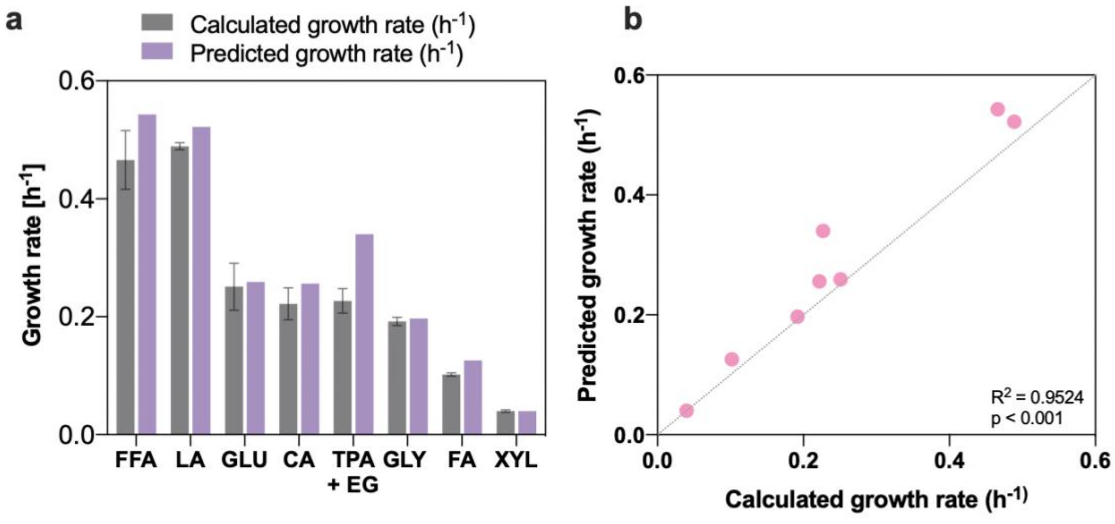


**Figure S17. Growth rate prediction by flux balance analysis (FBA) with a modified genome-scale metabolic model of *P. putida* (iJN1463).** FBA was performed with the tphKAB-XylABE model and constrained with consumption data obtained from each individual substrate cultivation, where growth was set as the objective function. The resulting growth rates (**a**) are accurately predicted by FBA (linear regression) (**b**). All experimental data points are mean ± SD of n = 3 biological replicates.

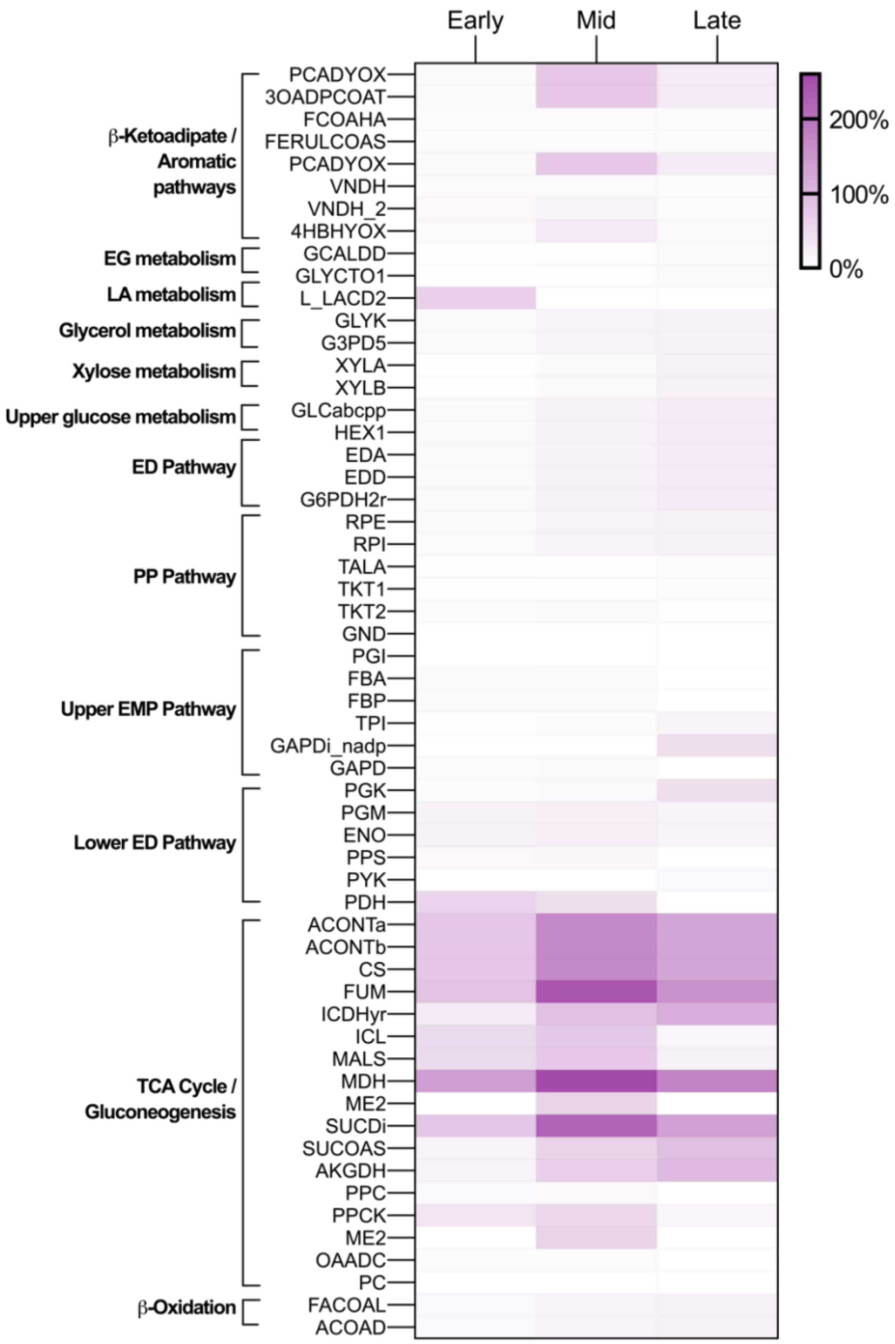


**Figure S18. Cellular flux distribution in an equimolar mixture of the nine MSW-derived feedstock.** Net predicted metabolic fluxes[5] (in mmol gCDW$^{-1}$ h$^{-1}$) by FBA from main central metabolic reactions normalized to the total uptake flux (%) during early, mid, and late-exponential phases in the equimolar MSW mixture. Model was constrained with uptake data obtained experimentally.

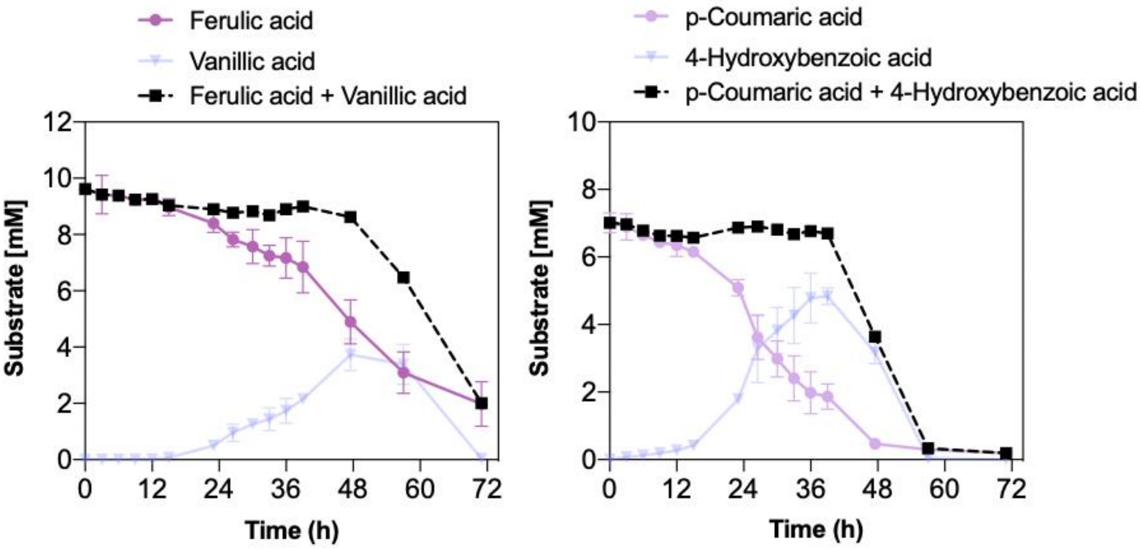


**Figure S19. Utilization of both hydroxycinnamic acids by engineered *P. putida*.** Accumulation and consumption of ferulic (left) and coumaric (right) acid and their pathway intermediates from the cultivation of tphKAB-XylABE in M9 media supplemented with all nine MSW-derived substrates. All data points are mean ± SD of n = 3 biological replicates.

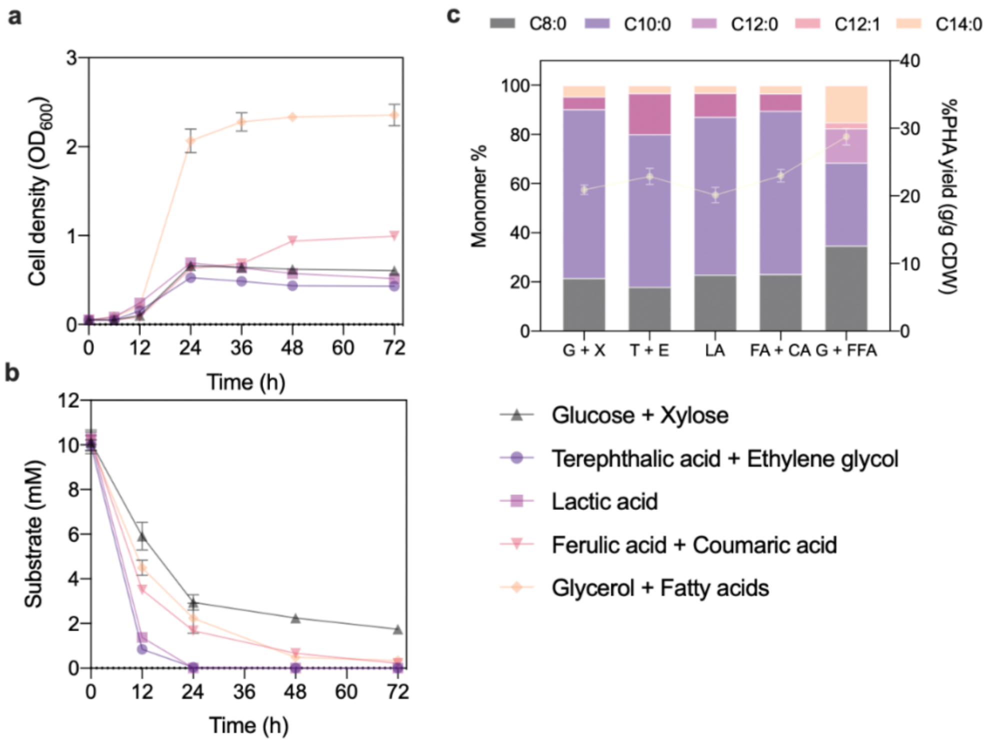


**Figure S20. Yield of PHA by MSW substrate type.** Biomass accumulation (**a**), substrate consumption (**b**) and PHA production (**c**) of tphKAB when grown in different substrate combinations. Cultivations were performed in M9 in shake-flasks with an initial substrate loading of 10 mM, and PHA accumulation/composition was determined at 72 hours. All data points are mean ± SD of n = 3 biological replicates.

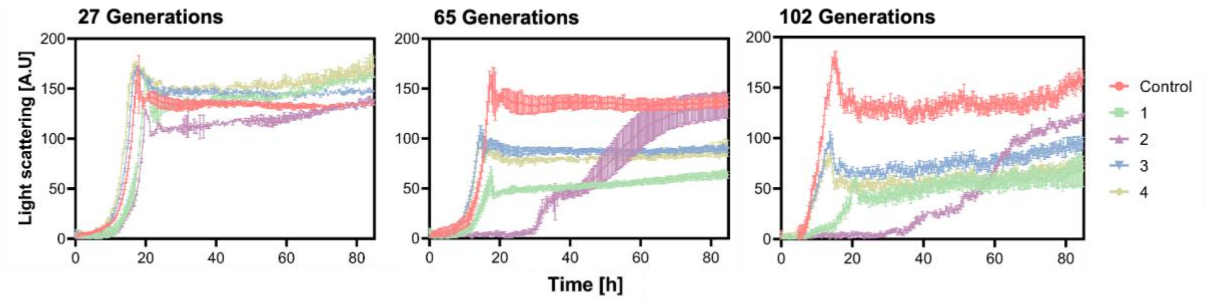


**Figure S21. Plasmid retention by tphKAB-XylABE passaged for a variable number of generations.** Strain tphKAB-XylABE was passaged for either 27, 65 or 102 generations in M9 supplemented with (1) 500 µg/ml carbenicillin + 10 mM glucose, (2) 10 mM glucose, (3) 5 mM glucose + 5 mM TPA, or (4) 500 µg/ml carbenicillin + 5 mM glucose + 5 mM TPA. Each passage condition was then evaluated for plasmid retention by evaluating strain growth in M9 + 10 mM TPA with a non-passaged tphKAB-XylABE strain as a control. All data points are mean + SD of n = 3 biological replicates.

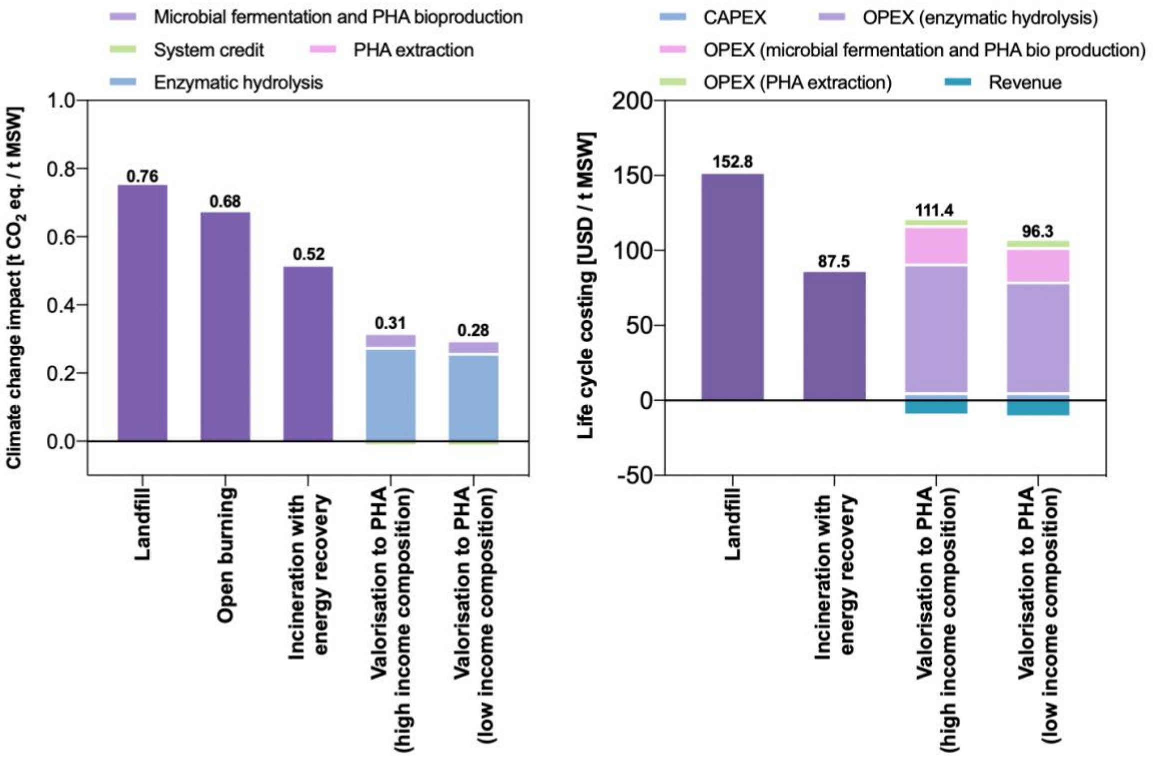


**Figure S21. Climate change and economic impact of the treatment and valorisation of 1 tonne of MSW.** Contribution analysis for the different stages included in the estimations of the total climate change impact (left) and Life cycle costing (right) for the proposed PHA valorisation process, and comparison with conventional technologies.

### Methods references

1 Moreno, J. *et al.* Life-cycle sustainability of biomass-derived sorbitol: Proposing technological alternatives for improving the environmental profile of a bio-refinery platform molecule. *Journal of Cleaner Production* **250**, 119568, doi:https://doi.org/10.1016/j.jclepro.2019.119568 (2020).

2 Andreasi Bassi, S., Boldrin, A., Frenna, G. & Astrup, T. F. An environmental and economic assessment of bioplastic from urban biowaste. The example of polyhydroxyalkanoate. *Bioresour Technol* **327**, 124813, doi:10.1016/j.biortech.2021.124813 (2021).

3 Francesco Longhini, O. P. D7.2 – Final report on Life-cycle assessment. (2021).

4 Zhong, Z. W., Song, B. & Huang, C. X. Environmental Impacts of Three Polyhydroxyalkanoate (PHA) Manufacturing Processes. *Materials and Manufacturing Processes* **24**, 519-523, doi:10.1080/10426910902740120 (2009).

5 Nogales, J. *et al.* High-quality genome-scale metabolic modelling of Pseudomonas putida highlights its broad metabolic capabilities. *Environ Microbiol* **22**, 255-269, doi:10.1111/1462-2920.14843 (2020).
